# Multi-ancestry conditional and joint analysis (Manc-COJO) applied to GWAS summary statistics

**DOI:** 10.64898/2026.01.30.702783

**Authors:** Xiaotong Wang, Yong Wang, Peter M Visscher, Naomi R Wray, Loic Yengo

## Abstract

Conditional and joint (COJO) analysis of genome-wide association study (GWAS) summary statistics to identify single nucleotide polymorphisms (SNPs) independently associated with a trait is standard in post-GWAS pipelines. GWAS meta-analyses are increasingly conducted across multiple ancestry groups but how to perform COJO in a multi-ancestry context is not known. Here we introduce Manc-COJO, a method for multi-ancestry COJO analysis. Simulations and real-data analyses show that Manc-COJO improves the detection of independent association signals and reduces false positives compared to COJO and *ad h*oc adaptations for multi-ancestry use. We also introduce Manc-COJO:MDISA, a follow-up within ancestry algorithm to identify ancestry-specific associations after fitting Manc-COJO identified SNPs. The C++ implementation of Manc-COJO substantially improves on computational efficiency (for single ancestry >120 times faster than GCTA-COJO software) and supports linkage disequilibrium references derived either from individual-level genotype data or pre-computed matrices, facilitating analysis when data sharing is limited.

## Introduction

GWAS have revolutionised understanding of the genetic architecture of complex traits^1^. Typically, the GWAS paradigm tests for association between a trait and SNPs, one at a time (giving marginal SNP effect estimates). However, associations detected in GWAS are often not independent because of linkage disequilibrium (LD) between SNPs. In principle, multivariate linear regression analysis can be applied to GWAS data to identify secondary SNP associations, which are SNP associations detected after conditioning on one or more significantly associated SNPs. However, such analyses are computationally demanding and are not possible in modern GWAS which are typically meta-analyses of summary statistics from contributing studies, without sharing of individual level data. To address this challenge, the conditional and joint (COJO) multiple-SNP analysis of GWAS summary statistics was developed and implemented into the GCTA software^2^. The algorithm requires an LD matrix to model the correlation structure between SNPs which the software generates from user-provided individual-level genotype data. The GCTA-COJO algorithm was developed with an implicit assumption that the GWAS was conducted in a single ancestry, and is hereafter called Sanc-COJO (**S**ingle-**anc**estry **co**nditional and **jo**int analysis).

GWAS are now commonly conducted as meta-analyses of GWAS across samples with diverse ancestries^3-5^. Ancestry groups differ in allele frequencies and LD patterns because of demographic and evolutionary processes such as random drift^6^. A key question when implementing post GWAS analyses such as COJO is how to model the correlation structure between SNPs in a multi-ancestry GWAS setting. Some studies have repurposed Sanc-COJO to analyse multi-ancestry GWAS data either by using LD matrices derived from European ancestry cohorts^5,7^, as these dominate (typically >80%) current GWAS meta-analyses^8^, or by constructing LD matrices from reference samples that match the ancestry composition in the meta-analysis^9,10^. Other approaches include conducting COJO analysis separately within each ancestry^11-13^ or performing COJO analysis in one ancestry and attempting replication in other ancestries^14,15^. However, to our knowledge these *ad hoc* approaches have not been formally evaluated, and optimal strategies require further consideration. Meanwhile, although Sanc-COJO was not originally designed as a fine-mapping tool, its ease of use has led many researchers to incorporate COJO into their fine-mapping pipeline^16,17^. However, interpretation of COJO-based independent associations as being causal merits further investigation.

In this study, we introduce **m**ulti-**anc**estry **co**nditional and **jo**int analysis (Manc-COJO), which extends the capabilities of stepwise conditional and joint association analysis to multi-ancestry datasets. Through extensive simulations and real data applications, we demonstrate that Manc-COJO improves the detection of independent trait-associated loci and reduces false positive discoveries. An underlying assumption of our method (supported by an increasing body of evidence) is that causal variants are largely shared across ancestries and have similar effect sizes across ancestries^18-21^. However, we demonstrate that Manc-COJO is robust to violations of this assumption. To address scenarios involving ancestry-specific causal variants^22,23^, we propose a follow-up analysis: multi-ancestry data-informed single ancestry analysis (Manc-COJO:MDISA), that applies a COJO analysis within a single ancestry while accounting for SNPs identified by Manc-COJO.

## Results

### Overview of Manc-COJO

Manc-COJO is a forward and backward stepwise conditional and joint analysis method that extends the current Sanc-COJO algorithm into multi-ancestry settings. Illustrated in **Figure 1** for two ancestries, Manc-COJO utilises GWAS summary statistics and LD reference matrices from each ancestry as input. Manc-COJO is equivalent (when an in-sample LD matrix is used) to stepwise forward and backward variable selection algorithm applied to individual-level genotypic data. Briefly, in each iteration, the top-associated SNPs are regressed out from the original phenotype within each ancestry, and the regression residuals serve as the new phenotype for a subsequent GWAS within each ancestry (conditional analysis, conditioned on the top-associated SNPs). A new SNP is then selected for inclusion in the model based on p-values from the meta-analysis of GWAS of each individual ancestry (forward selection). Following selection of a new SNP, all previously chosen SNPs, together with the newly selected SNP, are fitted jointly in a multiple linear regression model using the original phenotype within each ancestry (joint analysis). The newly selected SNP is retained as a COJO-SNP based on the squared z-value in the meta-analysis of joint-effects and adjusted R^2^ (coefficient of determination; an optional criterion) of the joint model within each ancestry. If its inclusion renders previously included SNPs insignificant in the meta-analysis of joint effects, those insignificant SNPs are removed, and the process continues (backward selection). This iterative process repeats until no further SNPs can be added or excluded from the model. Notably, Manc-COJO software has improved computational efficiency over the Sanc-COJO implementation in GCTA and has been developed with an option to take LD matrices as reference input (as opposed to individual level data to build the LD matrices).

**Figure 1.**
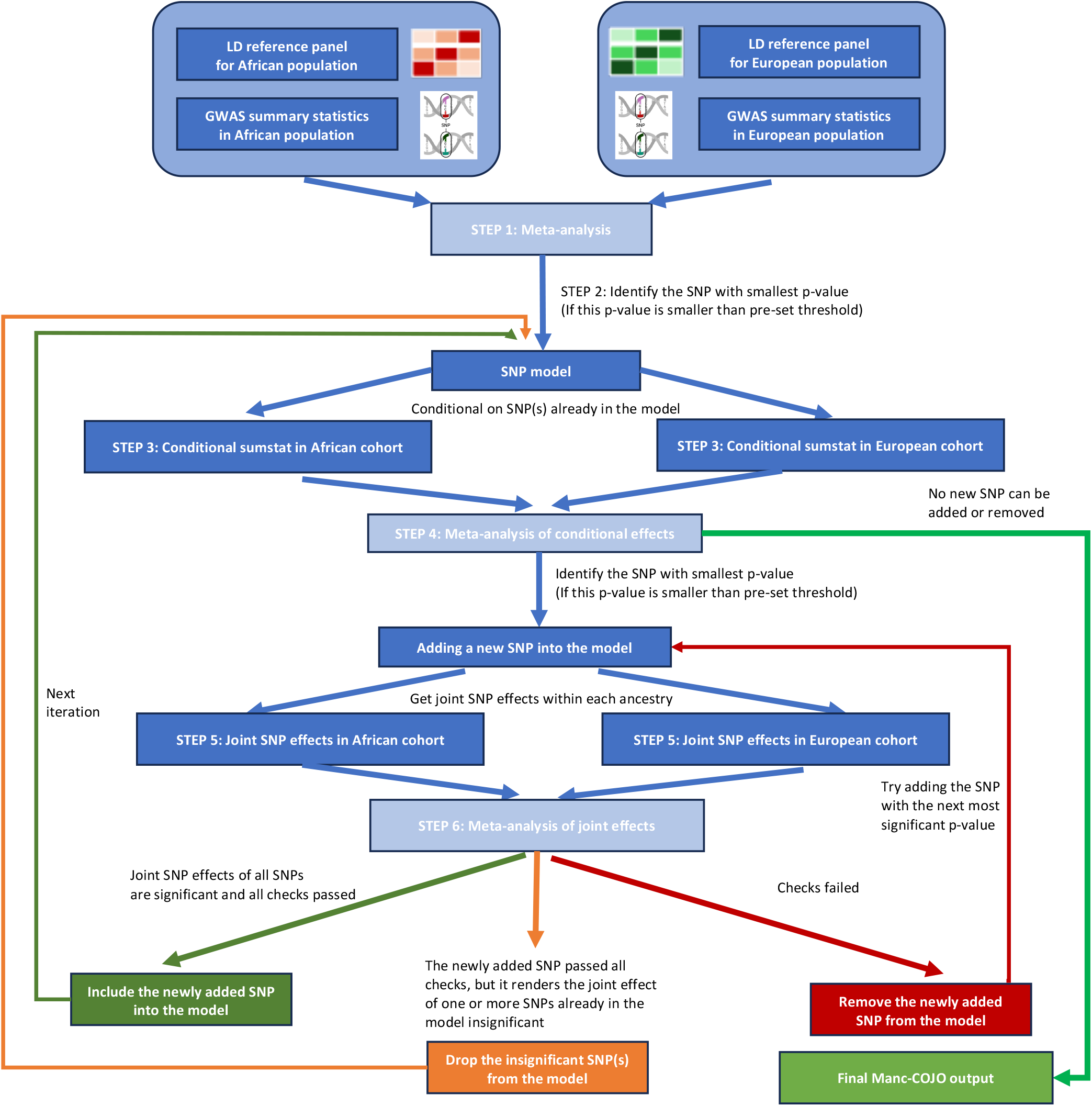
A schematic overview of the Manc-COJO algorithm. To facilitate the integration of multi-ancestry data, Manc-COJO is a a forward and backward stepwise conditional and joint analysis method that extends the current Sanc-COJO algorithm into trans-ancestry settings. Manc-COJO utilises GWAS summary statistics and LD references from different ancestries as input (illustrated here with two ancestries), enabling the use of LD matrices as reference inputs to support the global sharing of accurate LD structures.

### Simulation study results

We first compared the model output (hereafter referred to as “COJO SNPs”) from three model applications: 1) Manc-COJO applied to GWAS summary statistics from two ancestries (European and African) each of sample size N=6,901, 2) Sanc-COJO applied to the same meta-analysed GWAS summary statistics, denoted as Sanc-COJO->Manc 3) Sanc-COJO applied to European ancestry GWAS of the same total sample size (N=6,901^*^ 2) (**Table 1**). COJO SNPs were compared to the simulation ground truth (hereafter referred to as “causal SNPs”) using various criteria across 7,600 simulated scenarios in 38 LD independent blocks (see **Methods** and **Supplementary Methods**) with results summarised in **Figure 2**. Trends within individual LD blocks are generally consistent with the results pooled across blocks (**Supplementary Figure 1-2**).

**Table 1.**
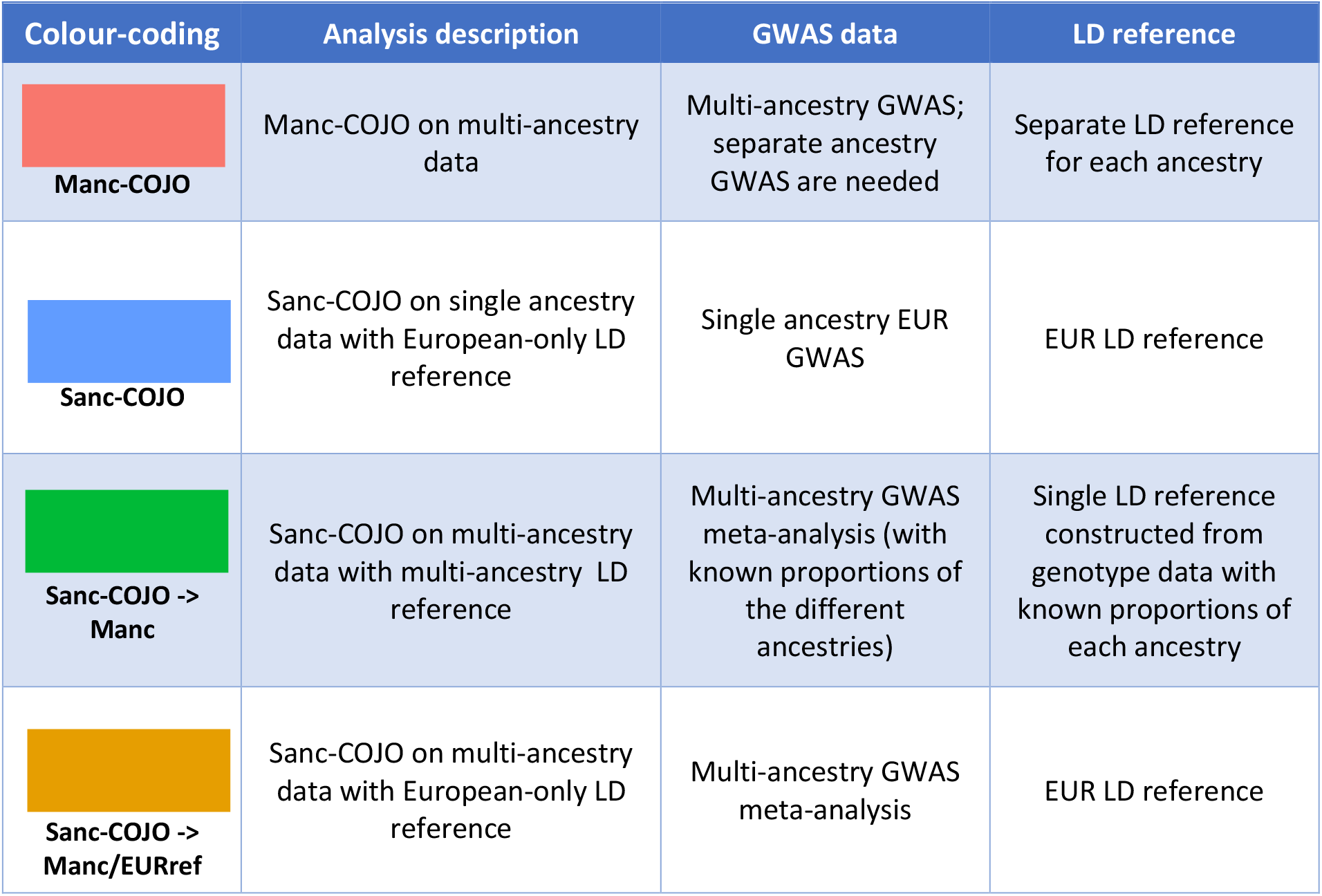
Summary of implemented methods. This table summarises the methods compared in this study with colour-coding matched to box plots used in Figures and Supplementary Figures. It clarifies the GWAS inputs and LD reference panels used for each method.

| Colour-coding | Analysis description | GWAS data | LD reference |
| --- | --- | --- | --- |
| **Manc-COJO** | Manc-COJO on multi-ancestry data | Multi-ancestry GWAS; separate ancestry GWAS are needed | Separate LD reference for each ancestry |
| **Sanc-COJO** | Sanc-COJO on single ancestry data with European-only LD reference | Single ancestry EUR GWAS | EUR LD reference |
| **Sanc-COJO -> Manc** | Sanc-COJO on multi-ancestry data with multi-ancestry LD reference | Multi-ancestry GWAS meta-analysis (with known proportions of the different ancestries) | Single LD reference constructed from genotype data with known proportions of each ancestry |
| **Sanc-COJO -> Manc/EURref** | Sanc-COJO on multi-ancestry data with European-only LD reference | Multi-ancestry GWAS meta-analysis | EUR LD reference |

**Figure 2.**
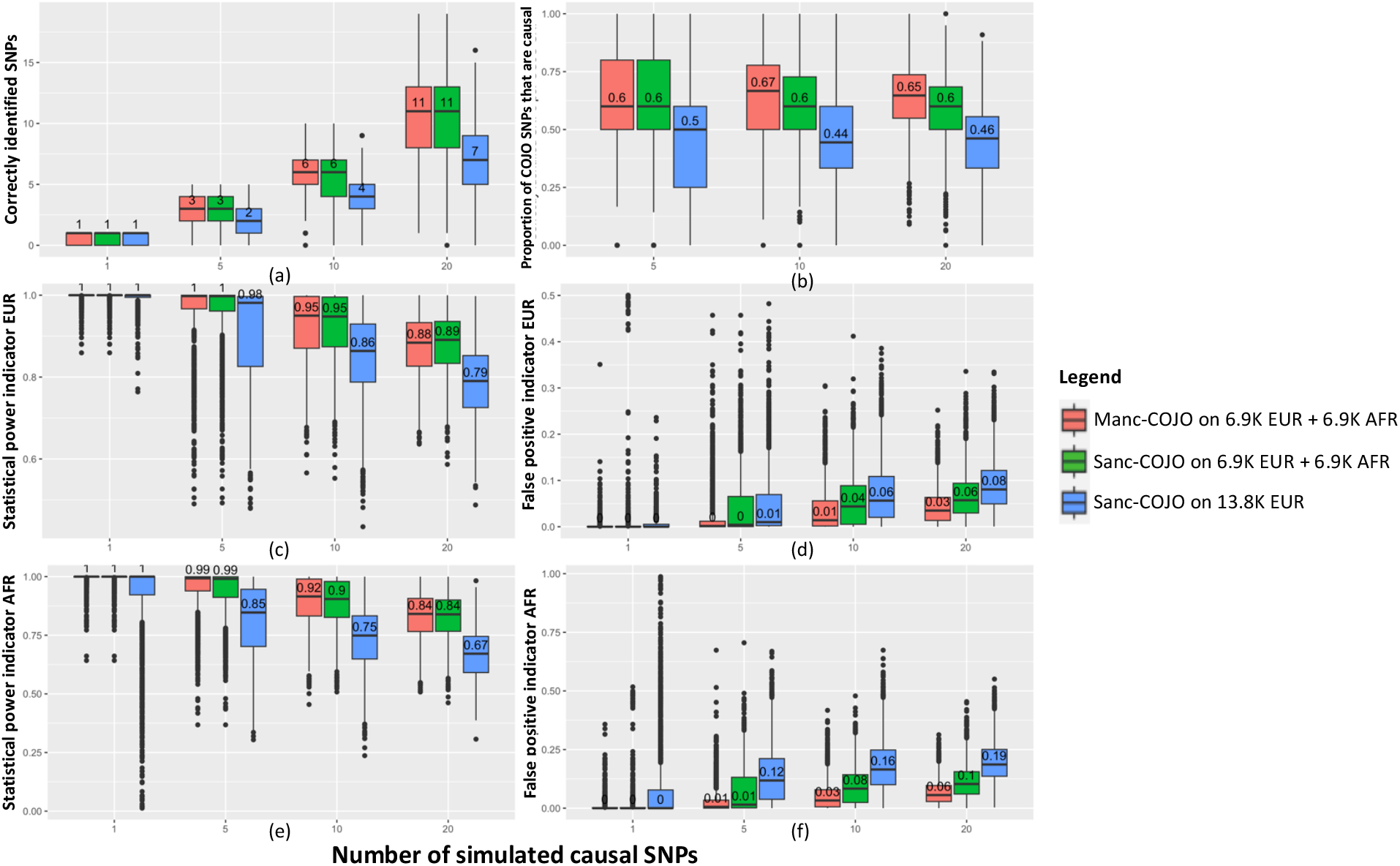
Manc-COJO on average identifies a similar number of causal SNPs, but exhibits superior power, and shows fewer false-positive discoveries in simulations. This figure summarises simulation results, and illustrates the performance of Manc-COJO in identifying causal SNPs, showcasing superior power and fewer false-positive discoveries. The algorithm was applied to simulations based on genotyped data of both a European (EUR) and African (AFR) cohort, each with a sample size of 6,900. The comparisons include Sanc-COJO applied to either 13,800 Europeans or a meta-analysis of 6,900 Europeans with 6,900 Africans. In all cases, in-sample LD was used. The x-axis represents the number of simulated causal SNPs per LD independent block using 38 cross-ancestry LD independent blocks on chromosome 22. (a) The y-axis denotes the number of causal SNPs correctly identified by each algorithm. (b) The y-axis denotes the proportion of SNPs identified by the COJO algorithm that are causal SNPs. (c) Given that COJO SNPs may not be the causal SNPs but maybe located in nearby locations, we identified the COJO SNP with the highest LD correlation (in European cohort) to each causal SNP and averaged these correlations across all causal SNPs. Higher value indicates better power in identifying causal variants. (d) For each COJO SNP, we identified the nearest true causal SNP, calculated their (European) LD, and then averaged these LD values across all COJO SNPs. We then subtracted this mean from 1 to obtain an indicator of false-positive discoveries (*i*.*e*., when selected SNPs are not in LD with causal SNPs, the average LD will be low, yielding a higher false-positive indicator). (e)(f) Power and false positive indicators as in (c) and (d), except that African LD is used in calculations. Each box plot is the distribution across 1,900 simulation replicate and displays the minimum, maximum, mean, and interquartile range. Numbers annotating each box plot are the mean values of each group.

Under idealised simulation conditions (i.e., well-powered GWAS in both ancestries, all causal SNPs genotyped, true SNP effects equal in both ancestries, and LD structure derived from the GWAS cohort), analyses incorporating multi-ancestry data (*i*.*e*.,Manc-COJO and Sanc-COJO->Manc) identify a greater number of causal SNPs (**Figure 2a,b**), exhibit superior statistical power (**Figure 2c,e**), and generate fewer false-positive discoveries (**Figure 2d,f**) compared to single-ancestry analyses with an equivalent total sample size (*i*.*e*., Sanc-COJO), gains which reflect the value of different LD patterns across ancestry group. Contrasting the two multi-ancestry approaches the median numbers of causal SNPs tagged by Manc-COJO versus Sanc-COJO->Manc are similar, but Manc-COJO produces fewer false-positive discoveries (**Figure 2d,f**), which becomes more evident when the simulated trait is highly polygenic (*i*.*e*., 10 or 20 causal SNPs within a block).

Out-of-sample prediction using the SNPs selected by a COJO model is an objective measure to compare validity of the SNPs selected by the different models. Despite Sanc-COJO resulting in a model with fewer true causal SNPs and more false-positive discoveries than Manc-COJO (and Sanc-COJO->Manc) from GWAS of the same total sample size, its out-of-sample prediction performance is comparable to that of Manc-COJO when predicting into an independent cohort of the same ancestry (**Extended Figure 1**). Hence, Sanc-COJO SNPs have good out-of-sample prediction accuracy within an independent sample sharing a similar LD structure to the GWAS samples. However, Sanc-COJO displays relatively low cross-ancestry portability, because the single ancestry COJO SNPs tag true causal SNPs less accurately than a multi-ancestry GWAS of the same total sample size which exploits differences in LD correlation between SNPs across different ancestries^24^.

In practice, model assumptions are often violated. To investigate the impact of such violations, we conducted a series of simulations in which one assumption was altered at a time. To mimic the scarcity of multi-ancestry data we simulated a scenario in which the European cohort sample size was increased ten-fold to 69,010. Compared with Sanc-COJO-based methods, Manc-COJO, on average, identifies a greater number of causal SNPs, exhibits superior power, and produces fewer false-positive discoveries in simulations (**Extended Figure 2**). This demonstrates that, even when sample sizes are heavily skewed towards European ancestry, incorporating African ancestry data still enhances the ability to detect independent associations with fewer false-positive discoveries. Notably, the application of Sanc-COJO to multi-ancestry data, Sanc-COJO→Manc—even when using an LD matrix constructed from the same individuals included in the analysis (i.e. in-sample mixed-ancestry LD)—results in a higher number of false-positive discoveries. Specifically, the proportion of SNPs selected by Sanc-COJO→Manc algorithm that are truly causal is 0.57, 0.50, and 0.48 for regions with 5, 10, and 20 causal SNPs, respectively. These values are not only lower than those obtained with Manc-COJO (0.80, 0.64, and 0.59 for the same scenarios) but also lower than those from single-ancestry Sanc-COJO (0.60, 0.55, and 0.50 for the same scenarios and total sample size).

When sample sizes are heavily biased toward European populations, some studies have applied a European-only LD reference even when the summary statistics were derived from multi-ancestry GWAS meta-analyses (i.e., they used the Sanc-COJO→Manc approach but with European-only LD references, hereafter referred to as Sanc-COJO→Manc/EURref)^5,7^. In simulations, when European ancestry dominates the sample (10:1 ratio), the approach performs reasonably well, achieving higher power in both ancestries and making fewer false positive discoveries in African ancestry, compared to Sanc-COJO applied to European-ancestry GWAS of the same total sample size (Manc-COJO is still superior). When the ancestry balance is equal (1:1 ratio), the use of a European-ancestry LD reference becomes ineffective, leading to marked inflation in false-positive discoveries, particularly in African ancestry samples (**Extended Figure 3**).

Other sensitivity analyses are reported in the **Supplementary Note 1** and **Supplementary Figures 3-5**. These included analysis on only HapMap3 SNPs to mimic when causal SNPs are not genotyped, simulations where effect sizes differ between ancestries and simulations where LD matrices were not calculated from the GWAS samples. As expected, the performance of all models declined under non-idealised conditions. Nonetheless, Manc-COJO consistently demonstrated greater robustness compared to the Sanc-COJO-based methods.

**Figure 3.**
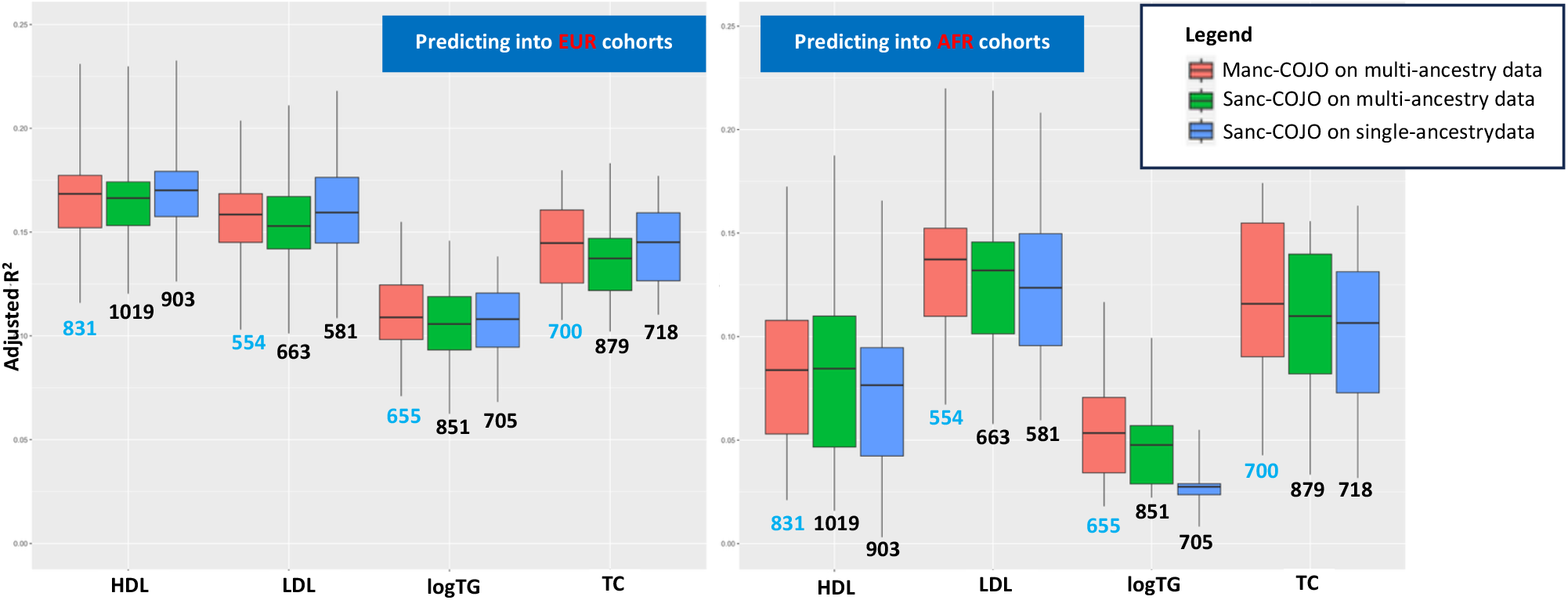
Manc-COJO selected fewer COJO-SNPs with superior trans-ancestry portability compared to the single-ancestry Sanc-COJO model. This figure summarises the out-of-sample prediction accuracies of Manc-COJO on multi-ancestry data, Sanc-COJO on multi-ancestry data, and Sanc-COJO on single-ancestry data for HDL, LDL, logTG, and TC when predicting into independent European and African cohorts. Numbers below each boxplot are the number of independently associated SNPs outputted from the algorithms, and the model with the smallest number of SNPs is highlighted in blue for each trait. (i) Manc-COJO input: European-specific GWAS summary statistics from GLGC (N ∼ 0.9 million), combined with African-specific GWAS from GLGC (N ∼ 90,000). All GWAS summary statistics excluded UK Biobank participants. The LD reference panels consisted of 69,010 unrelated European individuals and 6,901 unrelated African individuals from the UK Biobank. (ii) Sanc-COJO on multi-ancestry data input (Sanc-COJO->Manc): Meta-analysis of the European-specific and African-specific GWAS summary statistics described above. The LD reference panel was a merged set of the unrelated European and African individuals from the UK Biobank (69,010 Europeans and 6,901 Africans). (iii) Sanc-COJO on single-ancestry data input: European-specific GWAS used in i) meta-analysed with European-specific GWAS of additional independent UK Biobank participants (N ∼ 90,000) using pipelines consistent with the original GLGC studies. The LD reference panel merged the 69,010 unrelated Europeans with an additional independent sample of 6,901 unrelated Europeans from the UK Biobank. To ensure fair comparisons across methods, we harmonised the GWAS summary statistics to have identical SNP sets and approximately equal sample sizes, and we kept identical LD reference panel sizes across methods. Each box plot shows the distribution of the out-of-sample prediction R^2^ values for all groups (50 in EUR and 20 in AFR) and displays the minimum, maximum, mean, and interquartile range.

### Real data analysis

In real data analysis, since the true causal SNP and effect sizes are unknown, we use out-of-sample prediction accuracy from the COJO SNPs and model size as the primary criteria for comparing Sanc-COJO and Manc-COJO models. We analysed summary statistics from a GWAS meta-analysis of lipid traits (high-density lipoprotein cholesterol (HDL), low-density lipoprotein cholesterol (LDL), total cholesterol (TC), and log(triglyceride) (logTG)) including approximately 1 million samples (∼0.9 million EUR + ∼0.09 million AFR for multi-ancestry models, and ∼0.9 million EUR + ∼0.09 million EUR for single ancestry models; **Supplementary Table 1**).

We first assessed the prediction accuracy of single-ancestry Sanc-COJO- and Manc-COJO-derived predictors in an independent sample of 47,066 individuals of European ancestry. The prediction R^2^ (and standard error) of the Manc-COJO model was 0.168 (0.023), 0.159 (0.021), 0.109 (0.018), and 0.145 (0.020) for HDL, LDL, logTG, and TC, respectively. The R^2^ of the Sanc-COJO were similar: 0.170 (0.022), 0.159 (0.023), 0.108 (0.017), and 0.145 (0.020) for the same traits. When applied to an independent AFR cohort of 3,000 individuals, the Manc-COJO model consistently gave higher R^2^ values (although not statistically different given the sample size available) than Sanc-COJO: 0.084 (0.040) vs 0.077 (0.044) for HDL, 0.137 (0.046) vs 0.124 (0.040) for LDL, 0.0534 (0.028) vs 0.0274 (0.013) for logTG, and 0.116 (0.044) vs 0.107 (0.040) for TC. Notably, the number of independently associated SNPs identified by Manc-COJO was much lower than that of Sanc-COJO model: 831 vs 903 for HDL, 554 vs 581 for LDL, 655 vs 705 for logTG, and 700 vs 718 for TC (**Figure 3**). In summary, Manc-COJO achieved comparable predictive accuracy in the EUR sample and superior performance in the AFR sample, while selecting fewer SNPs, suggesting a lower rate of false-positive inclusions relative to the Sanc-COJO model. We replicated the out-of-sample prediction analysis in the All of Us cohort, and the results showed trends consistent with those observed in the UKB (**Table 2**).

**Table 2.** Replication of out-of-sample predictions in the All of Us cohort. This table summarises the out-of-sample prediction accuracies of the different methods in the All of Us cohort. Within each cell, the first row reports the model size; the second row reports the out-of-sample prediction accuracy among unrelated European participants; and the third row reports the corresponding accuracy among unrelated African participants. The observed pattern is consistent with the results from the UK Biobank (Figure 3): Manc-COJO selects fewer COJO SNPs while generally exhibiting superior trans-ancestry portability compared with the Sanc-COJO–based model.

| Method | HDL | LDL | logTG | TC |
| --- | --- | --- | --- | --- |
| **Manc-COJO** | 809<br>0.119 (0.021)<br>0.0830 (0.022) | 533<br>0.0914 (0.019)<br>0.100 (0.023) | 631<br>0.0876 (0.019)<br>0.0347 (0.016) | 693<br>0.0808 (0.018)<br>0.0843 (0.018) |
| **Sanc-COJO** | 884<br>0.122 (0.021)<br>0.0698 (0.019) | 565<br>0.0902 (0.019)<br>0.0866 (0.018) | 677<br>0.0911 (0.019)<br>0.0297 (0.016) | 700<br>0.0815 (0.019)<br>0.0665 (0.017) |
| **Sanc-COJO -> Manc** | 1022<br>0.117 (0.021)<br>0.0782 (0.020) | 638<br>0.0852 (0.018)<br>0.0949 (0.024) | 829<br>0.0871 (0.020)<br>0.0337 (0.015) | 880<br>0.0801 (0.019)<br>0.0755 (0.016) |
| **Sanc-COJO -> Manc/EURref** | 845<br>0.118 (0.021)<br>0.0818 (0.022) | 593<br>0.0903 (0.019)<br>0.0955 (0.020) | 650<br>0.0868 (0.020)<br>0.0382 (0.017) | 730<br>0.0813 (0.018)<br>0.0800 (0.015) |

In previous attempts to adapt Sanc-COJO for multi-ancestry GWAS, an *ad hoc* approach has employed LD matrices estimated from samples whose ancestry composition matches that of the GWAS sample^9,10^. In real-data analyses, the resulting models were substantially larger than those produced by Manc-COJO: the number of SNPs selected by the Sanc-COJO→Manc model was 1,019 for HDL, 663 for LDL, 851 for logTG, and 879 for TC—approximately 25% more SNPs than identified by Manc-COJO models (**Figure 3**). Despite this increased model complexity, predictive performance was marginally lower than that of Manc-COJO across all traits and scenarios. These results indicate that the Sanc-COJO→Manc model is more prone to false-positive SNP inclusion than the more parsimonious Manc-COJO approach, in agreement with the simulation findings.

In some studies^5,7^ Sanc-COJO has been applied to meta-analyses of multi-ancestry GWAS using EUR-only LD reference panels, particularly in studies where EUR populations dominate the sample sizes. The results aligned with observations from our simulations. When the European-to-African (EUR: AFR) sample size ratio was 10:1, this approach was reasonably effective— Sanc-COJO->Manc/EURref produced models with comparable out-of-sample prediction accuracy in both European and African cohorts, and the model sizes remained similar. However, as the ancestry balance shifted to a 1:1 ratio, this approach was no longer viable (evaluated through down-sampling; **Supplementary Note 2**). The number of SNPs selected by Sanc-COJO->Manc/EURref model increased substantially, reaching approximately twice the size of the corresponding Manc-COJO models (**Extended Figure 4**). Moreover, for all traits except TC, these model sizes also exceeded those of Sanc-COJO->Manc models (which were constructed with ancestry-proportion-matched LD reference panels), suggesting a higher rate of false-positive SNP inclusion.

### Manc-COJO:MDISA

All simulations and real-data analyses described above assumed shared causal SNPs across ancestries. We developed a complementary approach multi-ancestry data-informed single ancestry analysis (Manc-COJO:MDISA) which uses multi-ancestry information to increase power to identify variants which may have ancestry-specific effects. We designed a series of simulated scenarios in which 60% of the simulated SNPs in a block are common across ancestries, while 20% are specific to each of the two ancestries (**Methods**). Both multi-ancestry algorithms (Manc-COJO and Sanc-COJO->Manc) detect more shared causal variants than Sanc-COJO of the same total GWAS sample size, with Manc-COJO outperforming Sanc-COJO->Manc. Applying Manc-COJO:MDISA to the Manc-COJO output recovers around 10% - 20% additional ancestry-specific causal variants (**Figure 4**). In real data analysis, we applied Manc-COJO:MDISA (using ∼6.5 million SNPs) within the ∼1 million EUR and ∼0.09 million AFR cohorts separately to test for ancestry-specific associations. No new SNPs were identified in the AFR cohort. In the EUR cohort, compared with the Manc-COJO model, the increase in prediction R^2^ (and corresponding increase in model size) was 0.0007 (8), 0.0003 (4), 0.0005 (17), and 0.0002 (2) for HDL, LDL, logTG, and TC, respectively. Hence, most of the prediction R^2^ can be attributed to models based on the assumption of shared common causal SNPs across ancestries.

**Figure 4.**
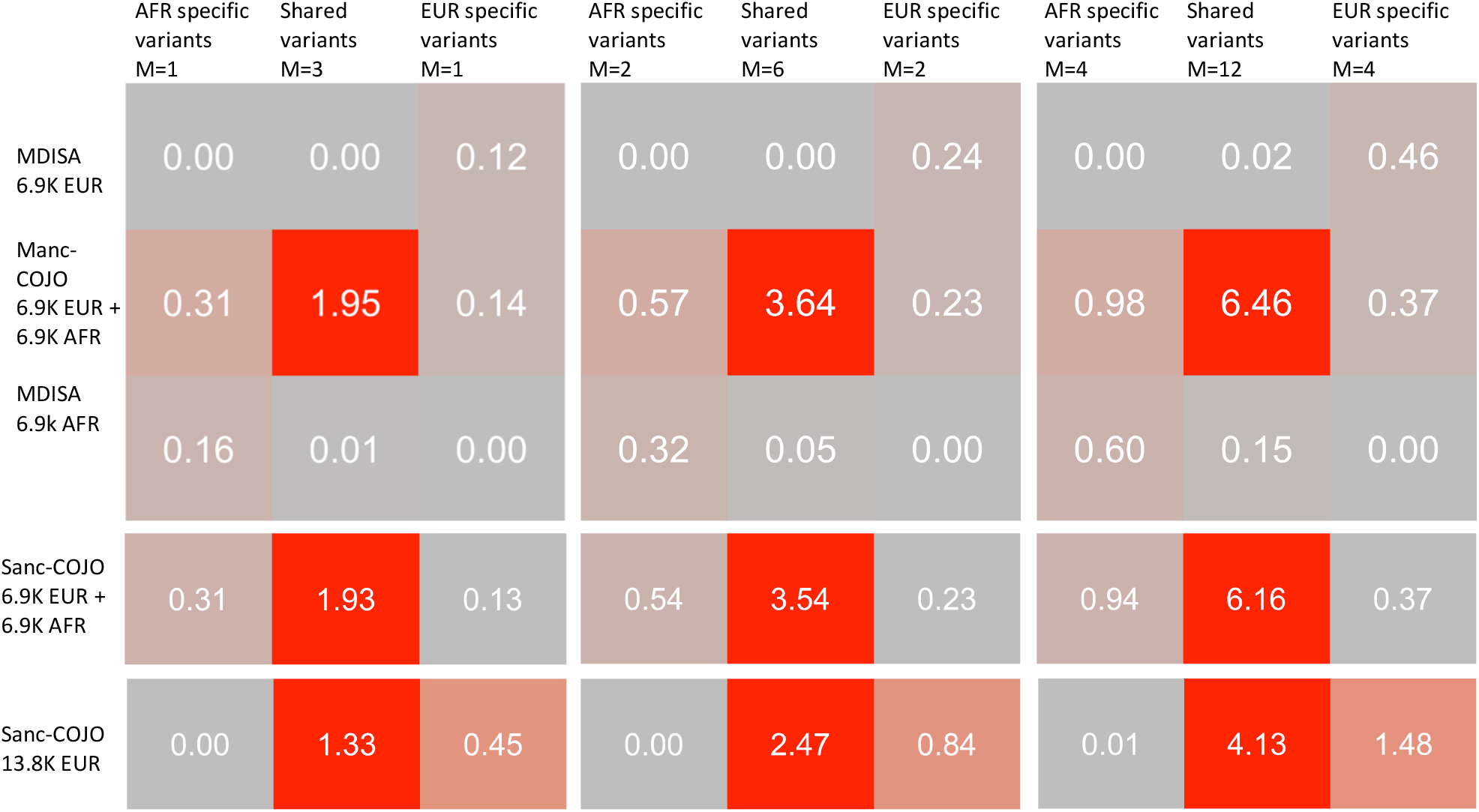
Manc-COJO identifies a greater number of shared causal SNPs in scenarios where ancestry-specific trait-associated variants exist, while Manc-COJO:MDISA recovers some ancestry-specific associations. This figure summarises results from simulations applied to 38 cross-ancestry LD independent blocks on chromosome 22, where 60% of the simulated SNPs in each block are common across ancestries and 20% are specific to each of the two ancestries. Each scenario, defined by the number of causal SNPs (5, 10, or 20), consists of 1,900 simulation replicates (38 blocks × 50 within-block replicates). Column labels indicate the number of simulated SNPs per block (M) and whether they are shared across ancestries or ancestry-specific. Row labels correspond to the analyses performed. The numbers in the boxes are the mean number of correctly identified SNPs calculated by averaging across 38 blocks and 1,900 simulation replicates. Consistent with prior simulations, we applied Manc-COJO to a simulated European cohort (*N* = 6,900) and a simulated African cohort (*N* = 6,900). We applied Manc-COJO:MDISA within each ancestry, fixing the SNPs selected by Manc-COJO to be in the model. Comparisons include Sanc-COJO applied to either 13,800 Europeans or a meta-analysis of 6,900 Europeans and 6,900 Africans, with in-sample LD utilised in all cases. Numbers in the plot represent the mean number of causal SNPs correctly identified by each algorithm within each category, with a colour gradient reflecting these values (red for higher numbers, grey for lower). Both multi-ancestry algorithms—Manc-COJO and Sanc-COJO applied to multi-ancestry data—detect more shared causal variants than a single-ancestry approach with the same sample size, with Manc-COJO outperforming Sanc-COJO on multi-ancestry data. By applying Manc-COJO:MDISA to the Manc-COJO output, we are able to recover some additional ancestry-specific associations.

### Software improvement compared to Sanc-COJO

Although our software is designed primarily for multi-ancestry analysis, it also supports running single-ancestry COJO as implemented in GCTA. We have introduced several coding improvements to the original Sanc-COJO implementation to make the output more robust (**Supplementary Note 3**). Importantly, our software achieves substantially greater computational efficiency than the current GCTA implementation. We benchmarked the total running time across all 22 chromosomes for single-ancestry analysis in ∼1 million EUR individuals (∼6.5 million SNPs; as described in the real-data analysis), using ∼75,000 unrelated UKB EUR samples as the LD reference. With five threads, our software required only 5.80, 4.83, 5.12, and 5.08 minutes for HDL, LDL, TC, and logTG, respectively (i.e., on average <30 seconds per chromosome when run in parallel). In contrast, GCTA required 914.80, 623.48, 756.12, and 719.92 minutes under the same CPU and memory settings, corresponding to speedups of 157.65, 129.02, 147.78, and 141.82 times, respectively. The design principles underlying these improvements may be relevant to other GWAS summary statistics software (**Extended Figure 5; Supplementary Note 4**).

### Insights into Manc-COJO and Sanc-COJO outputs

To assess and understand the differences between Manc-COJO and Sanc-COJO in a real-world data context, we compared their outputs in genomic regions surrounding SNPs that have been experimentally validated as causal through wet-lab studies. Previous work has identified the following functional SNP–gene pairs—together with their potentially associated lipid traits— using CRISPR-based assays: rs12740374–*SORT1*(LDL)^25^, rs2277862–*CPNE1*(HDL and TC), rs10889356–*DOCK7/ANGPTL3*(TG and TC), and rs10872142–*FRK*(LDL)^26^. In LDL, both Manc- and Sanc-COJO included rs12740374 into the model. For the remaining three SNPs, both Manc-COJO and Sanc-COJO selected variants that were in high LD with the experimentally validated causal SNPs in European ancestry samples. However, in African ancestry samples, Manc-COJO tends to select variants that were more closely correlated with the causal SNPs than those selected by Sanc-COJO. For example, for rs2277862 in TC, Manc-COJO selected rs12625762, which has signed LD correlations (r) of 0.994 (EUR) and 0.942 (AFR) with the causal SNP. In contrast, Sanc-COJO selected rs1010759, with corresponding LD correlations of 0.992 (EUR) and 0.918 (AFR). A similar pattern occurred for rs10889356 in TC: Manc-COJO selected rs995000, whereas Sanc-COJO selected rs626787. The LD correlations with the causal SNP were 0.962 (EUR) and 0.960 (AFR) for the Manc-COJO–selected SNP, compared with 0.955 (EUR) and –0.476 (AFR) for the Sanc-COJO– selected SNP. For rs10872142 in LDL, Manc-COJO selected rs13210143, which shows extremely high LD with the causal SNP in both EUR (0.998) and AFR (1.000). In contrast, Sanc-COJO selected rs1556857, whose LD correlations were 0.996 (EUR) and 0.899 (AFR) (**Figure 5; Supplementary Methods**).

**Figure 5.**
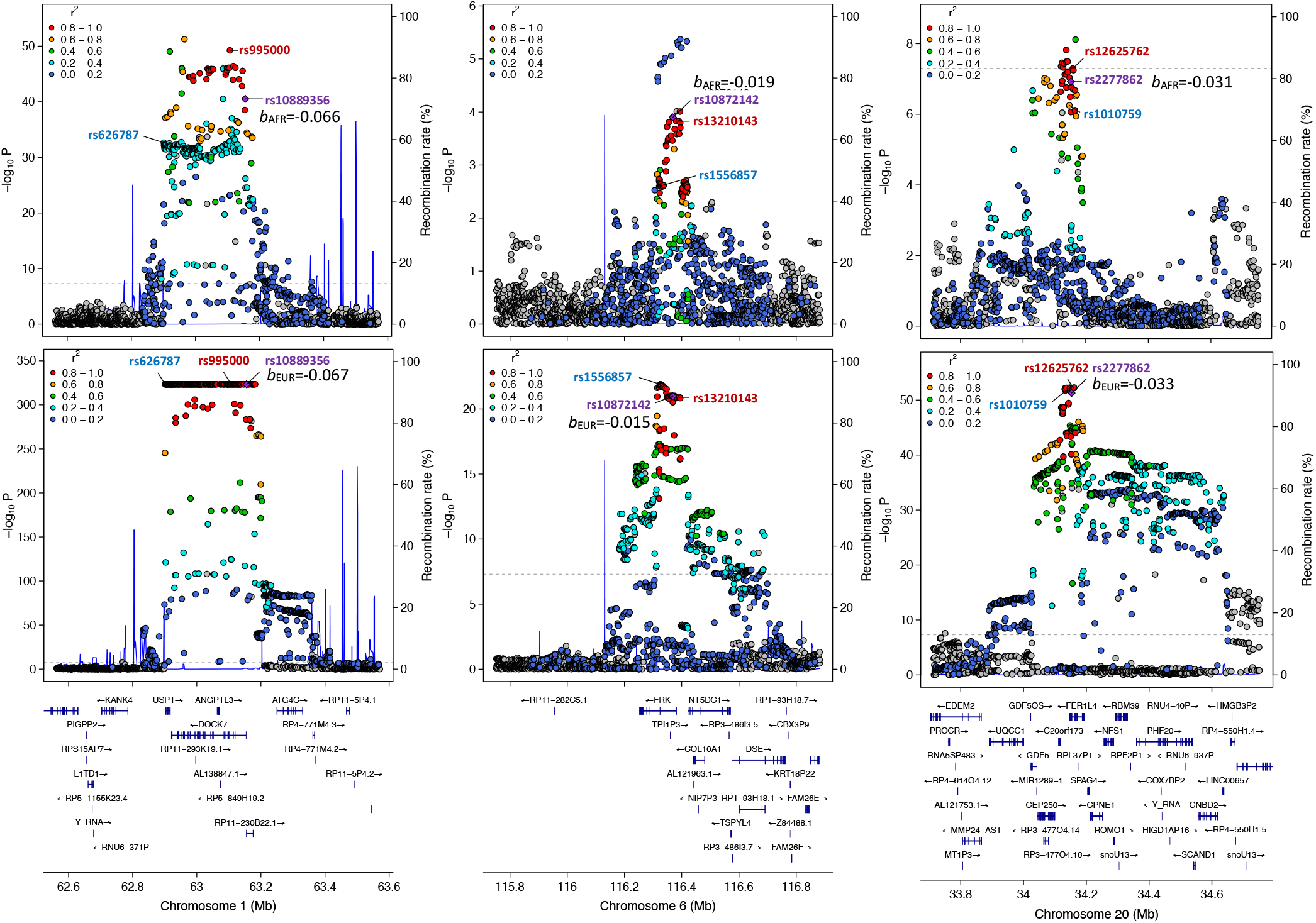
LocusZoom plots highlighting Manc-COJO vs Sanc-COJO SNPs in regions and for traits where the causal SNP is known. This figure shows regional LD and recombination structure near three wet-lab-validated causal variants for lipid traits. The left column corresponds to the rs10889356–DOCK7/ANGPTL3 SNP–gene pair in TC, the middle column shows rs10872142–FRK in LDL, and the right column shows rs2277862–CPNE1 in TC. The top row displays LD structure and GWAS summary statistics from AFR data (as in Figure 3), and the bottom row shows the corresponding EUR results. Wet-lab-validated variants are labelled in purple, SNPs flagged by Manc-COJO are highlighted in red, and those flagged by Sanc-COJO are highlighted in blue. Point colours indicate the squared LD correlation with the validated causal SNPs, and numbers next to causal variants denote their marginal effect sizes in the GWAS. LD structure in EUR is clearly more complex—particularly in the lower left panel—where several SNPs are in near-complete LD owing to limited recombination. By leveraging distinct LD patterns in African-ancestry cohorts, we can better disentangle LD blocks and identify independently associated variants that more closely reflect the true causal alleles. It is also notable that effect sizes for the causal variants are broadly similar across ancestries.

Similar trends were observed in out-of-sample prediction accuracies although the out-of-sample prediction sample sizes are too small to support statistically significant differences (see **Supplementary Note 5**).

## Discussion

We developed a novel multi-ancestry algorithm to perform conditional and joint analysis of GWAS summary statistics. We demonstrate through extensive simulations that Manc-COJO consistently outperformed Sanc-COJO applied on both single- and multi-ancestry data by correctly identifying, on average, more independent associated variants and reducing false positive associations. Moreover, Manc-COJO was robust to scenarios when sample sizes are biased, causal SNPs not all genotyped, true SNP effects different across ancestries, and imperfect LD references. We applied Manc-COJO to four polygenic lipid traits (which have genetic architectures that include large effect size associations) and validate SNPs selected by Manc-COJO using out of sample prediction and observed Manc-COJO yielded more parsimonious models with improved trans-ancestry portability compared to single ancestry Sanc-COJO model of the same sample size.

In real data applications, the differences between Manc-COJO and Sanc-COJO models in terms of variance explained by their COJO SNPs were not statistically significant. The lack of statistical significance likely reflects power (sample sizes of non-European GWAS are currently much smaller than for European GWAS, 10-fold for the lipid traits studied here, and non-European sample sizes for out-of-sample prediction are also limited). Nonetheless, the pattern of results, replicated across the traits and databases studied is consistent with expectations from well-powered simulations. The use of Manc-COJO applied to cross-ancestry GWAS meta-analyses will become increasingly relevant as samples sizes for non-European GWAS increase over time. Moreover, our results provide insights likely relevant to other post-GWAS methods that rely on GWAS summary statistics and LD references.

A key result from our analyses is that Manc-COJO achieves comparable predictive accuracy in European ancestry cohorts and markedly better portability in African ancestry cohorts, while selecting fewer SNPs. This indicates that the number of independent associations detected using single-ancestry data may be overestimated and tagging of true causal variants is improved through breakdown of the correlation structure in the genome which can be exploited even when the proportion of African ancestry represented in the GWAS is relatively modest (*e*.*g*., 10%).

Previous efforts to adapt Sanc-COJO for identification of independent associations in multi-ancestry GWAS meta-analyses likely include false discoveries, although current sample sizes mean the impact is limited. Using LD estimated in European ancestry samples in multi-ancestry analysis performs adequately when the EUR:AFR sample size ratio is around 10:1, but it leads to increasing false positives as this ratio approaches 1:1. Pinpointing the exact threshold where performance deteriorates is challenging (and likely varies by chromosomal region and genetic architecture). Likewise, constructing LD reference panels that match the ancestry proportions of the GWAS can work well under idealised conditions (e.g., in-sample LD, balanced ancestral representation, and sufficient power) but becomes prone to inflated false discoveries when these assumptions are violated. One potential contributing factor to inflated false discoveries is that all COJO models assume Hardy–Weinberg equilibrium (HWE) in their calculations; however, mixed-ancestry LD reference panels may violate this assumption (**Supplementary Note 6**).

Here, we focussed on individuals of inferred European and African ancestry as a proof of principle. However, the Manc-COJO methodology is readily extendible, and our software supports the inclusion of an arbitrary number of ancestries—limited only by computational and memory resources. Although this paper applies meta-analysis strategies to integrate data across ancestries, the software can also be adapted to perform mega-analysis (i.e., pooling individual-level genotype data) or analyses within admixed population, which in principle may offer greater power when in-sample LD is available and sample sizes are sufficiently large^27^, but may be prone to inflated false positives when these ideal conditions are not met (**Supplementary Note 6**).

This study has several limitations, some inherent to the nature of the algorithm, others attributable to data quality and the scarcity of real data. First, we made several assumptions: true causal SNPs act additively on the phenotype, causal SNPs are independent, and the number of true causal variants, as well as their effect sizes on standardised phenotypes, are identical across ancestries. These assumptions can be violated in various ways. For instance, dominant or recessive SNP effects may exist^28^ and/or may interact with other SNPs^29^, but these can only be identified in the context of large SNP effects. Previous research has shown that the additive effect model explains most genetic variation in complex traits^30,31^.

Second, although an increasing number of studies provide empirical evidence that causal SNPs are shared across ancestries with similar effect sizes^18-21^, ancestry-specific causal variants have also been reported^22,23^. To address this, we developed a complementary method, Manc-COJO:MDISA, to accommodate such scenarios. In real-data applications, this algorithm identified only a small number of additional variants in the EUR cohort. Importantly, these variants should not be interpreted as definitive evidence of ancestry-specific effects for lipid traits. On one hand, the additional variants accounted for only a negligible increase in out-of-sample prediction R^2^. On the other hand, their absence in AFR may simply reflect limited statistical power, causing true causal variants to fall below significance thresholds. Thus, at least in the near term, bioinformatics-based claims of ancestry-specific variants should be supported by direct functional validation.

With this in mind, we designed our software to allow users to fix selected SNPs in the model. Traditionally, bioinformatics analyses serve as an upstream step for wet-lab validation. However, our software enables a “two-way street”: causal variants confirmed experimentally can be incorporated into the model—even if they are not the most statistically significant (due to sampling variability) or do not meet genome-wide significance thresholds (due to limited sample sizes)—and will not be removed during subsequent variable selection. Incorporating wet-lab– validated variants directly into the modelling framework may further enhance the detection of independent association signals.

Throughout this paper, we frequently refer to causal variants and benchmark simulation outcomes against them, raising the question of whether COJO should be considered a fine-mapping tool. Simulations show that, in an idealised scenario where all causal variants are genotyped, no pairs of causal variants exhibit LD above 0.8, and studies are well-powered, Sanc-COJO identifies 40% of causal variants for less polygenic traits (5 causal SNPs in a region) and 16% for more polygenic traits (20 causal SNPs in a region). Manc-COJO increases this to 60% and 40%, respectively, by leveraging differing LD structures across ancestries. However, the assumptions (common to all fine-mapping methods) that all causal variants are genotyped (or imputed) and studies are well-powered are rarely met in practice. With whole-genome sequence data, where nearly all genetic variants are genotyped (though gaps remain, especially with short-read sequencing^32^), many variants are in complete or near-complete LD, necessitating indefinitely large sample sizes to completely resolve LD. Thus, Manc-COJO outcomes should be interpreted as a model of variants proximal to true causal variants that are nearly independently associated. In other words, while fine-mapping methods aim to pinpoint (using credible sets) the exact causal variant, Manc-COJO seeks to construct a model with nearly independent variants, where tagging variants close to causal ones suffices.

In summary, our study underscores the importance of collecting data from diverse ancestries, particularly as GWAS for some traits approach saturation in European ancestries^3^. It is essential to allocate resources to underrepresented populations during data collection efforts. With multi-ancestral data, approaches like Manc-COJO can better resolve LD and refine candidate variants for experimental validation. Given that LD blocks are smallest in African ancestry cohorts, GWAS in African ancestry cohorts will be the most cost-effective for optimisation of Manc-COJO SNPs as well as for fine-mapping methods.

## Methods

### Manc-COJO algorithm

Briefly, the Manc-COJO algorithm assumes GWAS summary statistics from two or more ancestries. The LD reference panel for each ancestry should closely match the cohort from which the GWAS was generated. The multi-ancestry COJO algorithm is summarised in eight steps (**Figure 1**) and, like Sanc-COJO, requires a pre-specified genome-wide significance threshold (here 5 × 10^-8^) and a pre-set collinearity threshold between SNPs (i.e. maximum LD *r*^2^, here 0.9).

**Step 1:** Conduct an inverse-variance meta-analysis (IVM) of GWAS summary statistics across all ancestries. The SNP with the smallest meta-analysis p-value is selected for inclusion in the model.

**Step 2:** SNPs with an LD correlation above the pre-set collinearity threshold with any SNP already included in the model in any ancestry are removed in all ancestries. This assumes that the SNP selected from the LD block is the best SNP to represent the true causal SNP in this region.

**Step 3:** Within each ancestry, a conditional analysis is performed for each SNP *i* not currently included, and the conditional effect estimate is:

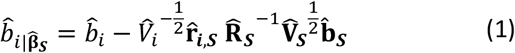

where,

- 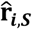 is a 1 x *s* vector of **observed** (signed) LD correlations between SNP *i* and each of *s* SNPs already selected into the model (in the first iteration, only 1 SNP is in the model)
- 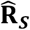 is an *s* x *s* matrix of **observed** (signed) LD correlations between s SNPs that are already in the model (in the first iteration, this reduces to a 1×1 matrix with the value equal to 1)
- 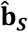 is an *s* x 1 vector of **observed marginal effects** of *s* SNPs in the model
- 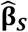 is an *s* x 1 vector of **observed joint effects** of *s* SNPs in the model
- 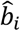 is the **observed** marginal effect of the SNP *i*
- 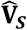 is an *s* x *s* diagonal matrix with *s*-th elements 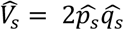 (assuming HWE)
- 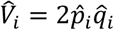 (assuming HWE)

**Step 4:** The conditional association statistic is estimated for each SNP (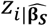). As in Sanc-COJO, to avoid over-fitting, a conservative approach is used

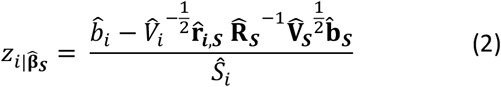

where 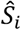 is the standard error of the marginal effect of SNP *i*. The conditional p-value is then calculated for each SNP and the SNP with the smallest meta-analysis p-value is selected for inclusion in the model.

**Step 5:** After getting the z-statistics for each of the remaining SNPs, the SNP with the smallest p-value is selected into the model (i.e. the SNP with largest z^2^). Afterwards, a joint analysis is performed as follows:

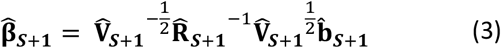

Here, the definition of 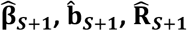, and 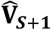, are the same as defined in equation (1), except that now there are *S*+1 SNPs in the model. As in Sanc-COJO, the estimation error of joint effects of *S*+1 SNPs can be expressed as:

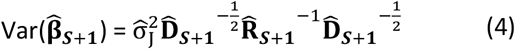

Where 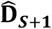 is an (*S*+1)x(*S*+1) diagonal matrix with *s*-th elements 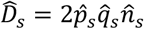 _+_, and 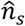 is the effect sample sizes of SNP *s*. Details on the estimation of 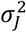 can be found in the **Supplementary Methods**.

**Step 6:** An IVM of the joint effects of the SNPs included in the model is performed, and any SNP whose meta-analysis p-value for the joint effects—or whose inclusion results in an adjusted *R*^’^(an optional criterion; **Supplementary Note 7**) that exceeds the pre-set threshold—is removed from the model.

**Step 7:** Steps 2 through 6 are repeated until no SNP can be added to or removed from the model.

**Step 8 (Manc-COJO:MDISA):**

To detect ancestry-specific associations, single-ancestry COJO is applied to each individual ancestry GWAS using models in which all Manc-COJO SNPs are included prior to the iterative addition or removal of SNPs. The Manc-COJO SNPs are fixed within the model and are not subject to exclusion during subsequent variable selection iterations.

Further details of the algorithm are provided in the **Supplementary Methods**.

### Simulations

Full details of the simulation framework are provided in the **Supplementary Methods**. Briefly, we used real genotype data from chromosome 22 of UK Biobank participants to construct our simulations. The UK biobank recruited approximately 500,000 people with 88% recorded as white British. Three cohorts were defined: an African cohort (AFR1), comprising 6,901 unrelated participants of inferred African ancestry; and two European cohorts (EUR1 and EUR2), each including 6,901 randomly selected, unrelated individuals of inferred European ancestry, with no overlap or relatedness between EUR1 and EUR2^24^. EUR1 and EUR2 were then combined to form the EUR1_EUR2 cohort (N = 13,802); AFR1 and EUR1 were then combined to form the AFR1_EUR1 cohort (N = 13,802). The sample sizes of EUR1 and EUR2 were matched to AFR1 to reflect previous findings that combining equal-sized European and African ancestry cohorts maximises power for detecting causal associations in multi-ancestry meta-analyses^33,34^. Within these cohorts, chromosome 22 was partitioned into blocks following the definition by Berisa and Pickrell^35^, and we selected 38 blocks for simulation. In each block, we randomly designated 1, 5, 15, or 20 SNPs as causal variants, and to mitigate collinearity, LD correlations between all pairs of causal SNPs were restricted to ≤ 0.8. The effect sizes of these variants were simulated to be identical across ancestries, ensuring that SNP heritability for each causal SNP ranged between 0.5% and 1.5% in all European and African cohorts. Necessarily, if effect sizes are held constant across ancestries, then the contributions to variance explained will differ across ancestries depending on SNP allele frequency. For each combination of block and number of causal variants, the simulation was repeated 50 times, drawing different sets of causal variants and effect sizes in each iteration. This design yielded 7,600 scenarios in total, reflecting 38 blocks, four levels of causal SNP count, and 50 replicates of causal SNP sets and effect sizes.

Using these simulated causal SNP sets and effect sizes, phenotypes were generated within each of the defined cohorts. For each of the 7,600 scenarios, a GWAS was conducted in each cohort using PLINK 2.0 (--glm hide-covar cols=+a1freq). An inverse-variance meta-analysis of summary statistics from EUR1 and AFR1 was then performed using custom R code to create the AFR1_EUR1 cohort. We compared outputs across the following GWAS and model combinations: Manc-COJO with EUR1 and AFR1, Sanc-COJO with EUR1_EUR2 and Sanc-COJO->Manc using AFR1_EUR1. Default settings were applied in all scenarios for both Sanc-COJO and Manc-COJO, with in-sample LD used throughout. For the AFR1_EUR1 cohort, in-sample LD was estimated from the merged genotypes of AFR1 and EUR1 using PLINK 1.9 (--bmerge). For Sanc-COJO->Manc, we also applied a European-only LD reference (EUR1_EUR2 in this case) in addition to the in-sample mixed-ancestry LD (**Table 1**).

To assess model performance, we established several benchmarking criteria, described in detail in the **Supplementary Methods**. These included the number of COJO-selected SNPs that were true causal variants, the proportion of COJO-selected SNPs that were causal, and metrics reflecting both power and false-positive discoveries. In addition, we assessed out-of-sample prediction accuracy using the estimated joint SNP effect sizes of COJO-selected SNPs in independent European (N = 13,800, randomly divided into 23 groups of 600) and African ancestry cohorts (N = 2,000, randomly divided into 20 groups of 100). Although the COJO algorithm is not designed to optimise out-of-sample risk prediction, comparing out-of-sample prediction using polygenic scores (PGS) from COJO-selected SNPs provides a practical evaluation criterion that is relevant in both real-data analyses and simulation studies.

We also assessed the performance of different COJO algorithms under several non-idealised scenarios (see **Supplementary Methods**). Briefly, we repeated the above analyses under conditions where in-sample LD references were unavailable—using a random sample of *N* = 503 European and 503 African participants from the 1000 Genomes Project as the LD reference^36^. Additional scenarios included: biased sample sizes favouring the European population at a ratio of 10:1; ancestry-specific effect sizes (but with consistent directions of association across ancestries); ancestry-specific causal variants; and incomplete genotyping of causal variants, in which case we restricted the analysis to HapMap3 SNPs only^37^.

### Real data analysis

We evaluated the performance of Manc-COJO in real-world data, using ancestry-specific GWAS summary statistics of European and African ancestries for four lipid traits chosen because of their large sample sizes for both European and African ancestries. Global Lipids Genetics Consortium (GLGC) GWAS summary statistics (which excluded UK biobank) were available for high-density lipoprotein cholesterol (HDL), low-density lipoprotein cholesterol (LDL), total cholesterol (TC), and log-transformed triglyceride levels (logTG)^38^. Only common SNPs (minor allele frequency, MAF ≥ 0.01) present in both ancestries and bi-allelic SNPs were retained for analysis. To allow for out-of-sample prediction analysis into the UK biobank, only SNPs available in the UK Biobank across both ancestries and had an MAF ≥ 0.01 among unrelated African participants (N = 6,901; genetic relatedness <5%) were retained for Manc-COJO analysis.

For each trait, Manc-COJO input files were GWAS summary statistics (one for each ancestry) and corresponding ancestry specific LD reference panels (i.e., a randomly sampled set of 69,010 unrelated European individuals and 6,901 unrelated African individuals from the UK Biobank). Default parameters were applied: a window size of 10Mb (assuming SNPs more than 10Mb apart to be independent), a collinearity threshold of r^2^ < 0.9, and a significance threshold of p < 5×10^-8^, consistent with the original Sanc-COJO.

We benchmarked Manc-COJO against Sanc-COJO in both European-only meta-analysis and inverse-variance-weighted meta-analysis combining multiple ancestries (multi-ancestry meta-analysis). Sanc-COJO was applied under three distinct scenarios: (1) single-ancestry meta-analysis with the European-only LD reference, (2) multi-ancestry meta-analysis with the ancestry-matched LD reference, and (3) multi-ancestry meta-analysis with the European-only LD reference (**Supplementary Table 1-2**).

To enable a fair comparison based on equal sample sizes across scenarios, we generated GWAS summary statistics for each lipid trait using 90,000 unrelated UK Biobank participants of inferred European ancestry (see **Supplementary Methods; Supplementary Table 2**). These UKB results were then meta-analysed with the European-specific GWAS from the GLGC to construct a European-only meta-analysis.

LD reference panels for Sanc-COJO based methods were created using PLINK’s --bmerge function, from either a merged cohort comprising 69,010 individuals of European ancestry and 6,901 individuals of African ancestry (ancestry-matched LD reference), or a merged cohort of 69,010 and 6,901 individuals both of European ancestry (European-only LD reference).

Given that the GLGC GWAS was heavily skewed towards European ancestry—with an approximate 10:1 ratio of European:African participants following the exclusion of UK Biobank samples—we conducted a down-sampling analysis to investigate a more balanced ancestral representation (approximately 1:1). To do this, we employed GWAS summary statistics derived from the GLGC African cohort (*N* ∼ 0.09 million) and then used a UKB European subset of the same *N*. For the single-ancestry analyses, GWAS summary statistics were generated from 180,000 UKB participants of European ancestry for each lipid trait, following the same analytical pipeline used in the full-scale analysis (**Supplementary Table 3; Supplementary Note 2**). The multi-ancestry LD reference panel consisted of 6,901 unrelated European and 6,901 unrelated African individuals, whereas the European-only LD reference comprised 13,802 unrelated European individuals, excluding those included in the GWAS. In this analysis, we again benchmarked Manc-COJO applied to multi-ancestry data against three alternative approaches: (1) Sanc-COJO applied to meta-analysed multi-ancestry data using an ancestry-matched LD reference (Sanc-COJO->Manc); (2) Sanc-COJO applied to meta-analysed multi-ancestry data using a European-only LD reference (Sanc-COJO->Manc/EURref); and (3) Sanc-COJO applied to single-ancestry GWAS using a European-only LD reference.

To evaluate objectively the Manc-COJO SNP sets we used out-of-sample predictive performance. We used 47,066 unrelated UK Biobank participants of inferred European ancestry who were not included in both the LD reference panel and GWAS analyses, as well as 3,000 participants of inferred African ancestry from the UK Biobank. Residuals for all lipid traits were computed using the same covariate adjustments as applied in the GLGC GWAS analyses (see **Supplementary Methods**). SNP effect sizes estimated from jointly fitting all SNPs identified by each Manc-COJO and Sanc-COJO model using the --cojo-joint function were used as SNP allele weights in PGS. This was performed separately for the European-only meta-analysis and the African ancestry GWAS summary statistics from the GLGC consortium (**Supplementary Table 2-3**). PGS were generated using PLINK (“—score” command). To estimate the prediction error, we randomly divided 47,066 unrelated UK Biobank European participants for lipid traits into 50 groups, and 3,000 unrelated UK Biobank African participants into 20 groups, with one group having more individuals if the sample size was not evenly divisible. Within each group, the out-of-sample R^2^ from a simple linear regression (residual ∼ PGS + error) was calculated and averaged across groups for each trait and ancestry, and the standard error of the estimate was computed.

We repeated our analysis of out-of-sample predictive performance using the All of Us cohort. Because SNPs in the All of Us data are aligned to the GRCh38 reference genome, we first constructed a conversion table between SNP names and genomic coordinates in the UK Biobank (GRCh37) and All of Us (GRCh38) cohorts using the UCSC LiftOver tool. A small proportion of SNPs (∼0.3 million out of 8.5 million SNPs with MAF ≥ 0.01 in the UK Biobank) could not be successfully converted from GRCh37 to GRCh38 coordinates. To ensure a fair comparison, we repeated the Manc-COJO and Sanc-COJO analyses using the approximately ∼1-million-sample version of the summary statistics (∼0.9 million EUR plus 0.09 million AFR or EUR) as described above, but retained only SNPs that were present in both the UK Biobank and All of Us cohorts at the start of the analysis. For phenotype generation in the All of Us cohort, we extracted lipid traits and applied adjustments as detailed in the **Supplementary Methods**. We retained European ancestry samples (∼80,000 per trait) and African ancestry samples (∼25,000 per trait), with ancestry defined according to the file:

“gs://fc-aou-datasets-controlled/v8/wgs/short_read/snpindel/aux/ancestry/ancestry_preds.tsv” Samples were divided into groups of 1,000, and we applied the same protocol used in the UK Biobank to compute PGS.

## Data availability

UK Biobank and All of Us data are available through formal application to the UK Biobank (https://www.ukbiobank.ac.uk) and the All of Us Research Hub

(https://www.researchallofus.org), respectively. GWAS summary statistics for the four lipid traits analysed in this study are publicly available from the GLGC

(https://csg.sph.umich.edu/willer/public/glgc-lipids2021/). The 1000G Project genotype data used for linkage disequilibrium reference comparisons are available at https://figshare.com/articles/dataset/HapMap3-1KG_1000_Genomes_processed_genotypes_in_PLINK_bed_bim_fam_format/20802700.

## Code availability

Manc-COJO is implemented as a publicly available software package. The C++ source code and precompiled executable binaries are available at https://github.com/light156/multi-ancestry-COJO. Usage instructions and full documentation are provided at https://light156.github.io/multi-ancestry-COJO-docs/.

## Acknowledgement

This research was supported by the Michael Davys Trust at the University of Oxford, the Pioneer Center for Statistical and computational Methods for Advanced Research to Transform Biomedicine (SMARTbiomed), Danish National Research Foundation grant number P4. This study used data from the UK Biobank (project ID: 116122), the All of Us Research Program (workspace name: aou-rw-3508e912), and the 1000 Genomes Project. We thank the participants and investigators of these projects for making this research possible. We thank Dr Tian Lin, Dr Ang Li and Dr Siqi Wang for testing the software on two independent computing platforms across multiple traits to ensure its robustness.

## Authors Contributions

L.Y. and P.M.V. conceptualised the study. N.R.W. and L.Y. jointly supervised the research. X.W. derived the theoretical framework, developed the algorithms, and conducted the data analyses under the supervision of N.R.W and L.Y. Y.W. implemented the C++ software, and optimised the algorithms and analyses in collaboration with X.W. X.W. drafted the manuscript with contributions from Y.W., N.R.W., L.Y., and P.M.V. All authors reviewed and approved the final manuscript.

---

**Extended Figure 1.**
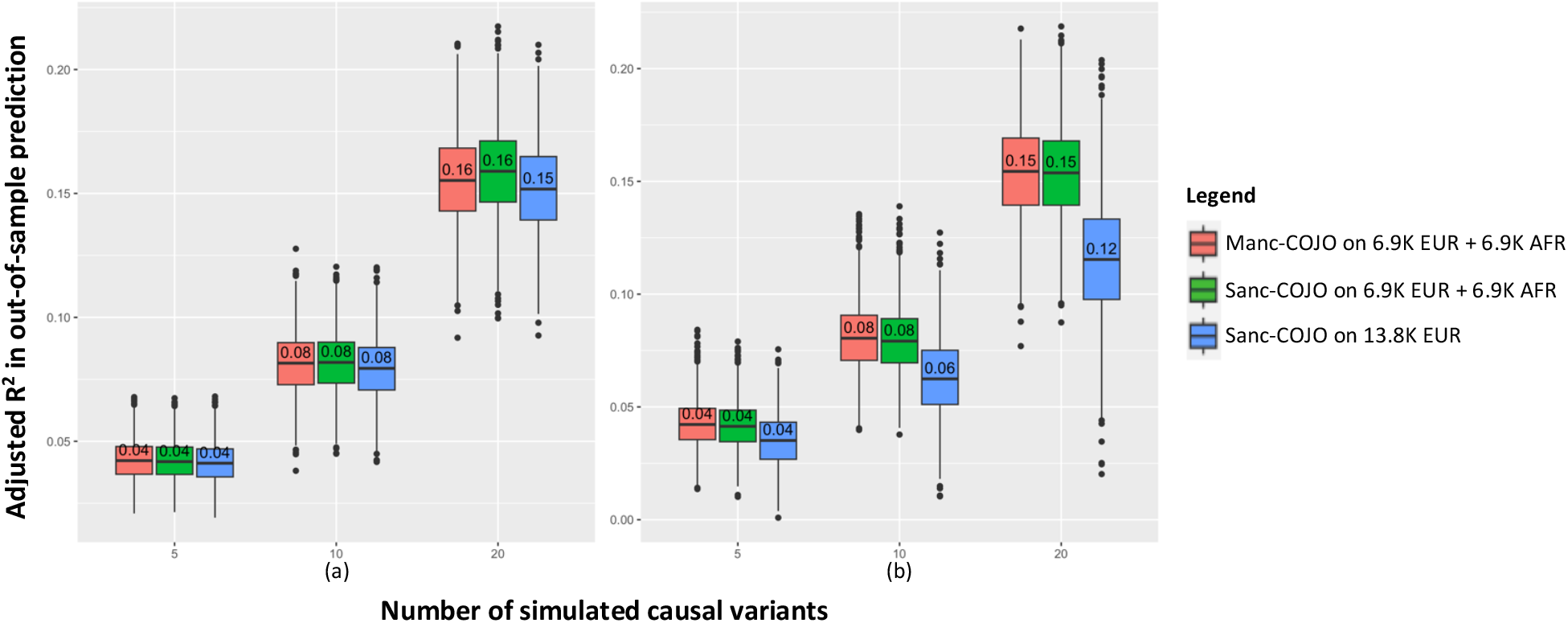
Application of Sanc-COJO to single-ancestry simulation yields a model with good out-of-sample prediction within the same ancestry but relatively low cross-ancestry portability in simulations. This figure summarises simulation results from 5,700 scenarios, illustrating the out-of-sample prediction performance of different COJO methods. SNP effect sizes were generated under the simulation settings presented in **Figure 2** and used to predict outcomes in an independent European cohort (*N* = 13,800) and an African cohort (*N* = 2,100). The x-axis represents the number of simulated causal SNPs per independent LD block. (a) The y-axis indicates the adjusted *R*^2^ when predicting into the European cohort. (b) The y-axis indicates the adjusted *R*^2^ when predicting into the African cohort. When Sanc-COJO is applied to single-ancestry data, the resulting model demonstrates out-of-sample prediction accuracy into an independent sample similar to Manc-COJO, yet it exhibits limited portability across ancestries. Each box plot summarizes 1,900 simulation replicates (38 LD-independent blocks × 50 within-block replicates) and shows the minimum, maximum, mean, and interquartile range, with the numbers indicating the mean.

**Extended Figure 2.**
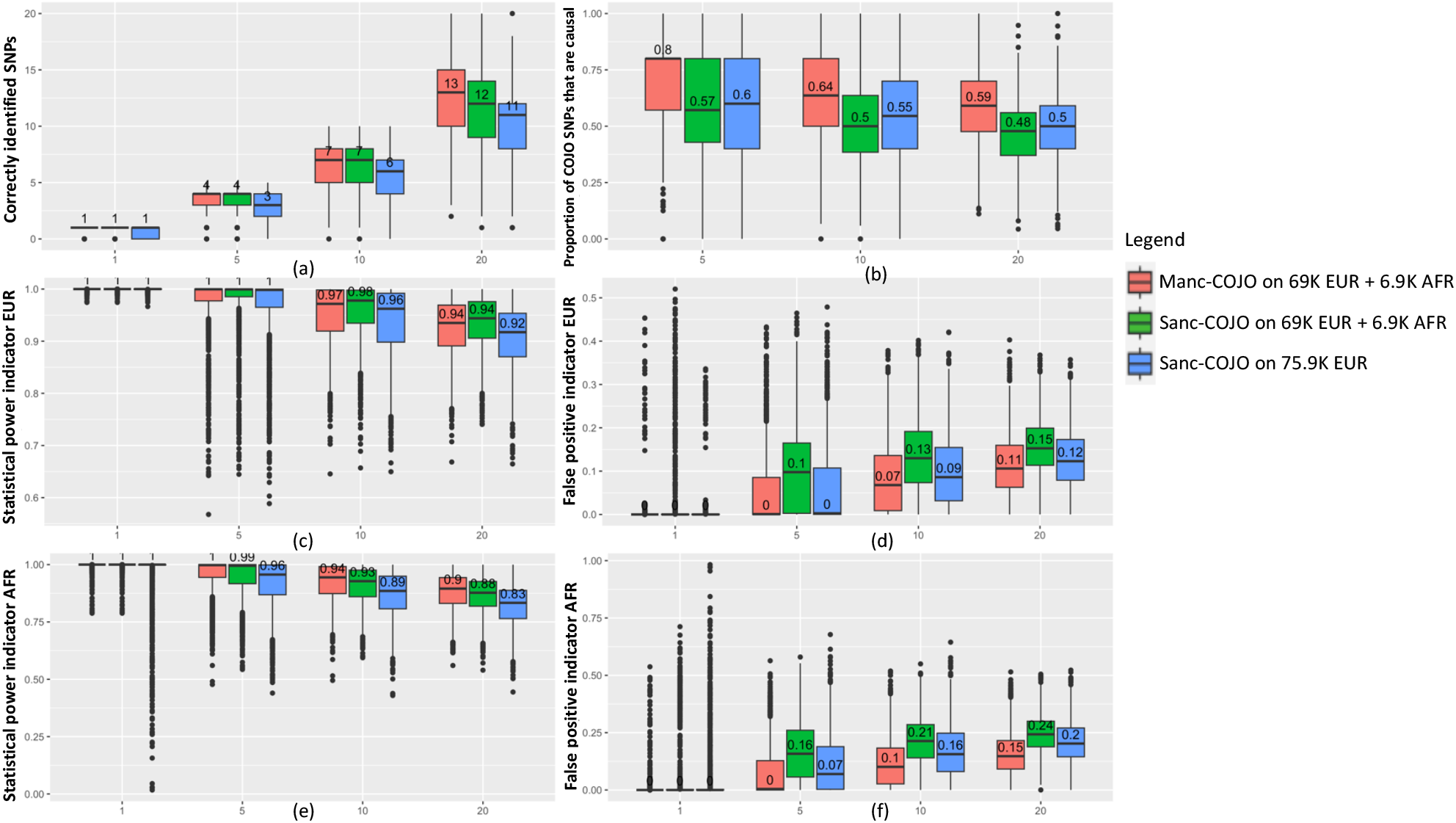
Manc-COJO demonstrates superior performance in simulations with uneven sample sizes across ancestries. This figure summarises simulation results from 7,600 scenarios. We applied our algorithm to a simulated European cohort (*N* = 69,010) and a simulated African cohort (*N* = 6,901), comparing these results with Sanc-COJO applied to either 75,911 Europeans or a meta-analysis of 69,010 Europeans and 6,901 Africans. In-sample LD was used in all scenarios. The x-axis represents the number of simulated causal SNPs. (a) The y-axis indicates the number of causal SNPs correctly identified by each algorithm. (b) The y-axis represents the proportion of identified SNPs that are causal. (c) As COJO algorithms may identify SNPs (termed COJO-SNPs) near the causal SNP rather than the causal SNP itself, we identified the COJO-SNP with the highest LD correlation (in the European cohort) to each causal SNP and averaged these correlations across all causal SNPs; higher values reflect greater power in detecting causal variants. (d) For each COJO SNP, we identified the nearest true causal SNP, calculated their (European) LD, and then averaged these LD values across all COJO SNPs. We then subtracted this mean from 1 to obtain an indicator of false-positive discoveries (*i*.*e*., when selected SNPs are not in LD with causal SNPs, the average LD will be low, yielding a higher false-positive indicator). (e)(f) Power and false-positive indicators as in (c) and (d), respectively, but calculated using African LD. Numbers in each plot represent the median value for each group. Each box plot summarizes 1,900 simulation replicates (38 LD-independent blocks × 50 within-block replicates) and shows the minimum, maximum, mean, and interquartile range, with the numbers indicating the mean. This figure illustrates that Manc-COJO, on average, identifies a greater number of causal SNPs, exhibits superior power, and produces fewer false-positive discoveries when sample sizes across ancestries are unequal.

**Extended Figure 3.**
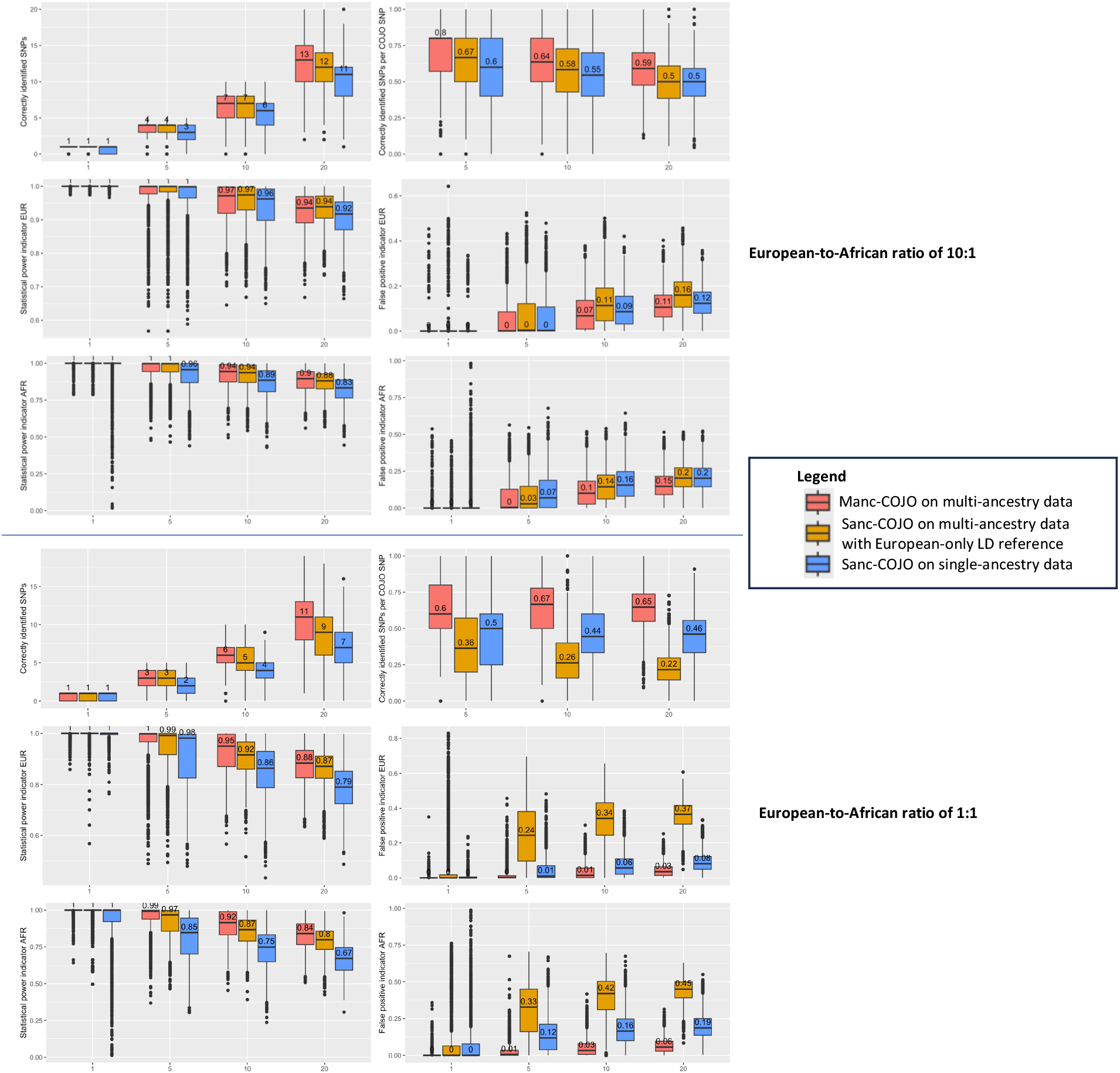
Effectiveness of European-only LD reference panels depends on ancestry composition of the study sample in simulations. This figure shows the performance of Sanc-COJO applied to multi-ancestry GWAS summary statistics (Sanc-COJO->Manc) using a European-only LD reference panel, under two different sample size scenarios: a European:African ratio of 10:1 (top panel) and 1:1 (bottom panel). Model performance is evaluated as in **Figure 2**. When Europeans dominate the sample (10:1 ratio), using a European-only LD reference performs reasonably well, achieving higher power in both ancestries and lower false positive rates in African samples, compared to Sanc-COJO applied to European-only summary statistics. However, when the ancestry balance is equal (1:1 ratio), the use of a European-only LD reference becomes ineffective, leading to marked inflation in false-positive discoveries, particularly in African samples. Each box plot summarizes 1,900 simulation replicates (38 LD-independent blocks × 50 within-block replicates) and shows the minimum, maximum, mean, and interquartile range, with the numbers indicating the mean.

**Extended Figure 4.**
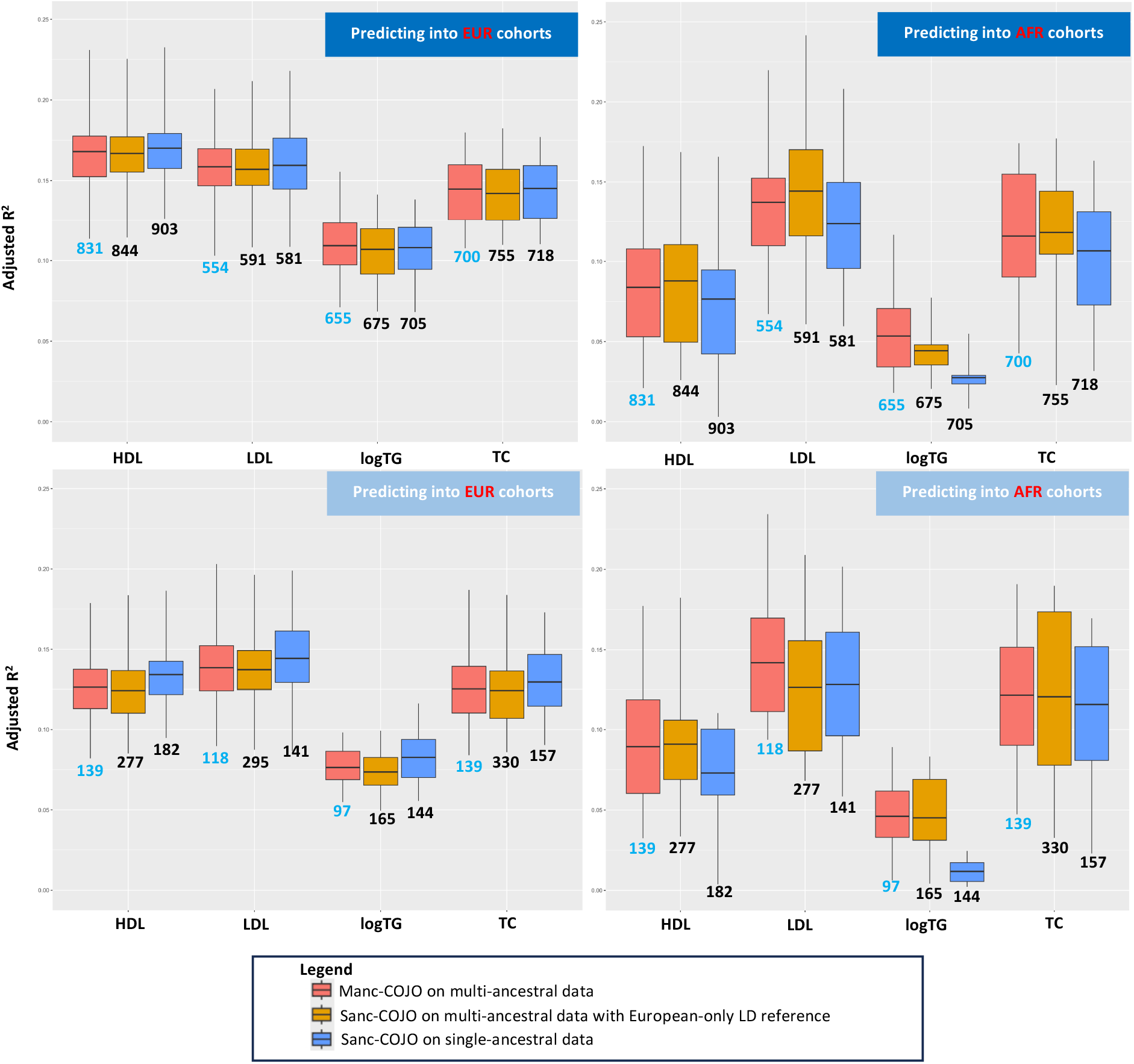
Applying Sanc-COJO to meta-analyses of multi-ancestry GWAS using European-only LD reference panels is only effective when European populations dominate the sample. Published studies have applied Sanc-COJO to multi-ancestry GWAS meta-analyses using European-only LD reference panels, particularly when European individuals comprised the majority of the sample. In this figure, we apply Sanc-COJO using a European-only LD reference under two scenarios: a European:African (EUR:AFR) sample size ratio of 10:1 (top row) and 1:1 (bottom row), predicting into both an independent European cohort (left column) and an African cohort (right column). The results mirrored our simulation findings. When Europeans dominated the sample (10:1), the approach performed reasonably well. However, when the ancestry balance shifted to 1:1, the strategy became ineffective. The number of SNPs selected by Sanc-COJO increased markedly, indicating a higher rate of false-positive inclusions and reduced model parsimony. Each box plot shows the distribution of the out-of-sample prediction R^2^ values for all groups (50 in EUR and 20 in AFR) and displays the minimum, maximum, mean, and interquartile range.

**Extended Figure 5.**
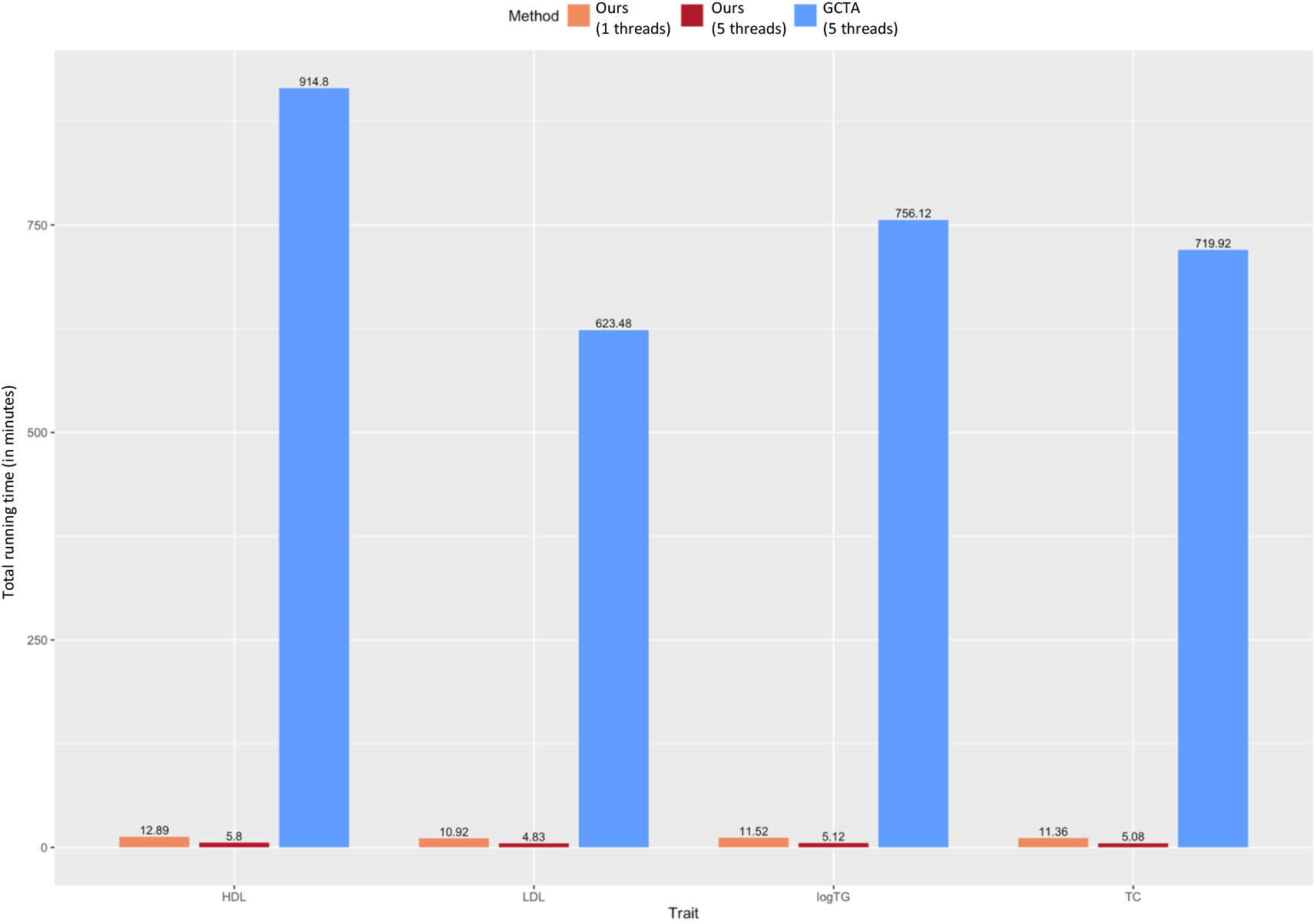
Improved computational efficiency of our software for running Sanc-COJO compared with the implementation in GCTA. Although our software is primarily designed for multi-ancestry analysis, it also supports single-ancestry COJO analyses as in GCTA. The x-axis shows the traits, and the y-axis shows the total runtime across all 22 chromosomes for single-ancestry analysis in ∼1 million EUR individuals (∼6.5 million SNPs, as described in the real-data analysis), using ∼75,911 unrelated UKB EUR samples as the LD reference. Even using a single CPU, the average per-chromosome runtime is below one minute—orders of magnitude faster than the Sanc-COJO algorithm currently implemented in GCTA.

