## Supplementary Documents for "Multi-ancestry conditional and joint analysis (Manc-COJO) applied to GWAS summary statistics"

### Supplementary Methods 1. Important considerations in Manc-COJO model

In this section, we first review the assumptions underlying the COJO models. We then detail our algorithms and outline the step-by-step model selection process. Finally, we demonstrate the validity of the algorithms implemented in our method through theoretical derivations.

#### 1.1 Concepts and assumptions

In GWAS and post-GWAS analyses, there are three SNP effect size concepts: the causal effect size ($\beta_c$), the marginal effect size ($b_m$), and the effect size from the joint analysis of SNPs ($\beta_J$).

##### 1.1.1 Modelling true effect in an idealised population setting

We denote $\beta_c$ as the causal effect size on the phenotype.

We make three assumptions about causal effects:

(1) $\beta_c$ of SNPs act on the phenotype additively, which means that for a specific SNP, each copy of the alternative allele adds, on average, $\beta_c$ to the phenotype.

(2) $\beta_c$ of different SNPs are independent, which means the effect of a specific SNP will not affect the effect of another SNP.

(3) The number of causal variants, and their effect sizes on **standardised** phenotypes, are identical in different ancestries. In other words, under an additive model, if a variant is causal in one ancestry, it is also causal in another ancestry. However, allele frequencies of causal SNPs can differ between ancestries because of genetic drift, bottleneck or differential selective pressure[1]). We later relax this assumption by including ancestry-specific variants.

Our data generation model assumes that phenotypes of **all** individuals in a population (size *N*) depend on causal SNP effects according to the following equation:

$$\mathbf{y} = \mathbf{X}_C \boldsymbol{\beta}_C + \mathbf{e}_C \qquad (1)$$

where

- $\mathbf{y}$ is an *N*x1 vector of **standardised** residuals of phenotypes (after regressing out all covariates)
- $\mathbf{X}_C$ is an *N*x*C* **mean-centred** matrix of *C* genotypes of causal variants shared across ancestries (mean-centered within each ancestry for multi-ancestry datasets)
- $\boldsymbol{\beta}_C$ is a *C*x1 vector of causal effects
- $\mathbf{e}_C$ is an *N*x1 vector of error terms, which is normally distributed under N(0,$\sigma_e^2$)

Importantly, causal variants and their effect sizes are typically unknown for several reasons. For example, not all causal SNPs are genotyped nor imputed from a SNP-array. Even with whole genome sequencing data, there are still gaps (for example, regions near the centromere and telomere[2]). LD adds complexity in identifying the causal variants, especially when there are multiple causal variants within a region of correlated SNPs. In following sections, we will model what is observable in practice, and link it to the unobservable causal effects.

##### 1.1.2 Modeling GWAS marginal effects in an idealised population setting

In GWAS analysis, one SNP at a time is tested for association with the phenotype. Here, we commence the derivation by assuming access to genetic data for **every individual** in the population; accordingly, all parameters are treated as **population parameters**.

Let $\mathbf{x}_m$ be an *N*x1 mean-centred vector of genotypes for the SNP *m* to be tested in the GWAS analysis, and $b_m$ the marginal effect size of the SNP $m$.

As the genotype and error are independent (*i.e.* $\mathbf{x}_m'\mathbf{e}_C = 0$), $\mathbf{x}_m'\mathbf{y} = \mathbf{x}_m'\mathbf{X}_C\boldsymbol{\beta}_C$.
It follows that,

$$b_m = (\mathbf{x}_m'\mathbf{x}_m)^{-1}\mathbf{x}_m'\mathbf{y} \qquad (2)$$

Notably, this expression is derived under an idealised population assumption, and we discuss the estimation of $b_m$ in Section 1.2.

Now, let $p_m$ to be the minor allele frequency of the SNP $m$, $q_m = 1 - p_m$. Based on the definition of binomial variance, and under the Hardy-Weinberg equilibrium (HWE), $\mathbf{x}_m'\mathbf{x}_m$ = $2p_mq_mN$. Let us denote $D_m = 2p_mq_mN$.

Applying equation (2) into equation (1), we get

$$b_m = \frac{\mathbf{x}_m'\mathbf{X}_C\boldsymbol{\beta}_C}{D_m} = \sum_{c=1}^{C} r_{m,c}\beta_c\sqrt{\frac{p_cq_c}{p_mq_m}} \qquad (3)$$

where, $r_{m,c}$ denotes the (signed) LD correlation between SNP *m* and the *c*-th causal SNP. In other words, the value of true marginal effect size ($b_m$) for the SNP $m$ in the GWAS analysis, can be viewed as a weighted sum of true effect size of all causal SNPs, where the weights are the LD correlations between the SNP $m$ and causal SNPs, adjusted for minor allele frequencies.

##### 1.1.3 Modelling joint effects in an idealised population setting

In a joint analysis, any set of *J* SNPs can be fitted together, yielding the corresponding true joint effects (assuming we have genetic data for every individual in the population). The major benefit of considering SNPs jointly is to account for the LD correlation between these SNPs. Joint effects of *J* SNPs can be related to the true marginal effects of those *J* SNPs and the LD correlation between those *J* SNPs:

$$\boldsymbol{\beta}_J = \mathbf{D}_J^{-\frac{1}{2}}\mathbf{R}_J^{-1}\,\mathbf{D}_J^{\frac{1}{2}}\mathbf{b}_M \qquad (4)$$

Here,

- $\mathbf{D}_J$ is a *JxJ* diagonal matrix with *j*-th elements being $D_{j,j} = 2p_jq_jN$
- $\boldsymbol{\beta}_J$ is a *Jx1* vector of true **joint** effect sizes when fitting those *J* SNPs jointly in a model
- $\mathbf{b}_M$ is a *Jx1* vector of true **marginal** effective sizes

- $\mathbf{R}_J$ is a *JxJ* true LD correlation matrix between *J* SNPs that are fitted jointly in the model

Applying equation (3) into equation (4), gives

$$\boldsymbol{\beta}_J = \mathbf{D}_J^{-\frac{1}{2}}\mathbf{R}_J^{-1}\mathbf{r}_{J,C}\mathbf{D}_C^{\frac{1}{2}}\boldsymbol{\beta}_C \qquad (5)$$

where,

- $\mathbf{D}_J$ is a *JxJ* diagonal matrix with *j*-th elements being $D_{j,j} = 2p_jq_jN$
- $\mathbf{r}_{J,C}$ is a *JxC* true LD correlation matrix between *J* SNPs that are fitted jointly in the model and *C* causal SNPs
- $\mathbf{D}_C$ is a *CxC* diagonal matrix with *c*-th elements being $D_{c,c} = 2p_cq_cN$
- $\boldsymbol{\beta}_C$ is a *Cx1* vector of **true** effect sizes of *C* causal SNPs

Now, let

- $\mathbf{V}_J$ be a *JxJ* diagonal matrix with *j*-th elements being $V_{j,j} = 2p_jq_j$
- $\mathbf{V}_C$ be a *CxC* diagonal matrix with *c*-th elements being $V_{c,c} = 2p_cq_c$

Assuming an idealised population, Equation (5) can be simplified to:

$$\boldsymbol{\beta}_J = \mathbf{V}_J^{-\frac{1}{2}}\mathbf{R}_J^{-1}\mathbf{r}_{J,C}\mathbf{V}_C^{\frac{1}{2}}\boldsymbol{\beta}_C \qquad (6)$$

When all C causal SNPs are fitted in a model, $\mathbf{R}_J = \mathbf{r}_{J,C}$, $\mathbf{V}_J = \mathbf{V}_C$. It follows that $\boldsymbol{\beta}_J = \boldsymbol{\beta}_C$**.** In other words, when all causal SNPs are included jointly in a model and all the assumptions described above are satisfied, the true joint SNP effects will equal the causal SNP effects.

##### 1.1.4 Modelling conditional effects in an idealised population setting

Having modelled the marginal and joint effects, we can now define an additional quantity: the conditional effect. Given a set of SNPs (set S), we fit these SNPs jointly into a model as in (4), and obtain the residual:

$$\mathbf{y}_S = \mathbf{y} - \mathbf{X}_S\boldsymbol{\beta}_S$$

For each SNP i not included in the model, the conditional effect of SNP i given set S, denoted $b_{i|\boldsymbol{\beta}_S}$, is essentially the marginal effect of SNP *i* in the following regression:

$$\mathbf{y}_S = \mathbf{x}_i b_{i|\boldsymbol{\beta}_S} + \mathbf{e}_S$$

Intuitively, the conditional effect represents the remaining contribution of SNP *i* to the phenotype after adjusting for the effects of SNPs in set S.

The conditional effect $b_{i|\boldsymbol{\beta}_S}$ can be expressed as follows:

$$\begin{aligned} b_{i|\boldsymbol{\beta}_S} &= b_i - V_i^{-\frac{1}{2}}\mathbf{r}_{i,S}\mathbf{V}_S^{\frac{1}{2}}\boldsymbol{\beta}_S \\ &= b_i - V_i^{-\frac{1}{2}}\mathbf{r}_{i,S}\mathbf{R}_S^{-1}\mathbf{V}_S^{\frac{1}{2}}\mathbf{b}_S \qquad (7) \end{aligned}$$

where,

- $\mathbf{r}_{i,S}$ is a 1 x *S* vector of LD correlations between SNP *i* and each of *S* SNPs that are already selected into the model (set S)
- $\boldsymbol{\beta}_S$ is an *S* x 1 vector of true **joint effects** of the *S* SNPs in set *S* in the model
- $\mathbf{R}_S^{-1}$ is an *S* x *S* matrix of LD correlations between S SNPs that are already in the model

- $\mathbf{b}_S$ is an $S$ x 1 vector of true **marginal effects** of $S$ SNPs in the model
- $b_i$ is the true marginal effect of the SNP *i*
- $\mathbf{V}_S$ is an $S$ x $S$ diagonal matrix with *s*-th elements $V_s = 2p_s q_s$ (Assuming HWE)
- $V_i = 2p_i q_i$ (Assuming HWE)

##### 1.1.5 Assumption of a homogenous LD structure

It is important to note that all derivations above implicitly assume a homogeneous LD structure within the population. Consider a scenario in which a population (denoted by the subscript *com*) is not homogeneous, but consists of two sub-populations (denoted by subscripts *1* and *2*) **com**bined together. Each sub-population is internally homogeneous, yet they differ in allele frequencies at the same SNPs, resulting in distinct correlation structures between the sub-populations. In such scenario, the genotypic variance of SNP *i* in the combined population ($V_{com,i}$) will no longer be $2p_{com,i}q_{com,i}$, but rather

$$V_{com,i} = 2p_{1,i}q_{1,i}w_1 + 2p_{2,\mathrm{i}}q_{2,\mathrm{i}}w_2 + 4w_1w_2(p_{1,\mathrm{i}} - p_{2,\mathrm{i}})^2 \qquad (8)$$

where $w_1$ and $w_2$ are the proportion of sub-population 1 and 2 in the combined population, respectively. In other words, the Hardy–Weinberg equilibrium (HWE) assumption is violated when the population is not homogeneous.

Let $V_{a,i}$ be the genotypic variance of SNP *i* in the population *a*, where $V_{a,i} = 2p_{a,i}q_{a,i}$, and $r_{\mathrm{a},i,k}$ be the (signed) LD correlation between SNP *i* and SNP *k*. The LD correlation between SNP *i* and SNP *k* in the combined population, can be expressed as:

$$r_{\mathrm{com},i,k} = w_1 r_{1,\mathrm{i},k}\sqrt{\frac{V_{1,i}V_{1,k}}{V_{\mathrm{com},i}V_{\mathrm{com},k}}} + w_2 r_{2,i,k}\sqrt{\frac{V_{2,i}V_{2,k}}{V_{\mathrm{com},i}V_{\mathrm{com},k}}} + \frac{4w_1w_2(p_{1,\mathrm{i}}-p_{2,\mathrm{i}})(p_{1,\mathrm{k}}-p_{2,\mathrm{k}})}{\sqrt{V_{\mathrm{com},i}V_{\mathrm{com},k}}} \qquad (9)$$

It is clear that $r_{\mathrm{com},i,k} = r_{1,i,k}$ will hold for all SNP pairs *i* and *k* only in a homogeneous population (*i.e.* $w_1 = 1, w_2 = 0$, $V_{1,i} = V_{\mathrm{com},i}$ , $V_{1,k} = V_{\mathrm{com},k}$, $p_{1,\mathrm{i}} = p_{2,\mathrm{i}}$, and $p_{1,\mathrm{k}} = p_{2,\mathrm{k}}$)

#### 1.2 Addressing the challenges of real-world data

In an idealised scenario, where genetic data are available for all individuals in a population, the population is homogeneous, and all model assumptions hold, fitting all causal SNPs jointly would recover the causal effect sizes. In practice, such conditions never exist, and some assumption violations are beyond our control. Therefore, the aim of our algorithm (and, arguably, all COJO- or fine-mapping–type algorithms) is to identify a set of SNPs that approximates the causal variants—in both number and genomic location—while controlling for manageable sources of error and recognising inherent limitations of the model. To achieve this aim, we can implement an extended version of conditional and joint analyses in each iteration of the algorithm. As in Sanc-COJO[3], LD correlations between SNPs can be estimated from a reference sample, with allele frequencies and marginal effect sizes available from GWAS (meta-analysis) summary statistics. However, when using summary statistics several additional quality control procedures are needed to make sure results are comparable to scenarios where indivual level genotype data are available.

##### 1.2.1 Considerations in dealing with real-world data

- *Estimation of GWAS marginal effects*

In practice, it is impossible to obtain genotype data from every individual in a population. Consequently, we analyze a sample drawn from the population and perform an association test on the observed data. It follows from Equation (2) that the ordinary least-squares estimator of the marginal effect $\hat{b}_m$for SNP $m$can be written as

$$\hat{b}_m = (\mathbf{x}_m'\mathbf{x}_m)^{-1}\mathbf{x}_m'\mathbf{y} = b_m + \mathrm{e}_m$$

Here, $\mathbf{y}$ now denotes the $N_m$ x 1 vector of **observed** phenotypes, and $\mathbf{x}_m$ now denotes corresponding $N_m$ x 1 vector of **observed** genotypes at SNP $m$, where $N_m$ is the sample size for SNP m. It follows that,

$$\mathrm{E}[\hat{b}_m] = b_m$$

- *Phenotypic variance*

In our model we assumed phenotypic variance to be 1 across studies. However, this may not be true in practice (even when traits are described as standardised this is often prior to accounting for covariates). Therefore, as in Sanc-COJO, the phenotypic variance in each study is estimated from GWAS summary statistics. For each SNP $i$, the estimate of $\mathbf{y}'\mathbf{y}$ can be estimated from:

$$\hat{\mathbf{y}}'\hat{\mathbf{y}} = \hat{D}_i\hat{S}_i^2(n_i - 1) + \hat{D}_i\hat{b}_i^2$$

where $\hat{S}_i^2$ is the squared standard error of the marginal effect, $\hat{b}_i$ is observed marginal effect, $n_i$ is the per-SNP sample size of SNP i (*i.e.* not the population size), and $\hat{D}_i = 2\hat{p}_i\hat{q}_i$.
Phenotypic variance ($V_P$) can then be estimated by:

$$\hat{V}_P = \hat{D}_i\hat{S}_i^2 + \frac{\hat{D}_i\hat{b}_i^2}{n_i - 1} \qquad (10)$$

In our analyses, we took the median of $\hat{V}_P$ across all SNPs in the GWAS summary statistics as the estimate of the phenotypic variance.

- *Effective sample size*

As discussed in the Sanc-COJO paper, when data sets contributing to the GWAS results include relatives, the reported sample size will be larger than the effective sample size. In our analyses we estimated the per-SNP sample size for any SNP $i$ using the equation[3]:

$$\hat{n}_i = \frac{\hat{V}_P - 2\hat{p}_i\hat{q}_i\hat{b}_i^2}{2\hat{p}_i\hat{q}_i\hat{S}_i^2} + 1 \qquad (11)$$

- *Per-SNP sample size*

In real data sets, per-SNP sample size can be significantly different between SNPs. Within a single data set differences in sample size between SNPs occur because of insufficient SNPs in a region to allow accurate imputation, which in turn may reflect genotype missingness. In GWAS meta-analyses different sub-studies may contribute different SNPs which can generate very large differences in sample sizes between SNPs. We suggest exclusion of SNPs to avoid extreme differences in per-SNP sample sizes, and outline strategies for handling genotype missingness among the remaining SNPs in **Supplementary Note 3**.

- *Choice of reference sample to estimate LD correlation matrix*

In Section 1.1.5, we examined the assumption of a homogeneous LD structure within a GWAS cohort. When GWAS summary statistics are obtained from meta-analyses of multiple cohorts, or when they are combined with an external LD reference cohort (*i.e.,* out-of-sample LD), as in Sanc-COJO, an additional layer of complexity is introduced. For the analysis to yield reliable outcomes, all cohorts contributing to the meta-analysis within a single inferred ancestry should exhibit a broadly similar LD structure (*i.e.,* heterogeneity may exist between GWAS meta-analysis cohorts but not within each cohort). Furthermore, it is essential to ensure that the LD structure of the GWAS cohort and that of the LD reference cohort are sufficiently similar to produce robust results.

Ideally, the LD estimated from the same sample used to generate the GWAS should also be employed in COJO analyses. For this reason, we support sharing of LD matrices together with GWAS summary statistics thereby facilitating information sharing when access to raw data is restricted. When the use of out-of-sample LD is unavoidable, its selection may be informed by prior knowledge or by approaches such as principal component (PC) analysis, which lies beyond the scope of this paper.

In addition, we recommend testing for differences in allele frequency between the reference and GWAS cohorts. When the sample size is large enough, the distribution of allele frequency of SNP *i* can be approximated by a normal distribution. $\hat{p}_i \sim \mathrm{N}\left(p, \frac{pq}{2N}\right)$. Hence, we can consider removing SNPs that are significantly different at specified threshold (for example, P = 5x10$^{-8}$) between the GWAS (subscript *GWAS*) and reference data (subscript *ref*) sets based on the following z-score:

$$Z = \frac{\hat{p}_{ref} - \hat{p}_{GWAS}}{\sqrt{\frac{\hat{p}_{ref}\hat{q}_{ref}}{2n_{ref}} + \frac{\hat{p}_{GWAS}\hat{q}_{GWAS}}{2n_{GWAS}}}} \qquad (12)$$

Another important consideration is the size of the reference sample. Empirical evidence suggest that reference sample sizes > 5,000 will generate relatively accurate LD correlation estimates[3].

##### 1.2.2 Overview of single-ancestry COJO algorithm

In our study, we re-derived the equations used in the original GCTA-COJO (*i.e.* Sanc-COJO) so that the LD correlation matrix could be used as the LD reference, in addition to individual-level genotype data.

In the first iteration of the Sanc-COJO algorithm, the SNP with the smallest p-value is selected into the model. Conditional on this SNP, the conditional z-statistics of all other SNPs (taking SNP *i* as an example) are then calculated as:

$$z_{i|\widehat{\boldsymbol{\beta}}_S} = \frac{\hat{b}_i - \hat{V}_i^{-\frac{1}{2}} \hat{\mathbf{r}}_{i,S} \, \widehat{\mathbf{R}}_S^{-1} \widehat{\mathbf{V}}_S^{\frac{1}{2}} \hat{\mathbf{b}}_S}{\hat{S}_i} \quad (13)$$

where,

- $\hat{\mathbf{r}}_{i,S}$ is a 1 x $S$ vector of **observed** LD correlations between SNP *i* and each of the *S* SNPs already selected into the model (in the first iteration, only 1 SNP is in the model)
- $\widehat{\mathbf{R}}_S$ is an *S* x *S* matrix of **observed** LD correlations between S SNPs that are already in the model (in the first iteration, this reduces to a 1×1 matrix with the value equal to 1)
- $\hat{\mathbf{b}}_S$ is an S x 1 vector of **observed marginal effects** of *S* SNPs in the model
- $\widehat{\boldsymbol{\beta}}_S$ is an S x 1 vector of **observed joint effects** of *S* SNPs in the model
- $\hat{b}_i$ is the **observed** marginal effect of the SNP *i*
- $\widehat{\mathbf{V}}_S$ is an *S* x *S* diagonal matrix with *s*-th elements $\hat{V}_s = 2\hat{p}_s\hat{q}_s$ (assuming HWE)
- $\hat{V}_i = 2\hat{p}_i\hat{q}_i$ (assuming HWE)
- As in GCTA-COJO, to avoid over-fitting, a conservative approach is used to get the conditional p-value, where $\hat{S}_i$ denotes the standard error of the observed marginal effect of SNP *i*.

After getting the z-statistics for each of the remaining SNPs, the SNP with the smallest p-value is selected into the model (*i.e.* the SNP with largest $z^2$). Afterwards, a joint analysis is performed as follows:

$$\widehat{\boldsymbol{\beta}}_{S+1} = \widehat{\mathbf{V}}_{S+1}^{-\frac{1}{2}} \widehat{\mathbf{R}}_{S+1}^{-1} \widehat{\mathbf{V}}_{S+1}^{\frac{1}{2}} \hat{\mathbf{b}}_{S+1} \quad (14)$$

Here, the definition of $\widehat{\boldsymbol{\beta}}_{S+1}$, $\hat{\mathbf{b}}_{S+1}$, $\widehat{\mathbf{R}}_{S+1}$, and $\widehat{\mathbf{V}}_{S+1}$, are the same as defined in equation (13), except that now there are *S*+1 SNPs in the model. As in Sanc-COJO, the estimation error of joint effects of *S*+1 SNPs can be expressed as:

$$\text{Var}(\widehat{\boldsymbol{\beta}}_{S+1}) = \hat{\sigma}_J^2 \widehat{\mathbf{D}}_{S+1}^{-\frac{1}{2}} \widehat{\mathbf{R}}_{S+1}^{-1} \widehat{\mathbf{D}}_{S+1}^{-\frac{1}{2}} \quad (15)$$

Where $\widehat{\mathbf{D}}_{S+1}$ is an (*S*+1)x(*S*+1) diagonal matrix with *s*-th elements $\widehat{D}_s = 2\hat{p}_s\hat{q}_s\hat{n}_s$, and $\hat{n}_s$ is the effect sample sizes of SNP *s* (as defined in equation 11). Here, the residual variance of the joint analysis, can be estimated with the equation:

$$\hat{\sigma}_J^2 = \hat{V}_P - \hat{\mathbf{b}}_{S+1}{}' \widehat{\mathbf{V}}_{S+1} \widehat{\boldsymbol{\beta}}_{S+1}$$

The vector of z-scores for the joint SNP effects for these *S+1* SNPs is given by

$$\mathbf{z}_{S+1} = \text{diag}(\text{Var}(\widehat{\boldsymbol{\beta}}_{S+1}))^{-\frac{1}{2}} \widehat{\boldsymbol{\beta}}_{S+1} \quad (16)$$

In this single ancestry COJO algorithm, in each iteration, SNPs whose joint p-value fall above certain significance threshold are removed. The above conditional and joint analysis steps are repeated until no SNP can be removed or added. In our analyses we set the significance threshold for selection as p-value < $5 \times 10^{-8}$.

##### 1.2.3 Extending COJO analysis into multi-ancestry settings

The single-ancestry COJO algorithm outlined in the previous section can be extended into a multi-ancestry setting. By leveraging diverse LD structures from different ancestries, we expect to gain better resolution in tagging causal variants (shared across ancestries), and hence in estimating the number of independent associations.

- *Overview of the multi-ancestry COJO algorithm*

The multi-ancestry COJO algorithm assumes GWAS summary statistics of two or more ancestries which have been processed through quality control steps described in section 1.2.1. The multi-ancestry COJO algorithm is summarised in 8 steps and we have implemented it in the new Manc-COJO software. The software requires a predefined genome-wide significance threshold (we used $5 \times 10^{-8}$) and a pre-set collinearity threshold between SNPs (*i.e.* maximum LD $r^2$, we used 0.9).

**Step 1:** Conduct an inverse-variance meta-analysis (IVM) of GWAS summary statistics across all ancestries. The SNP with the smallest meta-analysis p-value is selected for inclusion in the model.

**Step 2:** SNPs above the pre-set collinearity threshold with any SNPs already included in the model in any ancestry are removed in all ancestries.

**Step 3:** Within each ancestry, a conditional analysis is performed and the conditional effects of each SNP are calculated using Equation (13).

**Step 4:** An IVM of the conditional effects of each SNP is then conducted and the SNP with the smallest meta-analysis p-value is selected for inclusion in the model.

**Step 5:** Subsequently, a joint analysis is conducted within each ancestry using Equation (14).

**Step 6:** An IVM of the joint effects of SNPs included in the model is performed, and SNPs whose meta-analysis p-value for joint effects exceeds the pre-set threshold are removed from the model.

**Step 7:** Steps 2 through 6 are repeated until no SNP can be added to or removed from the model.

**Step 8:** To detect ancestry-specific associations, single-ancestry COJO is applied to each individual ancestry GWAS using models in which all Manc-COJO SNPs are included prior to the iterative addition or removal of SNPs. The Manc-COJO SNPs are fixed within the model and are not subject to exclusion during subsequent variable selection iterations.

- *Optional $R^2$ parameter*

In variable selection algorithms, in addition to the approach implemented in the Sanc-COJO model, another commonly used strategy is to consider changes in the adjusted $R^2$ (Coefficient of determination) of model. Using summary statistics and an LD reference, the $R^2$ of the multiple linear regression can be approximated as:

$$\hat{R}^2 = \frac{\hat{\mathbf{b}}_S{}' \hat{\mathbf{V}}_S \hat{\boldsymbol{\beta}}_S}{\hat{V}_P}$$

where $\hat{\mathbf{b}}_S, \hat{\mathbf{V}}_S, \hat{\boldsymbol{\beta}}_S$ are as defined above. The estimated adjusted $R^2$ of the model can then be calculated as:

$$\hat{R}^2_{adj} = 1 - \frac{(1 - \hat{R}^2)(\hat{n} - 1)}{\hat{n} - s - 1}$$

Where $s$ is the number of SNPs in the model, $\hat{n}$ is the effective sample size as defined in equation (11).

In the Manc-COJO algorithm, it is possible to use changes in $\hat{R}^2_{adj}$ as an *optional* selection criterion for researchers who prefer a more conservative model in regions with extremely complex LD structure. Researchers can specify thresholds that require the $\hat{R}^2_{adj}$ of the model to increase by a certain percentage during both forward and backward selection in *both* ancestries for a SNP to be included in the model.

For example, setting the forward-selection $\hat{R}^2_{adj}$ threshold to 0.1 means that, in each iteration, after including a SNP, the $\hat{R}^2_{adj}$ of the model must be at least 1.1 times the value from the previous iteration in both ancestries. Conversely, setting the backward-selection threshold to −0.1 means that, if adding a new SNP causes other SNPs to become insignificant and be dropped, then after excluding those SNPs, the $\hat{R}^2_{adj}$ of the resulting model must remain at least 0.9 times the value from the previous iteration for the new SNP to be retained. Otherwise, the algorithm reverts to the model from the last iteration and continues screening other SNPs for inclusion. By default, the $\hat{R}^2_{adj}$ thresholds for both forward and backward selection are set to −1, meaning that the selection criterion based on $\hat{R}^2_{adj}$ is not applied.

- *The benefit of multi-ancestry analyses*

By leveraging differences in LD structures across ancestries, Manc-COJO may enhance the ability to detect independent associations. To illustrate this, we begin with the single-ancestry analysis.

Under the infinitesimal model, and given the standardised phenotypic variance, the estimation error in GWAS for SNP $m$ ($\hat{S}_m$) can be approximated by $\frac{1}{\sqrt{2\hat{p}_m\hat{q}_m\hat{n}_m}}$. In GWAS analyses, a common simplifying assumption is that the genotype matrix is fixed, even though it typically represents only a sample of the population. Consequently, we can approximate $\frac{1}{\sqrt{2\hat{p}_m\hat{q}_m\hat{n}_m}} \approx \frac{1}{\sqrt{2p_m q_m n_m}}$ (Note, this assumption again underscores the importance of selecting an appropriate LD reference, as discussed previously, and highlights the need to understand the limitations of each model arising from its underlying assumptions.).

Therefore for SNP $m$, the association z-score in GWAS is,

$$z_m = \frac{\hat{b}_m}{\hat{S}_m} = \frac{b_m + e_m}{\hat{S}_m} = e_m\sqrt{2p_m q_m n_m} + \sum_{c=1}^{C} r_{m,c}\beta_c\sqrt{2p_c q_c n_m}$$
$$= \sum_{c=1}^{C} r_{m,c}\beta_c\sqrt{2p_c q_c n_m} + \varepsilon_m \qquad (17)$$

where $\varepsilon_m = e_m\sqrt{2p_m q_m n_m}$, and it follows that $\varepsilon_m \sim \mathrm{N}(0,1)$

When there is only one causal SNP within an LD block, equation (17) can be simplified to

$$z_m = r_{m,c}\beta_c\sqrt{2p_c q_c n_m} + \varepsilon_m \qquad (18)$$

where $\beta_c\sqrt{2p_c q_c}$ is a constant across all SNPs (note $r_{m,c}$, $\beta_c$, $p_c$, and $q_c$ here are population parameters). In other words, in the absence of genotype missingness—so that the sample sizes of all SNPs are **equal**—and when causal SNP effect sizes are **large**, it is relatively straightforward to identify the SNP closest to the causal SNP (*i.e.,* the SNP with the highest $|r_{m,c}|$). However, even in this idealised scenario, if per-SNP sample sizes differ due to genotype missingness, or if there are multiple tightly linked variants, the causal SNP may not correspond to the largest $|z_m|$, even when it is genotyped.

When there is more than one causal SNP in a region of correlated SNPs, it is evident from equation (4) that the most significant SNP may not be the SNP closest to causal SNPs owing to effects such as genetic masking[4] (which refers to when trait increasing alleles are present on different haplotype backgrounds). Recognising that genomic regions are likely to habour multiple causal SNPs motivates the joint analysis approach.

In multi-ancestry meta-analysis, the z-score of SNP *i* ($z_{i,manc}$), derived from an inverse-variance meta-analysis of summary statistics from each ancestry, can be expressed as follows:

$$z_{i,manc} = \sum_{anc}\sqrt{\frac{D_{i,anc}}{D}}\, r_{i,c,anc}\beta_c\sqrt{D_{c,anc}} + e \qquad (19)$$

Here,

- $z_{i,manc}$ is the z-score of the SNP *i* in the multi-ancestry analysis IVM
- $\mathrm{D_{i,anc}} = 2p_{i,anc}q_{i,anc}n_{anc}$, where $p_{i,anc}$ is the minor allele frequency (MAF) of the SNP *i* in a specific ancestry
- $D = \sum_{anc} D_{i,anc}$
- $r_{i,c,anc}$ is the LD correlation between SNP *i* and the causal SNP *c* in a specific ancestry
- $\beta_c$ is the causal effect, which is assumed to be identical across ancestries
- Assuming the estimation error of effect sizes are independent across ancestries, it follows $e \sim \mathrm{N}(0,1)$

Note that in the equation (19), $p_{i,anc}$, $r_{i,c,anc}$, $q_{i,anc}$, and $\beta_c$ are all population parameters. When there are no genotype missingness, $\beta_c\sqrt{D_{c,anc}}$ is a constant for an ancestry. Assuming causal variants are shared across ancestries, $r_{i,c,anc}$ will reach its maximum (*i.e.*, 1) in all ancestries when the SNP *i* is either the causal SNP or in complete LD with causal SNP in all ancestries. Given that LD structures differ across ancestries, extending COJO algorithms across multiple ancestries can narrow the LD window, thereby improving resolution in identifying the causal variants. However, as discussed previously, when per-SNP sample sizes vary, causal effect sizes are small, or causal SNPs are rare (i.e., low MAF, resulting in lower $D_{c,anc}$ values), the SNP selected by the COJO algorithm may not correspond to the SNP closest to the causal variant, due to the influence of the error term.

When there is more than one causal SNP in a region, equation (19) can be further extended to

$$z_{i,manc} = \sum_{anc} \sqrt{\frac{D_{i,anc}}{D}} \sum_{c=1}^{C} r_{i,c,anc} \beta_c \sqrt{D_{c,anc}} + \varepsilon \quad (20)$$

Simulations show that the multi-ancestry COJO algorithm performs better than the single ancestry analysis of the same sample size in terms of finding independent associations.

### Supplementary Methods 2. Simulations

We conducted a series of simulations to compare the multi-ancestry COJO analysis with the single-ancestry COJO analysis of the same sample size with multiple criteria.

#### 2.1 Simulation settings

##### 2.1.1 Simulation cohort

We used real genotype data from chromosome 22 of UK Biobank participants to establish our simulation framework. From all unrelated individuals (coefficient of relationship ≤ 0.05, as calculated using GCTA's default algorithm) of inferred European ancestry (EUR; *N* ~ 340,000) and African ancestry (AFR; *N* = 6,901)—defined based on published ancestry projections[5]—we selected five cohorts. The AFR cohort (AFR1) comprises 6,901 unrelated participants of inferred AFR. EUR cohort 1 (EUR1) and cohort 2 (EUR2) each consist of 6,901 randomly selected, unrelated individuals of inferred EUR, with no overlap or relatedness between EUR1 and EUR2. We then combined EUR1 and EUR2 to form a cohort named EUR1_EUR2 (*N* = 13,802). Additionally, we randomly selected 13,802 unrelated European participants, independent of EUR1_EUR2, for out-of-sample prediction (named EURpred), and selected 2,000 African participants for out-of-sample prediction in African ancestry (named AFRpred), independent of AFR1. The sample sizes of EUR1 and EUR2 were matched to AFR1 because previous studies have shown that combining equal-sized European and African ancestry cohorts maximises power for detecting causal associations in multi-ancestry meta-analyses[6,7].

##### 2.1.2 Partitioning the genome

Previously, Berisa & Pickrell partitioned the genome and defined LD-independent blocks separately for African and European ancestries[11], assuming that LD correlations between SNPs in distinct LD-independent blocks are negligibly different from zero. In this study, we first partitioned chromosome 22 separately for the African and European cohorts (N=33 and N=23 blocks, respectively). The blocks were partially overlapping, and N=52 overlapping sections were extracted (**Supplementary Figure 1**). Within each of these 52 overlapping blocks, we calculated allele frequencies for all SNPs (using PLINK 1.9 --freq) in the AFR1, EUR1, EUR2, EURpred and AFRpred cohorts, retaining only SNPs with MAF>=0.01 across all data sets. Blocks with fewer than

1,000 SNPs remaining after this filtering were excluded, resulting in 38 blocks for use in simulation.

##### 2.1.3 Simulation of causal SNPs

Within each of the 38 selected blocks, we randomly selected 1, 5, 15, or 20 SNPs to be causal variants. Causal effects ($\beta_c$) were simulated to be identical across ancestries and such that the average per-SNP heritability ($h_c^2 = 2p_c q_c \beta_c^2$) of causal SNP ranged between 0.5% and 1.5% in all cohorts. However, heritability values could differ across ancestries owing to varying MAFs for the same SNP. To avoid collinearity, LD $r^2$ between all pairs of SNPs selected to be causal were required to be ≤ 0.8 in both ancestries. For each combination of block and number of causal variants, we repeated the simulation 50 times, using different sets of causal variants and effect sizes in each iteration. This resulted in 7,600 scenarios, accounting for 38 blocks, 4 different numbers of causal SNPs, and 50 variations of causal SNP sets and effect sizes.

##### 2.1.4 Simulation of phenotypes

We simulated phenotypes for individuals within each cohort as in equation (1)**.** Now,

- $\mathbf{y}$ is an *N* x 1 vector of simulated phenotypes.
- $\mathbf{X}_C$ is an *N* x *C* genotype, with entries encoded as 0, 1, or 2, corresponding to the number of alternative alleles at each SNP. Missing values in $\mathbf{X}_C$ are imputed as the mean value of genotypes of that SNP in that specific cohort.
- $\boldsymbol{\beta}_C$ is a *C* x 1 vector of simulated effect sizes as described in the previous section, and $\mathbf{e}_C$ is an *N* x 1 error vector, and each element of the vector is assumed to be drawn from a normal distribution N ~ (0, var($\mathbf{X}_C \boldsymbol{\beta}_C$)*($\frac{1-h^2}{h^2}$)), where, $h^2 = \sum_c 2p_c q_c \beta_c^2$ ($\beta_c$ is *c*-th element of vector $\boldsymbol{\beta}_C$) is the total SNP-based heritability.

We simulated phenotypes within each of the six cohorts (AFR1, EUR1, EUR2, EUR1_EUR2, EURpred, AFRpred).

##### 2.1.5 GWAS simulation, and benchmark against existing method

For each of the 7,600 scenarios, we conducted a GWAS in four cohorts (EUR1, EUR2, EUR1_EUR2 and AFR1) using PLINK 2.0 (--glm hide-covar cols=+a1freq). Subsequently, we performed an inverse-variance meta-analysis of summary statistics from EUR1 and AFR1 using custom *R* code, naming the resulting cohort EUR1_AFR1. We compared outputs across the following GWAS and model combinations: Manc-COJO with EUR1 and AFR1, Sanc-COJO with EUR1_EUR2 and Sanc-COJO->Manc using EUR1_AFR1. Default settings were applied in all scenarios for both Sanc-COJO and Manc-COJO, with in-sample LD used throughout. For the EUR1_AFR1 cohort, in-sample LD was estimated from the merged genotypes of AFR1 and EUR1 using PLINK 1.9 (--bmerge). For Sanc-COJO->Manc, we also applied a European-only LD reference (EUR1_EUR2 in this case, which we named Sanc-COJO->Manc/EURref) in addition to the in-sample mixed-ancestry LD.

##### 2.1.6 Model performance measurement metrics

To assess model performance, we established several criteria to evaluate Manc-COJO outputs. First, to compare Manc-COJO outputs with true effects, we recorded how many COJO-selected

SNPs were simulated as causal SNPs and the proportion of COJO-selected SNPs that were causal. This criterion is justified because the primary objective of Manc-COJO is to construct a model with SNPs that have independent associations. However, in some instances, rather than selecting causal SNPs, nearby SNPs in high LD with the causal SNPs may be included in the final selected set, which still aligns with the goal of the method. To account for such scenario, we calculated two additional measures as indicators for power and false positive discovery: we first build a *CxM* LD correlation matrix, where *C* is the number of causal SNPs, and *M* is the number of COJO SNPs. For each of *M* rows, we pick the highest value, and then calculate the mean of this group of highest values. This can be viewed as an indicator for power, as it measures for each causal SNP the highest LD correlation to COJO SNPs. In an idealised situation, where all causal SNPs have been correctly identified, this number should be 1. Similarly, for each of *M* COJO SNPs, we calculated the mean of the highest values from each column, and subtracted the mean from 1. The resulting value provides an indicator of false-positive discoveries: it equals 0 when all COJO SNPs are causal SNPs and approaches 1 when COJO SNPs are less correlated with the causal SNPs. LD correlation matrices were calculated from both EUR1_EUR2 and AFR1 cohort.

Beyond comparisons with the ground truth, we also assessed out-of-sample prediction accuracies in independent European and African ancestry cohorts using joint SNP-effect sizes of COJO SNPs. Although the COJO algorithm is not designed to maximise out-of-sample risk prediction, comparing out-of-sample prediction using COJO selected SNPs is a criterion that can be applied in real data analysis as well as in simulation. We randomly divided the 13,800 EURpred individuals into 23 groups of 600, calculated polygenic scores for each group ($\mathbf{y}_{\mathrm{Pred}} = \mathbf{X}_{\mathrm{Pred}}\boldsymbol{\beta}_{\mathrm{COJO}}$), and fitted a linear model in R using lm() to obtain the adjusted coefficient of determination ($R^2_{adj}$). We then computed the mean and standard error of these 23 regression coefficients. Similarly, we randomly divided the 2,000 AFRpred individuals into 20 groups of 100, and calculated the adjusted coefficient of determination.

#### 2.2 Simulation sensitivity analysis

Section 2.1 outlined our simulation strategies under idealised conditions, where the ancestry composition maximises power to detect causal associations, all causal SNPs are genotyped, causal effects are identical across ancestries, and the LD structure in the reference panel matches that of the cohort used for GWAS. However, in practice, these assumptions are often violated. To assess the impact of such violations, we conducted a series of simulations in which one assumption was altered at a time.

We first considered a scenario where European samples constitute the majority of the cohort. All simulation settings remained identical to those described in Section 2.1, except that the EUR1 sample size was increased to 69,010. For all simulations in this scenario, in-sample LD reference panels were used. This means that the LD reference for EUR1 was also larger than in Section 2.1. As a result, the input for Manc-COJO was the GWAS summary statistics from EUR1 ($N$ = 69,010) and AFR1 ($N$ = 6,901). For single-ancestry Sanc-COJO, the input was GWAS from

EUR1_EUR2 (*N* = 75,911). For multi-ancestry Sanc-COJO with ancestry-matched LD reference, the input was the meta-analysis of EUR1 and AFR1 GWAS, and the LD reference panel was the merged EUR1 and AFR1 cohort (i.e. EUR1_AFR1). We also examined an additional setting: multi-ancestry Sanc-COJO using a European-only LD reference panel. In this case, the input GWAS was the same meta-analysis of EUR1 and AFR1, but the LD reference was EUR1_EUR2.

In a second simulation scenario we dropped the assumption that all causal SNPs are genotyped. To achieve this, we kept all simulation settings identical to those described in Section 2.1, with the sole exception that we restricted the analysis to HapMap3 SNPs (using gcta --extract and Manc-COJO --extract).

In the third simulation scenario we dropped the assumption of the in-sample LD reference panel. To do this, we downloaded genotype data from the 1000 Genomes Project and randomly sampled two sets of unrelated individuals (N = 503) from European participants, as defined by the 1000 Genomes Project, to serve as the LD reference for EUR. Similarly, we selected N = 503 African individuals as the LD reference for AFR.

In the final simulation scenario we allowed SNP effect sizes to differ across ancestries, although retaining direction of association. As in previous settings, all simulation parameters matched those in Section 2.1, except that we relaxed the requiring identical effect sizes across ancestries. Instead, we sampled the true effect sizes independently in the two ancestries, while ensuring that the directions (sign) of effect were consistent.

##### 2.3 Simulation for Manc-COJO:MDISA

Simulations in Sections 2.1 and 2.2 assumed shared causal SNPs across ancestries. In this study, we developed Manc-COJO:MDISA to account for scenarios where ancestry-specific causal variants exist, and tested it in simulation. In our simulations, 60% of the causal SNPs in each block were shared across ancestries, while 20% were specific to each of the two ancestries. For example, in a region with 10 simulated causal SNPs, 6 were shared, 2 were European-specific, and 2 were African-specific. We simulated scenarios with either 5, 10, or 20 causal SNPs per region. The effect sizes of shared causal SNPs were identical across ancestries. Phenotype simulation followed the same procedure as described in Section 2.1. In total, we conducted simulations in 38 blocks, across 3 different numbers of causal SNPs, with 50 replicates per combination, resulting in 5,700 simulated scenarios.

### Supplementary Methods 3. Real data analysis

We evaluated the performance of Manc-COJO in real data, using ancestry-specific GWAS summary statistics of European and African ancestries for four lipid traits chosen because of their large sample sizes for both European and African ancestries. Global Lipids Genetics Consortium (GLGC) GWAS summary statistics (which excluded UK biobank) were available for high-density lipoprotein cholesterol (HDL), low-density lipoprotein cholesterol (LDL), total cholesterol (TC), and log-transformed triglyceride levels (logTG)[8].

#### 3.1 Data quality control

We used GWAS summary statistics that excluded participants from the UK Biobank. Only common SNPs (MAF $\geq$ 0.01) present in both ancestries and bi-allelic SNPs were retained for analysis. Further filtering was conducted to retain only those SNPs that were genotyped in the UK Biobank across both ancestries and had an MAF $\geq$ 0.01 among unrelated African participants ($N$ = 6,901). Following quality control, the median per-SNP sample sizes and the number of SNPs included for each trait were as follows: HDL ($N_{AFR}$ = 90,804, $N_{EUR}$ = 881,859, $M$ = 6,514,392 SNPs); LDL ($N_{AFR}$ = 87,759, $N_{EUR}$ = 837,509, $M$ = 6,511,658 SNPs); TC ($N_{AFR}$ = 92,554, $N_{EUR}$ = 924,769, $M$ = 6,514,274 SNPs); logTG ($N_{AFR}$ = 89,467, $N_{EUR}$ = 858,015, $M$ = 6,514,129 SNPs).

#### 3.2 Data input

For each trait, the inputs to the Manc-COJO algorithm consisted of two separate GWAS summary statistics (one for each ancestry) and two corresponding LD reference panels: a randomly sampled set of 69,010 unrelated European individuals and 6,901 unrelated African individuals from the UK Biobank. Default parameters were applied: a window size of 10Mb (assuming SNPs more than 10Mb apart to be independent), a collinearity threshold of 0.9, and a significance threshold of $5\times10^{-8}$, consistent with the original Sanc-COJO protocol.

We benchmarked Manc-COJO against Sanc-COJO applied to both multi-ancestry and single-ancestry datasets. For the multi-ancestry analysis, the input was the inverse-variance meta-analysis of the aforementioned European and African GWAS summary statistics from GLGC. The LD reference was either a merged cohort of 69,010 European and 6,901 African ancestry individuals, or a merged cohort of 69,010 European and 6,901 European ancestry individuals (using PLINK's –bmerge function). For the single-ancestry analysis, new GWAS were conducted on 90,000 unrelated European UK Biobank samples for each of the lipid traits. Covariate adjustment followed the original GIANT and GLGC protocols: for lipid traits, covariates were age, age-squared, batch, sex, assessment centre, and the first 20 PCs. For individuals on cholesterol-lowering medications (Field IDs p6153_i0 and p6177_i0), LDL ($LDL_{adj}$ = LDL/0.7) and TC ($TC_{adj}$ = TC/0.8) levels were adjusted. Analyses were sex-stratified: residuals were computed separately for males and females and then pooled. Subsequently, we conducted an inverse-variance meta-analysis of new GWAS summary statistics derived from UKB European participants and aforementioned European-specific GWAS, excluding UK Biobank data, from the GLGC. The LD reference for single-ancestry Sanc-COJO was a merged cohort of 69,010 and 6,901 unrelated European UK Biobank individuals. In general, we ensured that the input GWAS summary statistics involved approximately equal sample sizes and identical SNP sets, and LD reference panel sizes are comparable across methods for fair comparison (**Supplementary Table 1**).

Given that samples from GLGC were heavily skewed towards European ancestry (approximately 10:1 ratio of European to African participants after excluding UK Biobank samples), we conducted a down-sampling analysis to examine whether closer ancestry balance (approximately 1:1) would enhance detection of independent associations. Four approaches were again evaluated: (1) Manc-COJO applied to multi-ancestry data; (2) Sanc-COJO applied to multi-ancestry data using mixed-ancestry LD references (Sanc-COJO→Manc); (3) Sanc-COJO applied to multi-ancestry data using European-only LD references (Sanc-COJO→Manc/EURref); and (4) Sanc-COJO applied to single-ancestry data. For Manc-COJO, we used African-specific GWAS from GLGC and newly generated European-specific GWAS from UK Biobank (N ~ 90,000 per lipid trait). For multi-ancestry Sanc-COJO, input GWAS were derived from inverse-variance meta-analysis of the African- and European-specific GWAS used for Manc-COJO. LD reference panels consisted of either 6,901 unrelated Europeans plus 6,901 unrelated Africans (Sanc-COJO→Manc) or two independent sets of 6,901 unrelated Europeans (Sanc-COJO→Manc/EURref). For single-ancestry Sanc-COJO, GWAS summary statistics were generated from ~180,000 unrelated EUR individuals per lipid trait in UK Biobank using the same analytical pipeline as in the full-scale analyses, and the corresponding LD reference comprised 13,802 unrelated Europeans excluding individuals contributing to the GWAS (**Supplementary Table 1**).

##### 3.3 Out-of-sample prediction in UK Biobank

To obtain the weights for polygenic prediction and ensure a fair comparison across all methods, we first identified the list of SNPs selected by the Manc-COJO and Sanc-COJO models. These SNPs were then jointly **re-estimated** using the GCTA --cojo-joint command, based on the summary statistics and LD reference panels specified in **Supplementary Table 1** (for full-sample analyses) and **Supplementary Table 2** (for down-sampled analyses). After obtaining the joint effect sizes for each model, polygenic scores were computed using the PLINK --score command.

To evaluate out-of-sample prediction performance in European cohorts, we included 47,066 unrelated European participants for lipid traits, none of whom were included in previous GWAS analyses. Prediction into African cohorts was assessed using 3,000 unrelated UKB African participants for lipid traits. Residuals for all traits were computed following the same covariate adjustments described for the GWAS analyses.

To estimate the prediction error, we randomly divided 47,066 unrelated UKB EUR participants into 50 groups, and 3,000 UKB AFR participants into 20 groups, with one group having more individuals if the sample size was not evenly divisible. Within each group, a simple linear regression (residual ~ PGS + error) was performed, and the out-of-sample $R^2$ was calculated as the coefficient of determination. The final $R^2$ estimate was averaged across groups for each trait and ancestry, and the standard error of the estimate was computed.

To compare genomic regions where the Manc-COJO and Sanc-COJO models identified different numbers of SNPs, we partitioned the genome into LD-independent blocks based on a previously published framework[9]. Note that the LD blocks used here differ from those in the simulation study, as we opted for a more conservative definition to reduce the likelihood of false positive claims in real data analysis. For each individual LD block, the out-of-sample prediction accuracy of each COJO model (Manc-COJO or Sanc-COJO) was estimated by first removing the COJO-selected SNPs within that block, recalculating the joint effect sizes using the remaining SNPs, and then computing the difference in prediction accuracy between the full model and the reduced model.

##### 3.4 Out-of-sample prediction in All of US

We reassessed out-of-sample predictive performance using data from the All of Us cohort. Because SNPs in All of Us are aligned to the GRCh38 reference genome, we first generated a conversion table linking SNP rsIDs and genomic coordinates between the UK Biobank (GRCh37) and All of Us (GRCh38) datasets using the UCSC LiftOver tool. A small subset of variants (~0.3 million of the 8.5 million SNPs with MAF ≥ 0.01 in the UK Biobank) could not be successfully mapped from GRCh37 to GRCh38. We then extracted genotype data from the ACAF call set, which includes SNPs with MAF ≥ 0.01 or an allele count ≥ 100 in any sub-population ("gs://fc-aou-datasets-controlled/v8/wgs/short_read/snpindel/acaf_threshold/plink_bed"). Genotypes were obtained for all AFR and EUR individuals, as defined in "gs://fc-aou-datasets-controlled/v8/wgs/short_read/snpindel/aux/ancestry/ancestry_preds.tsv". We excluded related individuals and those who did not pass quality control, as indicated in "gs://fc-aou-datasets-controlled/v8/wgs/short_read/snpindel/aux/relatedness/" and "gs://fc-aou-datasets-controlled/v8/wgs/short_read/snpindel/aux/qc/flagged_samples.tsv".

To ensure a fair comparison, we repeated the Manc-COJO and Sanc-COJO analyses using the approximately ~1-million-sample version of the summary statistics (~0.9 million EUR plus 0.09 million AFR or EUR) as described above, but retained only SNPs that were present in both the UK Biobank and All of Us cohorts at the start of the analysis.

For lipid traits, we extracted phenotypes defined as "Cholesterol [Mass/volume] in Serum or Plasma," "Cholesterol in HDL [Mass/volume] in Serum or Plasma," "Cholesterol in LDL [Mass/volume] in Serum or Plasma," "Cholesterol in LDL [Mass/volume] in Serum or Plasma by calculation," and "Triglyceride [Mass/volume] in Serum or Plasma." We retained only measurements reported in units of "milligram per deciliter" or "milligram per deciliter calculated." For individuals with multiple measurements of the same trait, we kept the earliest measurement. For participants taking lipid-lowering medications (Atorvastatin, Ezetimibe, Fluvastatin, Lovastatin, Pravastatin, Rosuvastatin, Simvastatin), LDL and total cholesterol levels were adjusted as in the GLGC study (LDL_adj = LDL/0.7; TC_adj = TC/0.8), provided the participant was actively taking the medication at the time of measurement, determined by comparing medication exposure start and end dates with the lipid measurement date.

We then adjusted each trait for age, sex, and the first 16 PCs, and calculated scaled residuals separately within each ancestry group. After generating residuals, we divided EUR (~80,000 per trait) and AFR (~25,000 per trait) samples into groups of 1,000 and applied the same protocol used in the UK Biobank to compute PGS.

#### Supplementary Notes 1. Simulation sensitivity analysis

In practice, model assumptions are often violated. In addition to the scenarios described in the main text, we conducted analyses restricted to HapMap3 SNPs to mimic situations in which causal SNPs are not genotyped, simulations in which effect sizes differed between ancestries, and simulations in which LD matrices were not calculated from the GWAS samples themselves. As expected, the performance of all models declined under these non-idealised conditions. Nonetheless, Manc-COJO consistently exhibited greater robustness than the Sanc-COJO-based methods.

Causal SNPs are not always genotyped in practice. To simulate this scenario, we retained the simulation settings used under the idealised condition but restricted the analysis to HapMap3 SNPs only. All three methods performed reasonably well in European ancestry, with the resulting models identifying SNPs at similar LD distances from the causal variants. However, the Manc-COJO method yielded fewer false positive findings. Specifically, the false positive indicator (calculated using LD from European ancestry samples) was lower for Manc-COJO compared to Sanc-COJO->Manc with in-sample, mixed-ancestry LD references: 0.03 vs 0.04, 0.09 vs 0.11, 0.11 vs 0.14, and 0.13 vs 0.16 for regions with 1, 5, 10, and 20 causal SNPs, respectively. In contrast, when calculating false positive and power metrics using LD from African ancestry samples, Sanc-COJO—as expected—exhibited reduced power and increased false positive rates. While the Sanc-COJO->Manc with in-sample, mixed-ancestry LD reference demonstrated comparable power to Manc-COJO, it consistently resulted in higher numbers in false positive indicator: 0.18 vs 0.17, 0.25 vs 0.21, 0.27 vs 0.23, and 0.30 vs 0.25 for 1, 5, 10, and 20 causal SNPs in a region, respectively (**Supplementary Figure 3**).

In our modelling framework, we initially assumed that the true effect sizes of causal SNPs were identical across ancestries. To assess robustness under non-idealised scenarios, we relaxed this assumption in simulation, while keeping all other parameters consistent with the idealised scenarios. In all simulated conditions, the effect sizes of the causal SNPs varied across ancestries; however, the direction of association remained consistent. Under these settings, Manc-COJO consistently outperformed both Sanc-COJO->Manc with in-sample, mixed-ancestry LD reference and Sanc-COJO by identifying a greater number of causal SNPs, demonstrating higher statistical power, and yielding fewer false positive discoveries (**Supplementary Figure 4**).

Finally, we simulated scenarios with imperfect LD references. The simulation settings were identical to those of the idealised scenarios, with the sole exception that, instead of using in-sample LD, the LD references were derived from the 1000 Genomes Project. For this, we randomly selected $N$ = 503 unrelated European individuals and $N$ = 503 unrelated African individuals as a reference sample (see **Supplementary Methods**). Generally, the trends remain consistent with the idealised scenario: Manc-COJO or Sanc-COJO->Manc identify a greater number of causal SNPs and exhibit superior statistical power compared to Sanc-COJO, while among the two multi-ancestry approaches, Manc-COJO generates fewer false-positive discoveries. However, compared to the scenario using in-sample LD, the ability to detect independent associations diminishes across all methods. For example, the median number of causal SNPs identified by Manc-COJO decreases from 6 to 5 and from 11 to 8 when there are 10 and 20 causal SNPs in a region, respectively. Concurrently, the median false-positive indicator rises sharply, increasing from 0 to 0.01, 0.01 to 0.09, and 0.03 to 0.19 when there are 5, 10, and 20 causal SNPs in a region, respectively (**Supplementary Figure 5**). This underscores the benefit of using in-sample LD.

#### Supplementary Notes 2. Down-sample analysis

In the main text, we down-sampled the EUR GWAS to match the sample size of the AFR GWAS to show that using a EUR-only LD reference can inflate false-positive signals when the ancestral proportion in the meta-analysis of GWAS summary statistics approaches 1:1. In addition to this analysis, we repeated the down-sampling procedure shown in **Extended Figure 4** to evaluate whether achieving a more balanced ancestry composition (~1:1) in GWAS sample sizes enhances the detection of independent associations (**Supplementary Table 2**). Given the reduced GWAS sample sizes the $R^2$ values in all analyses were naturally lower. In EUR cohorts, the out-of-sample $R^2$ (and standard error) for Manc-COJO models were lower than that of Sanc-COJO models: 0.126 (0.022) vs 0.134 (0.020) for HDL, 0.139 (0.022) vs 0.144 (0.024) for LDL, 0.076 (0.013) vs 0.083 (0.015) for logTG, and 0.125 (0.022) vs 0.130 (0.020) for TC. Nonetheless, in African cohorts, Manc-COJO models demonstrated consistently better trans-ancestry portability, achieving higher out-of-sample $R^2$ values than the Sanc-COJO models: 0.090 (0.042) vs 0.073 (0.032) for HDL, 0.142 (0.047) vs 0.128 (0.045) for LDL, 0.046 (0.024) vs 0.012 (0.0080) for logTG, and 0.121 (0.044) vs 0.116 (0.047) for TC. Notably, as the ancestry balance approached parity, the difference in model size between the methods became more pronounced: Manc-COJO consistently yielded more compact models with fewer selected SNPs—139 vs 182 for HDL, 118 vs 141 for LDL, 97 vs 144 for logTG, and 139 vs 157 for TC (**Supplementary Figure 6**). Overall, although Sanc-COJO models showed higher predictive accuracy in EUR cohorts under the down-sampled setting—because the EUR GWAS summary statistics used by Manc-COJO were now much less powered than those used by Sanc-COJO—Manc-COJO models still retained reasonable accuracy in European cohorts and outperformed Sanc-COJO in African cohorts while including up to 30% fewer SNPs. These results suggest that a more balanced ancestry composition may improve Manc-COJO's ability to detect independent associations, but we currently lack sufficiently large non-EUR datasets—particularly AFR datasets, which have the shortest LD among all ancestries—to directly demonstrate this. This highlights the importance of increasing sample sizes for non-European ancestries.

### Supplementary Notes 3. Algorithmic improvements over orginal COJO

#### Supplementary Note 3.1 More robust outputs

Although our software is designed for multi-ancestry analyses, it can also be used to perform single-ancestry COJO analyses as implemented in GCTA. Our implementation introduces two refinements to the Sanc-COJO algorithm in GCTA.

First, during early iterations, GCTA stores *p*-values that fall outside the representable range of the C++ double type ($1.7\times10^{-308}$ to $1.7\times10^{308}$) as zero. When several *p*-values are truncated in this way, multiple SNPs become indistinguishably significant, potentially causing a suboptimal SNP to be selected for the joint model. Our implementation instead prioritizes the SNP with the largest absolute z-score, which is mathematically equivalent but avoids numerical underflow.

Across the four lipid traits and 22 chromosomes (88 analyses), nine trait–chromosome combinations were affected by this refinement (**Supplementary Table 3**), with the majority of selected SNPs remaining identical to GCTA outputs. Incorporating this refinement results in slightly fewer selected SNPs and marginally improved prediction accuracy for single-ancestry COJO (**Supplementary Table 4**), as expected given that the algorithm is now guaranteed to retain the truly most significant SNP at each iteration.

Second, in rare scenarios involving complex LD patterns—such as the Sanc-COJO -> Manc analyses in the main text, where European and African genotype matrices are merged for pseudo–single-ancestry analyses—the final iteration may remove SNPs due to collinearity without adding new ones. In GCTA, this can create index inconsistencies and unstable *.cma.cojo* outputs. This issue does not occur for the --cojo-cond option, where the SNP set is fixed by the user, and we did not observe it in standard single-ancestry datasets such as the ~1 million European cohort analysed in this study.

We performed extensive checks and confirmed that our software produces numerically identical results to GCTA (for *.cma.cojo*, *.jma.cojo*, and *.ldr.cojo* outputs) for all analyses in this study, except for scenarios listed in **Supplementary Table 3,** where the implemented refinements improved upon the original GCTA software.

#### Supplementary Note 3.2 Improved computation efficiency

The main speedup in our implementation comes from recoding genotype values into compact binary form and operating on them with native bitwise instructions. Instead of storing SNP genotypes as arrays of 0, 1, and 2 and performing arithmetic on *int* or *double* types, we pack them into machine-width integers such as *uint64_t*. This enables each CPU instruction to process many SNPs simultaneously using bitwise AND, OR, NOT, and population-count operations. Because these instructions execute as low-level hardware operations, they avoid the overhead of scalar arithmetic and floating-point pipelines. As a result, genotype storage and computation become more memory-efficient, cache-aligned, and effectively executed at hardware speed, yielding substantial throughput gains for large-scale COJO analyses. Other improvements include reusing intermediate results across correlation matrices, integrating optimized C++ libraries to accelerate input data parsing, and other engineering refinements. Full implementation details are available on our open-source GitHub repository to ensure reproducibility.

#### Supplementary Note 3.3 Dealing with data missingness

In real data analyses, missingness can arise in both the genotypic data used to compute LD references—due to factors such as genotyping errors or imputation failure—and in the GWAS summary statistics themselves, where genotype missingness within a cohort or differences in the SNP sets reported across cohorts in a meta-analysis can lead to substantial variation in sample size across SNPs. Out-of-sample prediction introduces an additional layer of complexity, as genotype missingness may also occur in the target cohort used for prediction.

In LD calculations, different software handle missing genotypes differently. Tools such as GCTA impute missing genotypes by replacing them with the mean genotype value, whereas software like PLINK exclude all pairs containing missing values when computing pairwise correlations. Both approaches have limitations. Mean imputation reduces the genotypic variance of the SNP, causing the variance to deviate from HWE. In contrast, removing pairs with missing data leads to less precise estimates of genetic correlation, particularly when the LD-reference sample size is not large enough.

In addressing missingness in GWAS summary statistics, GCTA introduces an additional adjustment when computing joint effects (as in equation 4) and conditional effects (as in equation 7). Although this adjustment is not explicitly presented in the original GCTA-COJO paper, what GCTA effectively does is rescaling the pairwise LD correlations according to the per-SNP effective sample sizes in GWAS summary statistics. Specifically, the correlation between SNP *i* and SNP *j* is multiplied by $\min(\hat{n}_i,\hat{n}_j)/\hat{n}_j$, and the correlation between SNP j and SNP i by $\min(\hat{n}_i,\hat{n}_j)/\hat{n}_i$ where $\hat{n}_i$ is the per-SNP effective sample sizes of SNP *i* defined in equation (11).

When calculating polygenic scores, mean imputation is commonly used to handle missing genotype data.

In most COJO- or fine-mapping–based workflows, the calculation of joint effects for polygenic prediction is performed immediately after variable selection and typically uses the same strategy for handling missingness. Interestingly, our results show that variable selection and the estimation of joint effects are in fact two distinct steps that should be treated differently. In our study, we evaluated three approaches for variable selection—(i) excluding individuals with missing genotypes on a per–SNP-pair basis during pairwise LD computation, (ii) imputing missing values when calculating pairwise LD, and (iii) imputing missing values and adjusting for missingness in GWAS summary statistics using the GCTA-based method described above. We applied the same three strategies when computing joint effects for polygenic prediction, yielding nine possible combinations of variable-selection and joint-effect procedures.

For variable selection, the optimal strategy varied across traits. However, in out-of-sample prediction, the approach that imputes missing values and adjusts GWAS summary statistics for missingness consistently outperformed the other two strategies—an unsurprising result given how missingness is handled in the target sample (**Supplementary Table 5**). For simplicity, in the main text we report results using the “remove missing values” strategy for variable selection and the “impute missing values plus adjust for missingness” strategy for estimating joint effects.

Our software implements all three missingness-handling strategies, allowing users to choose the method most appropriate for their data or explore combinations beyond those we evaluated. For example, when the LD-reference sample is sufficiently large, a user may choose the “remove missing values” strategy and then correct marginal GWAS effects by scaling them with $\sqrt{INFO}$[10], where INFO can be approximated by the ratio of per-SNP sample size to the total sample size.

##### Supplementary Note 3.4 Other features

Beyond refinements described above, our software also introduces several new features, including support for PLINK-formatted LD matrices and the ability to fix known associated SNPs (e.g., validated through functional experiments) during iterative selection. In addition, it enables multi-trait conditional analyses. For example, to identify SNPs independently associated with bipolar disorder (BD) conditional on SNPs independently associated with major depressive disorder (MDD), a user may first run Manc-COJO on MDD and then fix the COJO SNPs identified for MDD when running Manc-COJO for BD.

#### Supplementary Notes 4. Choice of $R^2$ parameter

In the Manc-COJO algorithm, it is possible to use changes in $\hat{R}^2_{adj}$ as an *optional* selection criterion for researchers who prefer a more conservative model in regions with extremely complex LD structure. Users can specify thresholds that require the $\hat{R}^2_{adj}$ of the model to increase by a certain percentage during both forward and backward selection in *both* ancestries for a SNP to be included in the model as described in the **Supplementary Method** section.

To evaluate the impact of applying different $\hat{R}^2_{adj}$thresholds, we tested two scenarios in both simulation and real-data analyses: (1) $\hat{R}^2_{adj}$ in both forward and backward selection was set to 0. In this setting, for a SNP to be included in the model, the $\hat{R}^2_{adj}$ of the new model after forward and backward selections must be higher than that of the previous model in both ancestries. (2) $\hat{R}^2_{adj}$ in forward selection was set to 0, and in backward selection set to −1. In this scenario, we imposed a threshold only for forward selection, requiring an increase in $\hat{R}^2_{adj}$ to add a SNP, while allowing any decrease in $\hat{R}^2_{adj}$ during backward selection. We performed the simulation with a EUR:AFR ratio of 10:1 to match the ancestry proportions used in the real-data analysis.

In general, applying a more stringent $\hat{R}^2_{adj}$ threshold reduces the false positive inclusion of non-causal SNPs in the model, but at the cost of reduced power to include causal SNPs, especially when the number of causal SNPs increases (**Supplementary Figure 7**).

In real data analysis, applying an $\hat{R}^2_{adj}$ threshold of 0 in both forward and backward selection substantially reduced model size compared with the default setting that imposes no $\hat{R}^2_{adj}$ restriction. Model sizes decreased from 831 to 759 for HDL, 554 to 511 for LDL, 655 to 587 for logTG, and 700 to 654 for TC. Introducing an $\hat{R}^2_{adj}$ threshold generally led to a slight reduction in out-of-sample prediction accuracy in EUR; however, for HDL and LDL we observed a small increase in AFR despite the smaller model sizes, and the available sample size did not allow us to determine the significance of this pattern (**Supplementary Table 6**).

Overall, although the default setting without an $\hat{R}^2_{adj}$ threshold performs well in most scenarios, adding such a threshold may help reduce false-positive signals in regions with highly complex LD structures, as illustrated in HDL and LDL. The optimal choice may therefore depend on the genetic architecture of the trait under study.

#### Supplementary Notes 5. Insight into Manc-COJO vs Sanc-COJO output

In order to understand the factors contributing to differences in model size between the Manc-COJO and Sanc-COJO models, we partitioned SNPs identified in the full-sample analysis (using lipid GWAS traits, 6.5M SNPs, 0.9 million EUR + 0.09 million AFR for Manc-COJO and 0.9 million EUR + 0.09 million EUR for Sanc-COJO) into LD-independent blocks as defined previously[9] (**Supplementary Methods**). We then compared the number of SNPs included in the Manc-COJO model with those in the single-ancestry Sanc-COJO model within each LD-independent block (**Supplementary Figure 8**).

For most LD blocks (specifically, 64.2% of blocks for HDL, 66.2% for LDL, 61.7% for TC, and 68.7% for logTG) the Manc-COJO and Sanc-COJO models identified an identical number of independent associations. As the example shown in the main text, within these blocks, although the number of indepdent associations is the same, Manc-COJO identifies independently associated SNPs that are more closely correlated (in terms of LD) with the causal variants than those flagged by Sanc-COJO. We observed similar patterns in out-of-sample prediction accuracy. For LD-independent blocks containing rs2277862 in TC, Manc-COJO SNPs explained $2.7\times10^{-4}$ more phenotypic variance in AFR than Sanc-COJO SNPs, but $3.6\times10^{-6}$ less variance in EUR. For LD-independent blocks containing rs10889356 in TC, the corresponding differences were $3.1\times10^{-4}$ more in AFR and $3.6\times10^{-5}$ less in EUR. For blocks containing rs10872142 in LDL, Manc-COJO SNPs explained $3.7\times10^{-4}$ and $1.4\times10^{-5}$ more phenotypic variance in AFR and EUR, respectively. Because LD correlations between Manc- or Sanc-COJO SNPs and the causal SNPs are uniformly high and similar in EUR, it is not unexpected that Sanc-COJO SNPs can, due to random error, show slightly higher out-of-sample $R^2$ in this population. In AFR, however, Manc-COJO SNPs consistently yield higher out-of-sample $R^2$, and the magnitude of the difference is larger (on the order of $10^{-4}$). Although the out-of-sample prediction sample sizes are too small to support statistically significant differences, the observed patterns are notably consistent with expectations.

The main reason the Manc-COJO model is generally smaller than the Sanc-COJO model is that in many LD blocks, the Sanc-COJO model selects one or occasionally two SNPs, whereas the Manc-COJO model does not select any SNPs in those blocks. Specifically, this pattern accounts for 69 SNPs across 62 blocks for HDL, 50 SNPs across 46 blocks for LDL, 48 SNPs across 43 blocks for TC, and 70 SNPs across 64 blocks for logTG (**Supplementary Figure 8**). These SNPs identified by the Sanc-COJO model make up approximately 6–10% of all SNPs in the model and are not rare variants, with mean minor allele frequencies of 0.27 for HDL, 0.28 for LDL, 0.26 for TC, and 0.28 for logTG in the European population. However, they collectively explain only a small proportion of phenotypic variance. The sum of out-of-sample prediction accuracies for these SNPs when predicting into European cohorts was 0.0012 for HDL, 0.0018 for LDL, 0.0014 for TC, and 0.0025 for logTG. Prediction accuracies in African cohorts were even lower, at 0.00043, −0.0010, 0.0011, and −0.0021 for HDL, LDL, TC, and logTG, respectively. The fact that this large set of SNPs consists of common variants yet explains only a negligible proportion of phenotypic variance suggests that at least some of the independent associations tagged by the Sanc-COJO model in these blocks are more likely to represent false-positive signals rather than causal SNPs with small effects.

#### Supplementary Notes 6. Meta-analysis, mega-analysis, and admixed population

There has been extensive discussion on how best to integrate data from multiple ancestries. One approach is to perform GWAS separately within each ancestry and then combine the results using meta-analysis. Alternatively, one can conduct a mega-analysis, in which individual-level genotype data are pooled across ancestries (though this is often challenging due to data-sharing restrictions) and GWAS is performed on the combined cohort. Under idealised conditions—where in-sample LD is available and sample sizes are sufficiently large—mega-analysis can offer greater statistical power, as shown in previous research[11]. However, when in-sample LD is unavailable or conditions deviate from these ideal assumptions, simulations as shown in our study indicate that mega-analysis can be highly susceptible to inflated false-positive rates.

A similar issue arises for admixed populations, where prior work has shown that standard GWAS of admixed cohorts can surpass the power of equally sized multi-ancestry mega-analyses with homogeneous within-ancestry structure[12]. However, COJO analysis in admixed populations also relies heavily on having a perfectly matched LD reference. Considering the constraints on data sharing and the practical difficulties of pooling individual-level data, we therefore chose to use meta-analysis strategies in our study.

Nonetheless, our software can be readily adapted to perform COJO analysis either in a mega-analysis setting or for admixed populations. The main caveat is that, as shown in Equation (8), when underlying subpopulations differ in allele frequencies, the HWE assumption may be violated, and the genotype variance can no longer be approximated by 2pq. In these situations, researchers should replace the allele-frequency column in the input file with the effective allele frequency, which can be obtained by solving the quadratic equation that equates $2p(1 - p)$ with the observed genotype variance in the sample. Importantly, in-sample LD should always be used in these cases.

#### Supplementary Tables and Figures

| Variable Selection | | | | |
|---|---|---|---|---|
| **Scenarios** | **Methods** | **Input GWAS for EUR** | **Input GWAS for AFR** | **LD reference** |
| **Multi-ancestry Analysis** | Manc-COJO | N ~ 0.9 million EUR GWAS from GLGC | N ~ 0.09 million AFR GWAS from GLGC | 69,010 UKB EUR participants and 6,901 UKB AFR participants (separately) |
| | Sanc-COJO->Manc: Sanc-COJO on multi-ancestry data with multi-ancestry LD reference | | | 69,010 UKB EUR participants and 6,901 UKB AFR participants (Merged into a single cohort) |
| | Sanc-COJO->Manc/EURref: Sanc-COJO on multi-ancestry data with European-only LD reference | | | 75,911 UKB EUR participants (As a single cohort) |
| **Single-ancestry Analysis** | Sanc-COJO on single ancestry data with European-only LD reference | N ~ 0.9 million EUR GWAS from GLGC plus N ~ 0.09 million EUR GWAS from UKB | N/A | 75,911 UKB EUR participants (As a single cohort) |
| **Generating polygenic prediction weights by jointly fitting selected variables with --cojo-joint** | | | | |
| **EUR weight** | Output from all COJO models | N ~ 0.9 million EUR GWAS from GLGC plus N ~ 0.09 million EUR GWAS from UKB | N/A | 75,911 UKB EUR participants |
| **AFR weight** | Output from all COJO models | N/A | N ~ 0.09 million AFR GWAS from GLGC | 6,901 UKB AFR participants |

**Supplementary Table 1. Summary of full sample analyses.** The top table summarizes the GWAS summary-statistic inputs and LD reference panels used by each method for variable selection in the full-sample analyses (total sample size ~1 million). The bottom table shows that, after identifying an independent set of associated SNPs, polygenic prediction weights are derived by jointly fitting the selected variants using the –cojo-joint function on the same dataset across methods to ensure a fair comparison.

| Model-fitting | | | | |
| --- | --- | --- | --- | --- |
| **Scenarios** | **Methods** | **Input GWAS for EUR** | **Input GWAS for AFR** | **LD reference** |
| **Multi-ancestry Analysis** | Manc-COJO | N ~ 90,000 EUR GWAS from UKB | N ~ 90,000 AFR GWAS from GLGC | 6,901 UKB EUR participants and 6,901 UKB AFR participants (separately) |
| | Sanc-COJO->Manc: Sanc-COJO on multi-ancestry data with multi-ancestry LD reference | | | 6,901 UKB EUR participants and 6,901 UKB AFR participants (Merged into a single cohort) |
| | Sanc-COJO->Manc/EURref: Sanc-COJO on multi-ancestry data with European-only LD reference | | | 13,802 UKB EUR participants (As a single cohort) |
| **Single-ancestry Analysis** | Sanc-COJO on single ancestry data with European-only LD reference | N ~ 180,000 EUR GWAS from UKB | N/A | 13,802 UKB EUR participants (As a single cohort) |
| **Generating weight for polygenic prediction** | | | | |
| **EUR weight** | Output from all COJO models | N ~ 180,000 EUR GWAS from UKB | N/A | 13,802 UKB EUR participants (As a single cohort) |
| **AFR weight** | Output from all COJO models | N/A | N ~ 90,000 AFR GWAS from GLGC | 6,901 UKB AFR participants |

**Supplementary Table 2. Summary of down-sample analyses.** This table summarizes the GWAS inputs and LD reference panels used for each method, as in Supplementary Table 1, but for the down-sampled setting in which the ancestry proportions are approximately 1:1 (total sample size ~0.2 million).

| Trait, Chr | SNP number (Ours) | SNP number (original GCTA) | Common SNPs |
|---|---|---|---|
| HDL, Chr 15 | 37 | 38 | 31 |
| HDL, Chr 16 | 50 | 58 | 43 |
| LDL, Chr2 | 54 | 54 | 53 |
| LDL, Chr19 | 75 | 77 | 67 |
| logTG, Chr2 | 55 | 55 | 54 |
| logTG, Chr8 | 48 | 46 | 42 |
| TC, Chr1 | 64 | 64 | 63 |
| TC, Chr2 | 67 | 68 | 65 |
| TC, Chr19 | 74 | 76 | 69 |

**Supplementary Table 3. Comparison of the number of SNPs identified by different models in single-ancestry analyses.** This table summarizes the number of SNPs identified using either Sanc-COJO as implemented in GCTA or our refined version of Sanc-COJO. We analysed four lipid traits using 1 million EUR samples and a 75,911-sample EUR LD reference panel. For most trait–chromosome combinations, the number of independently associated SNPs identified is identical between the two approaches, except for the specific scenarios listed in the table.

| Trait | Sanc-COJO implemented in GCTA | Sanc-COJO implemented in our software |
|---|---|---|
| HDL | 903, 0.077±0.044, 0.170±0.021 | 894, 0.077±0.046, 0.170±0.022 |
| LDL | 581, 0.124±0.039, 0.159±0.022 | 579, 0.129±0.044, 0.159±0.023 |
| logTG | 705, 0.027±0.013, 0.108±0.017 | 707, 0.028±0.014, 0.108±0.017 |
| TC | 718, 0.107±0.040, 0.145±0.020 | 715, 0.110±0.044, 0.146±0.020 |

**Supplementary Table 4. Comparison between Sanc-COJO implemented in GCTA and our refined implementation.** This table compares the original Sanc-COJO implementation in GCTA with our updated software. The inputs are the same as those used in Supplementary Table 3. For each cell, the values shown from left to right are: (i) the total number of SNPs in the model, (ii) prediction accuracy in AFR (with s.e.), and (iii) prediction accuracy in EUR (with s.e.). Despite identifying a smaller number of SNPs, our refined Sanc-COJO algorithm achieves equal or higher prediction accuracy in both ancestries.

| Method (Selection_Prediction) | HDL | LDL | logTG | TC |
|---|---|---|---|---|
| GCTA_GCTA | 826<br>0.1675±0.0219<br>0.0836±0.0404 | 545<br>0.1575±0.0207<br>0.1432±0.0478 | 659<br>0.1071±0.0184<br>0.0490±0.0246 | 705<br>0.1406±0.0196<br>0.1139±0.0400 |
| imputeNA_GCTA | 847<br>0.1675±0.0218<br>0.0831±0.0402 | 567<br>0.1587±0.0205<br>0.1447±0.0496 | 684<br>0.1096±0.0183<br>0.0565±0.0298 | 720<br>0.1450±0.0195<br>0.1150±0.0430 |
| removeNA_GCTA | 831<br>0.1684±0.0228<br>0.0838±0.0403 | 554<br>0.1585±0.0212<br>0.1373±0.0456 | 655<br>0.1089±0.0181<br>0.0534±0.0283 | 700<br>0.1447±0.0204<br>0.1158±0.0437 |
| GCTA_imputeNA | 826<br>0.1658±0.0220<br>0.0841±0.0407 | 545<br>0.1541±0.0205<br>0.1412±0.0468 | 659<br>0.1000±0.0181<br>0.0482±0.0258 | 705<br>0.1320±0.0186<br>0.1104±0.0402 |
| imputeNA_imputeNA | 847<br>0.1638±0.0222<br>0.0835±0.0418 | 567<br>0.1532±0.0199<br>0.1439±0.0509 | 684<br>0.0982±0.0172<br>0.0559±0.0298 | 720<br>0.1353±0.0186<br>0.1097±0.0435 |
| removeNA_imputeNA | 831<br>0.1669±0.0233<br>0.0839±0.0432 | 554<br>0.1557±0.0211<br>0.1358±0.0453 | 655<br>0.1043±0.0179<br>0.0523±0.0283 | 700<br>0.1358±0.0194<br>0.1158±0.0453 |
| GCTA_removeNA | 826<br>0.1628±0.0219<br>0.0841±0.0403 | 545<br>0.1518±0.0204<br>0.1406±0.0452 | 658<br>0.0796±0.0172<br>0.0455±0.0256 | 704<br>0.1286±0.0185<br>0.1045±0.0378 |
| imputeNA_removeNA | 847<br>0.1439±0.0220<br>0.0829±0.0420 | 567<br>0.1441±0.0195<br>0.1419±0.0493 | 684<br>0.0742±0.0147<br>0.0548±0.0306 | 720<br>0.1263±0.0186<br>0.0739±0.0369 |
| removeNA_removeNA | 831<br>0.1616±0.0237<br>0.0833±0.0434 | 554<br>0.1502±0.0210<br>0.1352±0.0434 | 655<br>0.0880±0.0155<br>0.0519±0.0285 | 700<br>0.1279±0.0192<br>0.1137±0.0447 |

**Supplementary Table 5. Comparison of variable-selection and joint-effect estimation strategies in Manc-COJO.** In this study, we evaluated three approaches for variable selection that differ in how they calculate pairwise LD in the presence of missing genotype data: (i) removeNA, which excludes individuals with missing genotypes on a per–SNP-pair basis; (ii) imputeNA, which imputes missing genotypes when computing pairwise LD; and (iii) GCTA, which imputes missing values and additionally adjusts for missingness in GWAS summary statistics as described in Supplementary Note 3. We applied the same three strategies when estimating joint effects for polygenic prediction, resulting in nine combinations of variable-selection and joint-effect estimation methods. In each method label, the part before the underscore indicates the strategy used for variable selection, and the part after the underscore specifies the method used for joint-effect estimation in out-of-sample prediction. Each cell reports three quantities: the model size (top row), the prediction accuracy (± s.e.) in EUR (middle row), and the corresponding accuracy in AFR (bottom row). Data inputs are identical to those used in Figure 3.

| Trait | Manc-COJO ($R^2_{Adj}$ threshold - Forward selection: -1; Backward selection: -1 ) | Manc-COJO ($R^2_{Adj}$ threshold - Forward selection: 0; Backward selection: -1) | Manc-COJO ($R^2_{Adj}$ threshold - Forward selection: 0; Backward selection: 0) |
|---|---|---|---|
| HDL | 831 | 759 | 759 |
| LDL | 554 | 511 | 511 |
| logTG | 655 | 587 | 587 |
| TC | 700 | 654 | 654 |

**Size of models (i.e., number of COJO-SNPs):**
**Prediction $R^2$ (and s.e.) in EUR:**

| Trait | Manc-COJO ($R^2_{Adj}$ threshold - Forward selection: -1; Backward selection: -1 ) | Manc-COJO ($R^2_{Adj}$ threshold - Forward selection: 0; Backward selection: -1) | Manc-COJO ($R^2_{Adj}$ threshold - Forward selection: 0; Backward selection: 0) |
|---|---|---|---|
| HDL | 0.168 (0.023) | 0.165 (0.023) | 0.165 (0.023) |
| LDL | 0.158 (0.021) | 0.156 (0.021) | 0.156 (0.021) |
| logTG | 0.109 (0.018) | 0.107 (0.017) | 0.107 (0.017) |
| TC | 0.145 (0.020) | 0.142 (0.021) | 0.142 (0.021) |

**Prediction $R^2$ (and s.e.) in AFR:**

| Trait | Manc-COJO ($R^2_{Adj}$ threshold - Forward selection: -1; Backward selection: -1 ) | Manc-COJO ($R^2_{Adj}$ threshold - Forward selection: 0; Backward selection: -1) | Manc-COJO ($R^2_{Adj}$ threshold - Forward selection: 0; Backward selection: 0) |
|---|---|---|---|
| HDL | 0.0840 (0.040) | 0.0860 (0.046) | 0.0860 (0.046) |
| LDL | 0.137 (0.046) | 0.140 (0.048) | 0.140 (0.048) |
| logTG | 0.0530 (0.028) | 0.0510 (0.031) | 0.0510 (0.031) |
| TC | 0.116 (0.044) | 0.114 (0.046) | 0.114 (0.046) |

**Supplementary Table 6. Comparison of Manc-COJO performance using different adjusted $R^2$ thresholds in forward and backward selections.** This table summarizes the model sizes and prediction accuracies in European and African cohorts obtained under different adjusted $R^2$ threshold settings in the Manc-COJO models. The data inputs are identical to those used in Figure 3. For the lipid traits, using an adjusted $R^2$threshold of either –1 or 0 during backward selection does not affect the results when the forward-selection threshold is set to 0.

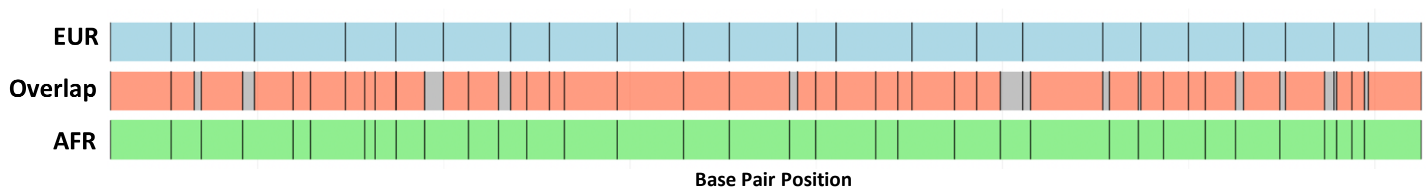

**Supplementary Figure 1. Partitioning of chromosome 22 for simulation.** In our study, Chromosome 22 was partitioned separately for the African and European cohorts, resulting in 33 and 23 blocks, respectively. These blocks were partially overlapping, and a total of 52 overlapping sections were extracted (middle row of the figure). Within each of these 52 overlapping regions, blocks containing fewer than 1,000 SNPs after quality control were excluded (shown in grey in the middle row), leaving 38 blocks (shown in red) for use in the simulation.

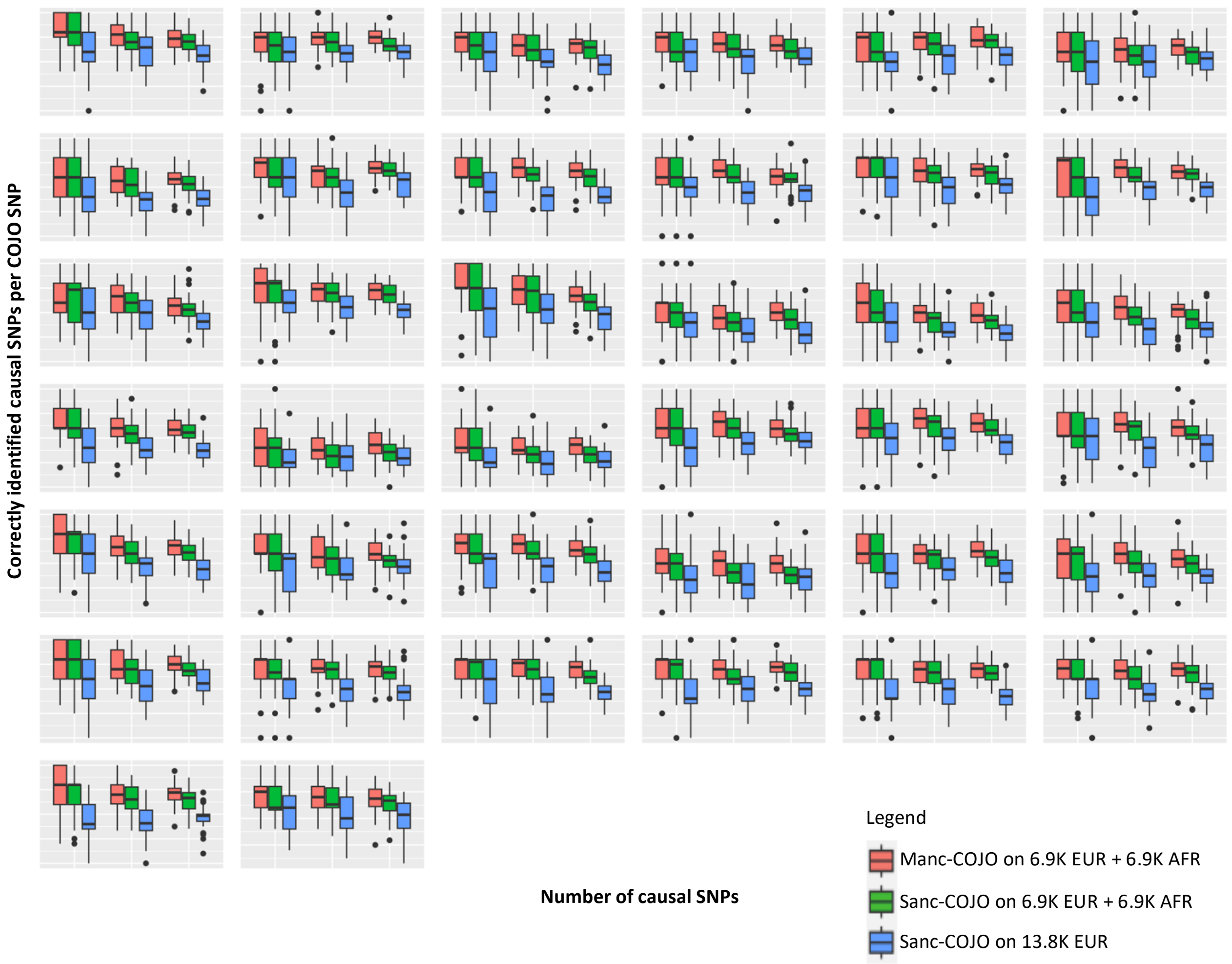

**Supplementary Figure 2. Simulation results within individual blocks are consistent with the pooled results.** This figure summarises the simulation results for all 38 blocks. The plot in the top-left represents Block 1, with Block 2 directly to its right, and so on. Within each plot, the x-axis denotes the number of simulated causal SNPs in that LD-independent block (5, 10, and 20 for the first, second, and third columns, respectively). The y-axis shows the proportion of COJO SNPs that are true causal SNPs. Overall, the results show that the trends observed within individual blocks align closely with those seen in the pooled analysis.

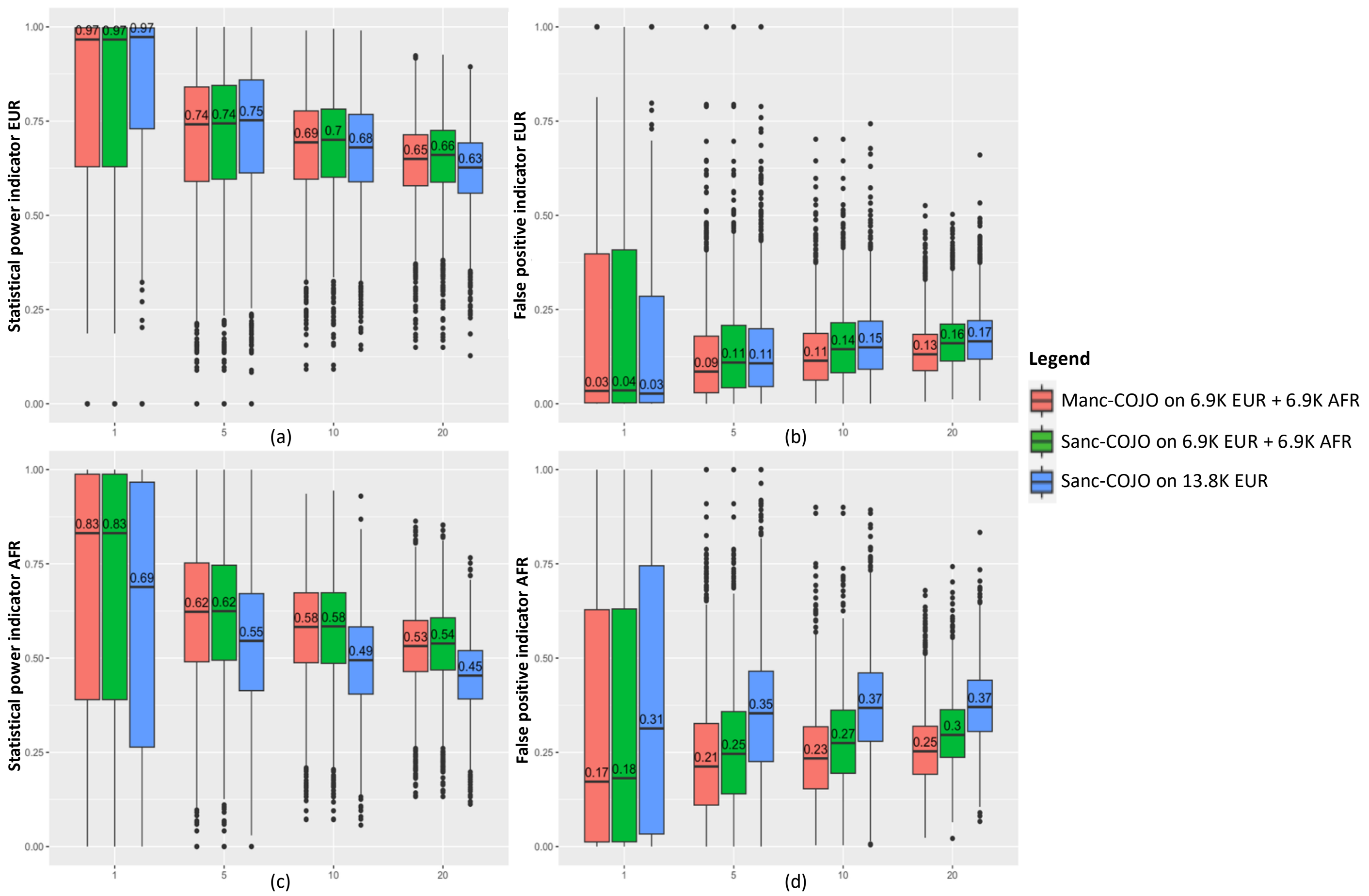

**Supplementary Figure 3. Manc-COJO demonstrates superior performance in simulations when causal SNPs are not genotyped**. This figure summarise simulation results of 7,600 scenarios, and illustrates the performance of Manc-COJO in identifying causal SNPs, showing, on average, superior power and fewer false-positive discoveries across ancestries. We applied our algorithm to both a simulated EUR and a simulated AFR cohort, each with a sample size of 6,900. Our comparisons include Sanc-COJO applied to either 13,800 Europeans or a meta-analysis of 6,900 Europeans with 6,900 Africans. In all scenarios, in-sample LD was utilised. Only Hapmap3 SNPs (~1M) are included in the analysis. The X-axis represents the number of simulated causal SNPs. (a) We identified the COJO SNP with the highest LD correlation (in European cohort) to each causal SNP and averaged these correlations across all causal SNPs. Higher value indicates better power in identifying causal variants. (b) For each COJO SNP, we identified the nearest true causal SNP, calculated their (European) LD, and then averaged these LD values across all COJO SNPs. We then subtracted this mean from 1 to obtain an indicator of false-positive discoveries (i.e., when selected SNPs are not in LD with causal SNPs, the average LD will be low, yielding a higher false-positive indicator). (c)(d) Power and false positive indicators as in (a) and (b), except that African LD is used in calculation. Numbers in each plots are the median value of each scenario.

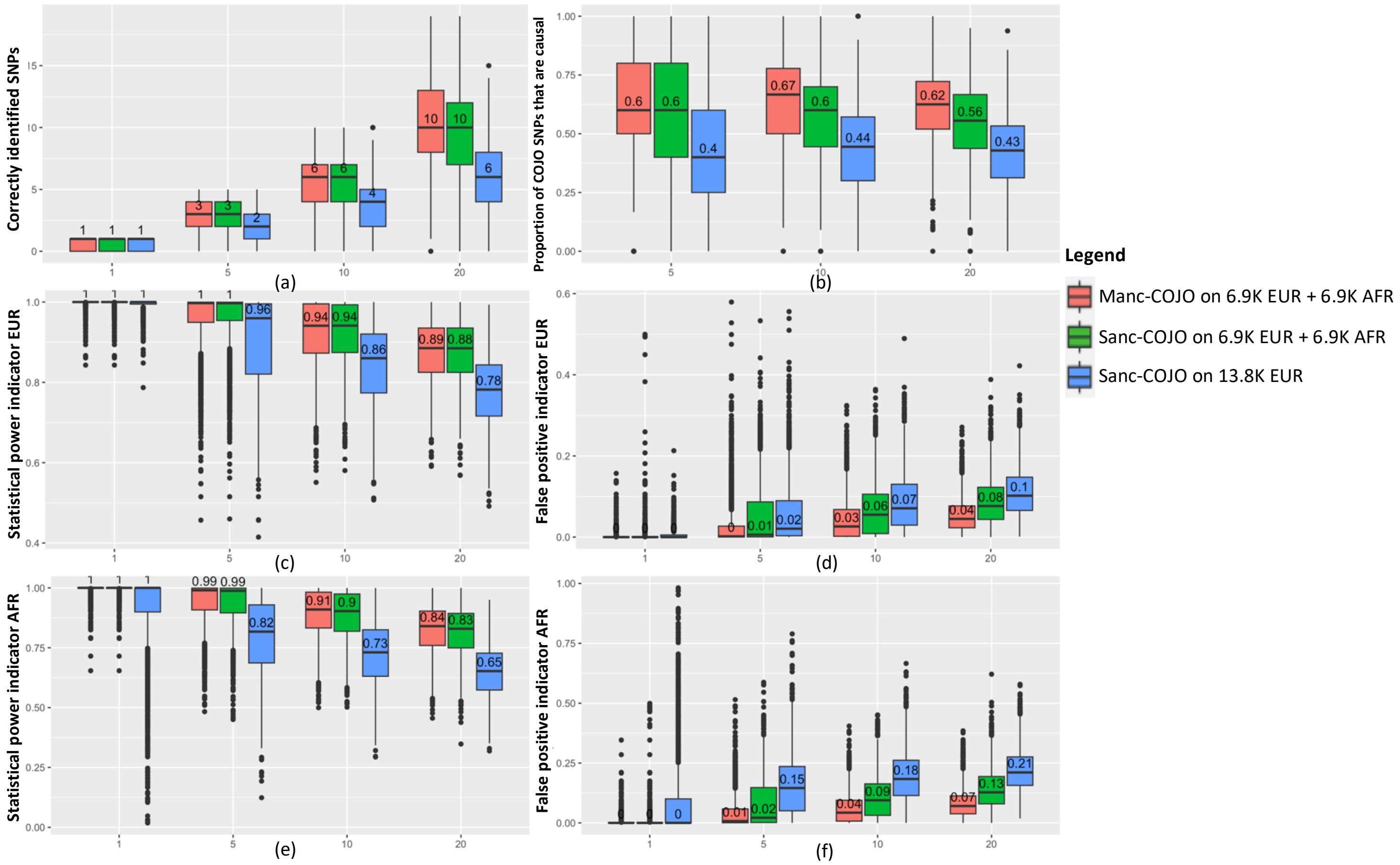

**Supplementary Figure 4**. **Manc-COJO demonstrates superior performance in simulations when effect sizes of causal SNPs are not identical across ancestries.** This figure summarises simulation results of 7,600 scenarios, and illustrates the performance of Manc-COJO in identifying causal SNPs, showing, on average, superior power and fewer false-positive discoveries across ancestries. We applied our algorithm to both a EUR and a simulated AFR cohort, each with a sample size of 6,900. Our comparisons include Sanc-COJO applied to either 13,800 Europeans or a meta-analysis of 6,900 Europeans with 6,900 Africans. In all cases, in-sample LD was utilised. The effect sizes of the causal SNPs varied across the simulated conditions, but the direction of association remained consistent. The X-axis represents the number of simulated causal SNPs. (a) The Y-axis indicates the number of causal SNPs correctly identified by each algorithm. (b) The Y-axis represents the proportion of identified SNPs (termed COJO SNPs) that are causal. (c) As COJO algorithms may identify SNPs near the causal SNP rather than the causal SNP itself, we identified the COJO SNP with the highest LD correlation (in the European cohort) to each causal SNP and averaged these correlations across all causal SNPs; higher values reflect greater power in detecting causal variants. (d) For each COJO SNP, we identified the nearest true causal SNP, calculated their (European) LD, and then averaged these LD values across all COJO SNPs. We then subtracted this mean from 1 to obtain an indicator of false-positive discoveries (i.e., when selected SNPs are not in LD with causal SNPs, the average LD will be low, yielding a higher false-positive indicator). (e)(f) Power and false-positive indicators as in (c) and (d), respectively, but calculated using African LD. Numbers in each plot represent the median value for each group. This figure illustrates that Manc-COJO, on average, identifies a greater number of causal SNPs, exhibits superior power, and produces fewer false-positive discoveries when sample sizes across ancestries are unequal.

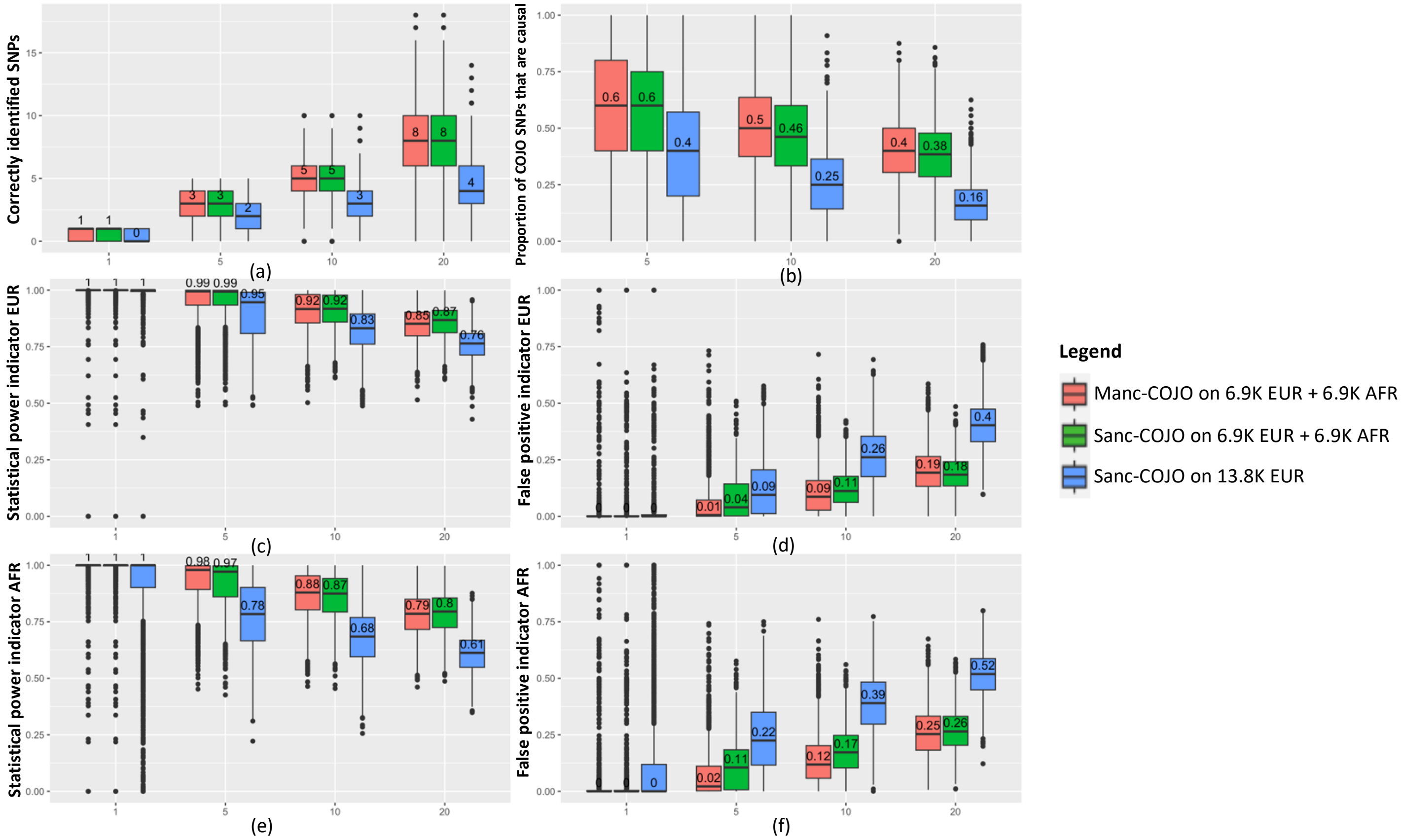

**Supplementary Figure 5. Manc-COJO is robust to imperfect LD reference in simulations**. This figure summarise simulation results of 7,600 scenarios, and illustrates Manc-COJO is robust to imperfect LD reference. We applied our algorithm to both a simulated EUR and a simulated AFR cohort, each with a sample size of 6,900. Our comparisons include Sanc-COJO applied to either 13,800 Europeans or a meta-analysis of 6,900 Europeans with 6,900 Africans. An LD reference from the 1000 Genomes Project was applied in both European (N = 503) and African ancestry (N = 503). The x-axis represents the number of simulated causal SNPs. (a) The y-axis denotes the number of causal SNPs correctly identified by each algorithm. (b) The y-axis denotes the proportion of COJO SNPs that are causal SNPs. (c) Instead of identifying the causal SNP, COJO algorithms may identify SNPs in nearby locations. We identified the COJO SNP with the highest LD correlation (in European cohort) to each causal SNP and averaged these correlations across all causal SNPs. Higher value indicates better power in identifying causal variants. (d) For each COJO SNP, we identified the nearest true causal SNP, calculated their (European) LD, and then averaged these LD values across all COJO SNPs. We then subtracted this mean from 1 to obtain an indicator of false-positive discoveries (i.e., when selected SNPs are not in LD with causal SNPs, the average LD will be low, yielding a higher false-positive indicator). (e)(f) Power and false positive indicators as in (c) and (d), except that African LD is used in calculation. Numbers in each plot are the median value of each group.

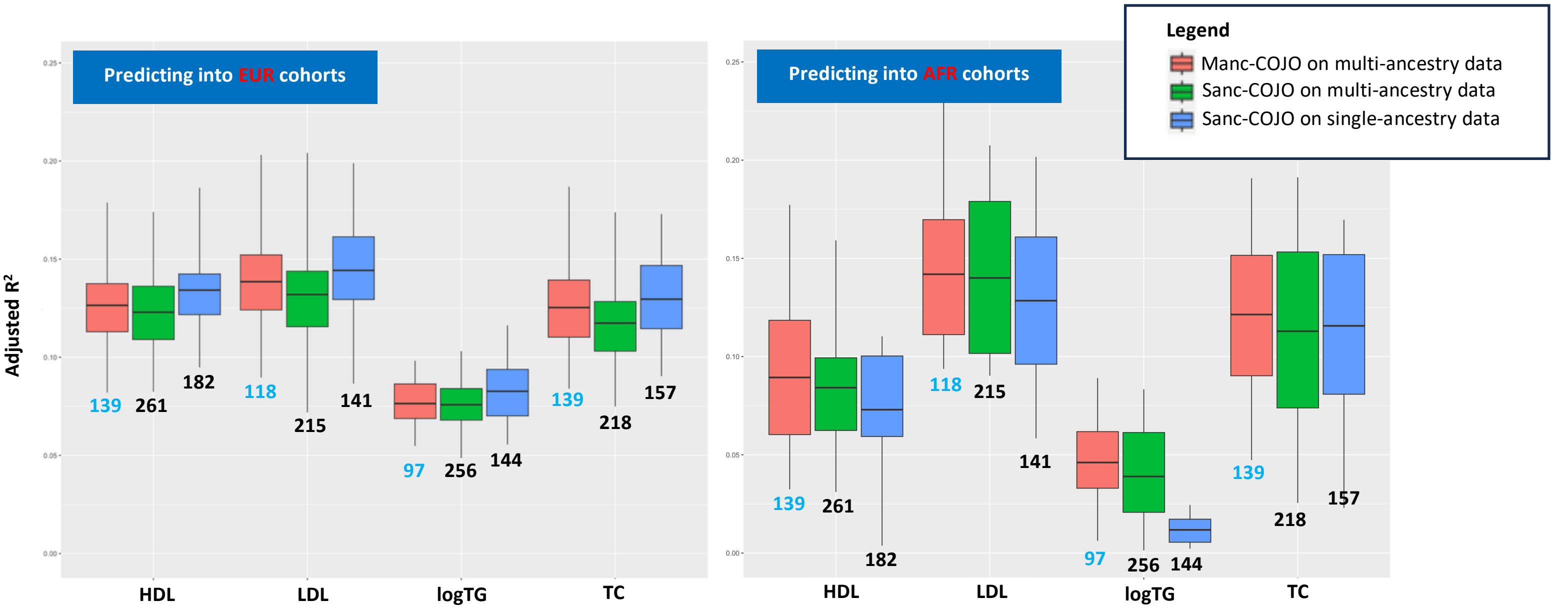

**Supplementary Figure 6. Manc-COJO demonstrates superior trans-ancestry portability compared to the single-ancestry Sanc-COJO model in down-sample analysis applied in real data.** We conducted a down-sampling analysis to examine whether closer ancestry balance (approximately 1:1) would enhance detection of independent associations. This figure summarises the out-of-sample prediction accuracies of Manc-COJO on multi-ancestry data, Sanc-COJO on multi-ancestry data, and Sanc-COJO on single-ancestral data for HDL, LDL, logTG, and TC when predicting into independent European and African cohorts. Numbers in the figure are sizes of the model (i.e., numbe of COJO-SNPs), and the model with the smallest number of SNPs is highlighted in blue for each trait. (i) Manc-COJO input: African-specific GWAS summary statistics from GLGC (N ~ 90,000), combined with new European-specific GWAS conducted in UK Biobank participants (N = 90,000) using pipelines consistent with the original GLGC studies. The LD reference panels consist of 6,901 unrelated European individuals and 6,901 unrelated African individuals from the UK Biobank. (ii) Sanc-COJO on multi-ancestry data input: Meta-analysis of the European-specific and African-specific GWAS summary statistics described above. The LD reference panel is a merged set of the unrelated European and African individuals from the UK Biobank (6,901 Europeans and 6,901 Africans). (iii) Sanc-COJO on single-ancestry data input: new European-specific GWAS conducted in UK Biobank participants (N = 180,000) using pipelines consistent with the original GLGC studies. The LD reference panel was a set 13,802 unrelated European and African individuals from the UK Biobank. To ensure fair comparisons across methods, we harmonised the GWAS summary statistics to have identical SNP sets and approximately equal sample sizes, and we kept identical LD reference panel sizes across methods. Overall, Manc-COJO models showed slightly lower out-of-sample prediction accuracy into European cohorts compared to single-ancestry Sanc-COJO models. However, Manc-COJO models were much smaller and consistently demonstrated much better trans-ancestry portability. Compared to the full-size analysis, the relative size difference between Manc-COJO and Sanc-COJO models decreased, suggesting that closer ancestry balance may enhance the detection of independent associations. Notably, Sanc-COJO models trained on multi-ancestry data continued to show sub-optimal out-of-sample performance across all scenarios.

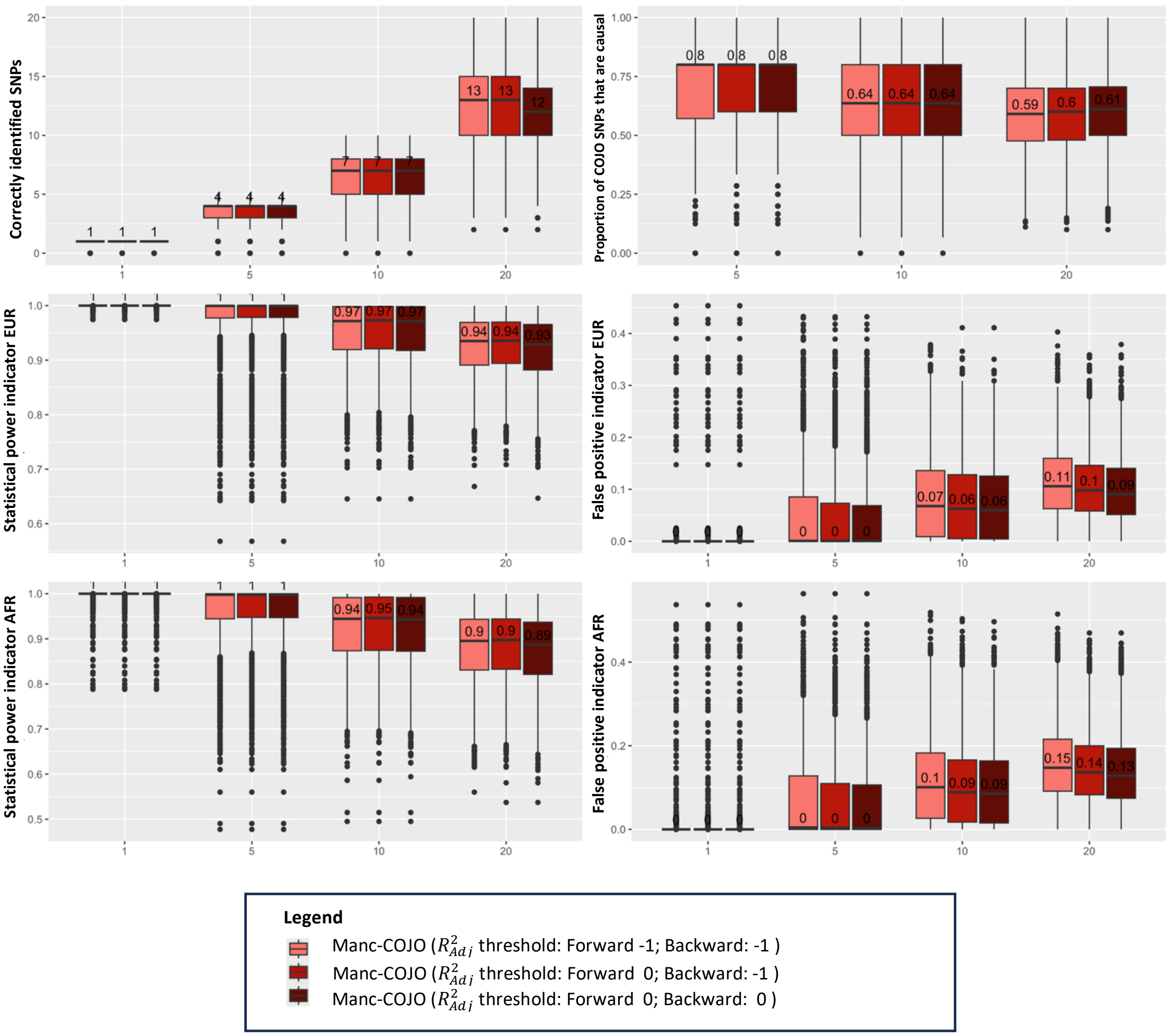

**Supplementary Figure 7. Impact of different adjusted $R^2$ threshold on Manc-COJO output in idealised situations**. The simulation settings and layout of this figure are identical to those in Figure 2. In this figure, we compare, under a well-powered scenario with a EUR:AFR ratio of 10:1, the impact of adding an additional optional variable selection algorithm based on the model's adjusted $R^2$. In general, applying a more stringent threshold reduces the false positive inclusion of non-causal SNPs in the model at the expense of reduced power to include causal SNPs.

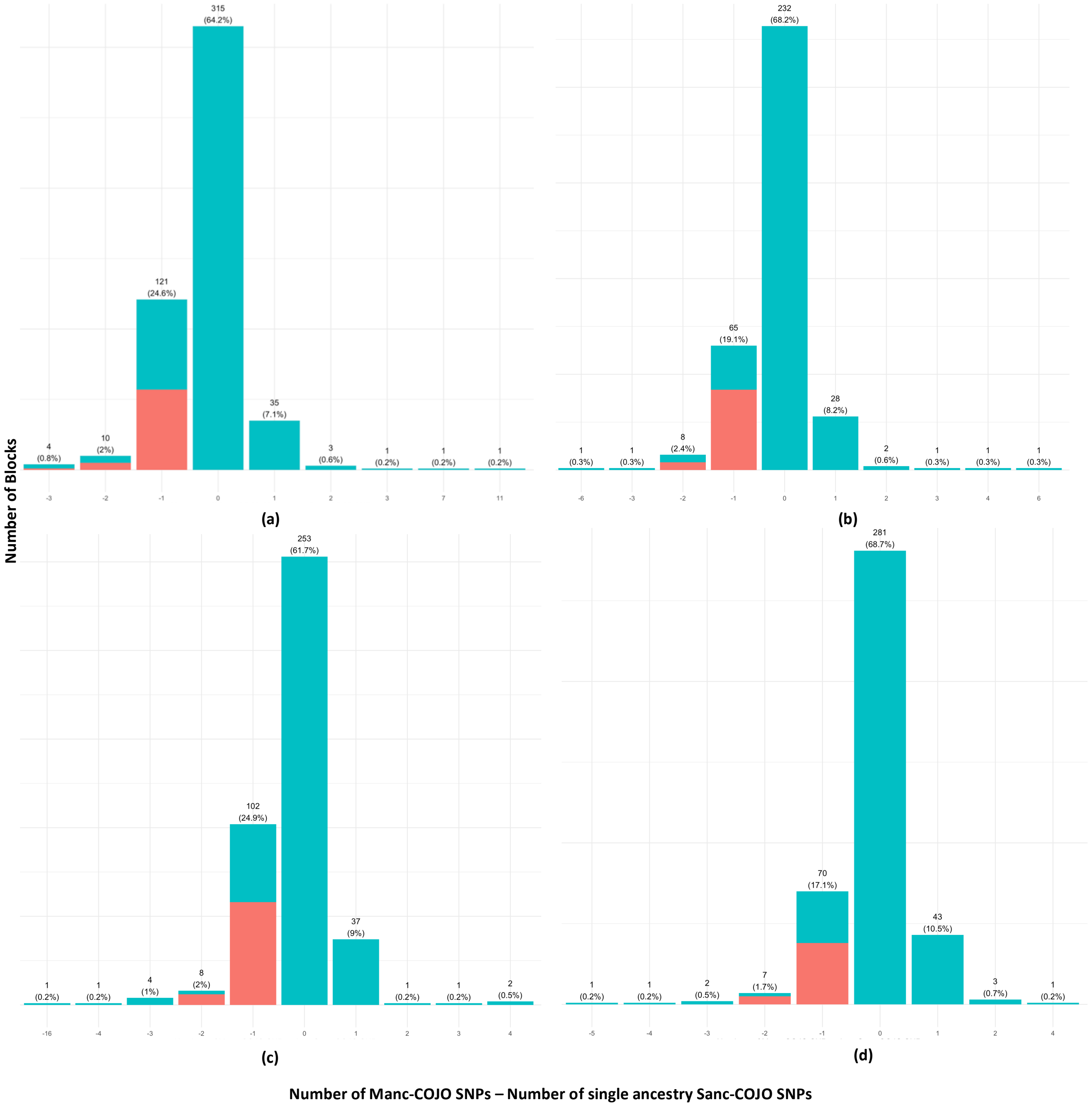

**Supplementary Figure 8. Comparison of Manc-COJO and Sanc-COJO outputs.** This figure compares the number of COJO SNPs selected by the Manc-COJO model (using GWAS input of 0.9 million EUR + 0.09 million AFR) with the single-ancestry Sanc-COJO model (using GWAS input of 0.9 million EUR + 0.09 million EUR) for (a) HDL, (b) LDL, (c) logTG, and (d) TC. COJO-selected SNPs in both models were partitioned into LD-independent blocks. The x-axis shows the difference in SNP counts between the models (number of Manc-COJO SNPs minus number of Sanc-COJO SNPs), where a positive value indicates that Manc-COJO identified more SNPs than Sanc-COJO. The y-axis represents the number of blocks. The number shown above each histogram bar is the count of blocks, with the percentage in parentheses indicating the proportion of all blocks corresponding to that x-value. The **red** sections of the histograms highlight blocks where the Manc-COJO model identified zero SNPs while the Sanc-COJO model identified more than one SNP.
